# Joint modelling of whole genome sequence data for human height via approximate message passing

**DOI:** 10.1101/2023.09.14.557703

**Authors:** Al Depope, Jakub Bajzik, Marco Mondelli, Matthew R. Robinson

## Abstract

Human height is a model for the genetic analysis of complex traits, and recent studies suggest the presence of thousands of common genetic variant associations and hundreds of low-frequency/rare variants. Here, we develop a new algorithmic paradigm based on approximate message passing, gVAMP, for identifying DNA sequence variants associated with complex traits and common diseases in large-scale whole genome sequence (WGS) data. We show that gVAMP accurately localizes associations to variants with the correct frequency and position in the DNA, outperforming existing fine-mapping methods in selecting the appropriate genetic variants within WGS data. We then apply gVAMP to jointly model the relationship of tens of millions of WGS variants with human height in hundreds of thousands of UK Biobank individuals. We identify 59 rare variants and gene burden scores alongside many hundreds of DNA regions containing common variant associations, and show that understanding the genetic basis of complex traits will require the joint analysis of hundreds of millions of variables measured on millions of people. The polygenic risk scores obtained from gVAMP have high accuracy (including a prediction accuracy of ∼ 46% for human height) and outperform current methods for downstream tasks such as mixed linear model association testing across 13 UK Biobank traits. In conclusion, gVAMP offers a scalable foundation towards a wider range of analyses in WGS data.

## Introduction

Efficient utilization of large-scale biobank data is crucial for inferring the genetic basis of disease and predicting health outcomes from DNA. By mapping associations to regions of the DNA, we seek to understand the underlying genetic architecture of phenotype. The common approach is to use single-marker or single-gene burden score regression [1–4], which gives marginal associations that do not account for linkage disequilibrium (LD) among loci. Fine-mapping methods are then employed with the aim of controlling for LD when conducting variable selection and resolving the associations to a set of likely causal variants [5].

Broadly speaking, inference for association testing [1–4], fine-mapping [5] and polygenic risk score [6, 7] tasks are performed either using Markov Chain Monte Carlo (MCMC) or variational inference (VI) approaches.

From a practical perspective, current software implementations of these algorithms limit either the number of markers or individuals. Mixed linear model association (MLMA) approaches are typically restricted to using less than one million SNPs to control for the polygenic background [1, 2], resulting in a loss of power [7] and the potential for inadequate control for fine-scale confounding factors [8]. Polygenic risk score algorithms are limited to a few million SNPs, and lose power by modeling only blocks of genetic markers [9, 10]. Likewise, fine-mapping methods are generally limited to focal segments of the DNA [5], and they are unable to fit all genome-wide DNA variants together. Thus, no existing approach can truly estimate all genetic effects jointly in large-scale whole genome sequence (WGS) data.

Here, we propose a new inference approach, gVAMP, a form of Bayesian whole genome regression, capable of fitting tens of millions of whole genome sequence (WGS) variants jointly for hundreds of thousands of individuals. We extensively benchmark gVAMP in simulations on WGS data and show it accurately localizes associations to variants with the correct frequency and position in the DNA, outperforming marginal testing and marginal testing plus fine-mapping. We then describe an application of gVAMP to WGS data and measurements of human height for hundreds of thousands of UK Biobank [11] participants. We additionally validate the application of gVAMP for polygenic risk score prediction on a number of datasets by benchmarking against the state of the art. Here, gVAMP outperforms widely used summary statistic methods such LDpred2 [9] and SBayesR [10]; its performance matches that of MCMC sampling schemes [7], but with a dramatic speed-up in time (analysing 8.4M SNPs jointly in hours as opposed to weeks); and it provides some improvement over a recently proposed individual-level variational inference approach [12]. When polygenic risk scores are used for association testing in a mixed-linear model framework, gVAMP also outperforms both the REGENIE [1] and LDAK-KVIK [12] software. In summary, our approach lays the foundations for a wider range of analyses in large WGS datasets.

### Design

Our focus is on joint modeling of all genetic variants genome-wide controlling for local and long-range LD. To do this, we consider a general form of whole-genome Bayesian variable selection regression (BVSR), common to genome-wide association studies (GWAS) [7, 10], estimating the effects vector ***β*** ∈ ℝ^*P*^ from a vector of normalized phenotype measurements ***y*** = (*y*_1_, …, *y*_*N*_) ∈ ℝ^*N*^ given by

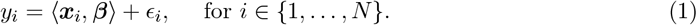

Here, ***x***_*i*_ is the row of the normalized genotype matrix ***X*** corresponding to the *i*-th individual, 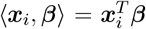 denotes the inner product, and ***ϵ*** = (*ϵ*_1_, …, *ϵ*_*N*_) is an unknown noise vector with multivariate normal distribution 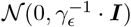 and unknown noise precision 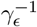. We note that the normalized genotype design matrix was used in its original form, without correcting for sample covariances, because the analysis performed in this work is targeting a genetically homogeneous population. To allow for a range of genetic effects, we select the prior on ***β*** to be of an adaptive spike-and-slab form:

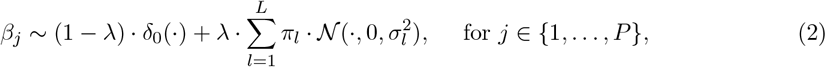

where we learn from the data: the DNA variant inclusion rate *λ* ∈ [0, 1], the unknown number of Gaussian mixtures *L*, the mixture probabilities 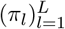 and the variances for the slab components 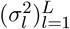.

We apply the statistical model of Equations (1) and (2) by developing a new approach for GWAS inference, dubbed *genomic Vector Approximate Message Passing* (gVAMP). Approximate Message Passing (AMP) [13–15] refers to a family of iterative algorithms with several attractive properties: *(i)* AMP allows the usage of a wide range of Bayesian priors; *(ii)* the AMP performance for high-dimensional data can be precisely characterized by a simple recursion called state evolution [16]; *(iii)* using state evolution, joint association test statistics can be obtained [17]; and *(iv)* AMP achieves Bayes-optimal performance in several settings [17–19]. However, we find that existing AMP algorithms proposed for various applications [20–23] cannot be transferred to biobank analyses as: *(i)* they are entirely infeasible at scale, requiring expensive singular value decompositions; and *(ii)* they give diverging estimates of the signal in either simulated genomic data or the UK Biobank data. To address the problem, we combine a number of principled approaches to produce an algorithm tailored to whole genome regression as described in the Method Details section of Star Methods (Algorithm 1).

gVAMP provides both *(i)* a point estimate of the posterior mean 𝔼[***β*** | ***X, y***] which can be directly used to create a polygenic risk score, and *(ii)* a statistical framework, via state evolution, to test whether each marker makes a non-trivial contribution to the outcome. Specifically, state evolution yields posterior inclusion probabilities (PIP) for each SNP (see Quantification and Statistical Analysis section of Star Methods). Finally, we learn all unknown parameters in an adaptive Bayes expectation-maximization (EM) framework [24, 25], which avoids expensive cross-validation and yields biologically informative inference of the phenotypic variance attributable to the genomic data (SNP heritability, 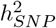) allowing for a full genome-wide characterization of the genetic architecture of human complex traits in WGS data.

## Results

### Variable selection in whole genome sequencing data

We begin by modeling the UK Biobank WGS data, curating a set of 16,854,878 WGS variants observed for 426,462 individuals of European genetic ancestry. To create this set, we remove individuals of first-degree ancestry to avoid close-familial environmental sharing, and because of cloud computing restrictions, we group highly correlated common variants with absolute correlation ≥ 0.6, keeping all rare variation within the data with minor allele frequency, MAF, ≥ 0.0001 (see Method Details section of Star Methods).

We first assess the ability of gVAMP to estimate genetic effects within this curated UK Biobank WGS dataset, by simulating 12 different scenarios on top of data from chromosomes 11-15 (3,285,117 SNPs). The 12 scenarios differ in: *(i)* the relationship of effect size and MAF; *(ii)* the ratio of common to rare causal variants; and *(iii)* the number of causal variants (500 or 1000) that contribute 7.25% to the phenotypic variance (see Method Details section of Star Methods and Supplementary Table S1 for an overview of the scenarios). Chromosomes 11-15 represent ∼ 18.8% of the total length of the DNA and if we extrapolate genome-wide, then our simulated phenotypes would have ∼ 39% phenotypic variance attributable to between approximately 2,700 and 5,300 underlying causal variants genome-wide. This gives an expected average variance contributed per DNA variant of 39%/5300 = 0.0074%, comparable to that of human height of 70%/10000 = 0.007% suggested by recent results [26]. These settings are chosen to give adequate power to select variables and to enable clear comparisons of fine-mapping methods. We note also that in Supplementary Note 1 we conduct extensive benchmarks of gVAMP in imputed SNP data for traits of highly polygenic architecture with different effect size distributions, different relationships of marker frequency and effect size, different numbers of causal variants and different variances attributable to the SNPs, across 12 additional settings each with 4 sets of data, for a total of hundreds of models trained with sample size ∼ 400k and up to > 8M SNPs (see Method Details section of Star Methods and Supplementary Note 1).

We analyze the WGS data with either: *(i)* our proposed gVAMP; or *(ii)* the FINEMAP [27] and SuSiE variable selection models [5], run on 3MB regions surrounding each significant variant discovered via GWAS marginal testing one-SNP-at-a-time, following standard practice. With access to 2240 virtual CPU and just under 9TB (35 parallel instances of 256GB and 64 CPU), running gVAMP for all replicates of the 12 scenarios requires 5 days, with FINEMAP requiring 8 days. However, running SuSiE required 30 days to complete 4 scenarios (with standard LD and eigen-value calculation in LDstore2, see Method Details section of Star Methods), and given the consistently comparable performance to FINEMAP, we restrict our comparisons with SuSiE to only the settings presented in Figure 1. We first calculate the true positive rate (TPR = number of identified variants that are causal / number of causal variants) and the false discovery rate (FDR = number of identified variants that are not causal / number of identified variants) at a given threshold. gVAMP shows improved power (TPR) for identifying the correct underlying causal variants and a well controlled FDR (FDR ≤ 5% corresponding to a selection of variants with posterior inclusion probability, PIP, ≥ 95%), as compared to both the methods of FINEMAP and SuSiE across most scenarios (Figure 1a and S1 to S3). FINEMAP shows a well-controlled FDR in only five of the 12 settings (settings *f, h, j, k* and *l*, in Figures S1 to S3), where half of the simulated causal variants are common and the power relationship of minor allele frequency and effect size is weaker. In contrast, we find only two settings where the FDR of gVAMP rises slightly above 5% (settings *i* and *k* in Figures S1 to S3). In these two settings, the relationship between effect size and minor allele frequency is weakly negative and variants are distributed randomly across the WGS data. As a result, most causal variants are rare and each makes a very small contribution to the phenotypic variance, making them difficult to detect. Consequently, these are the lowest powered settings, where the TPR of both gVAMP and FINEMAP are the smallest and the fewest variants overall are recovered (Figure S1). gVAMP outperforms FINEMAP in terms of TPR in all but one scenario, namely the challenging setting *k*, where the TPR of both methods is low (Figures S1 and S3). SuSiE performs similarly to FINEMAP as it is typically unable to control the FDR (Figure 1a). SuSiE has higher TPR than FINEMAP, but it is still consistently outperformed by gVAMP, setting *f* being the only one where all methods perform similarly (Figure 1a).

**Figure 1.**
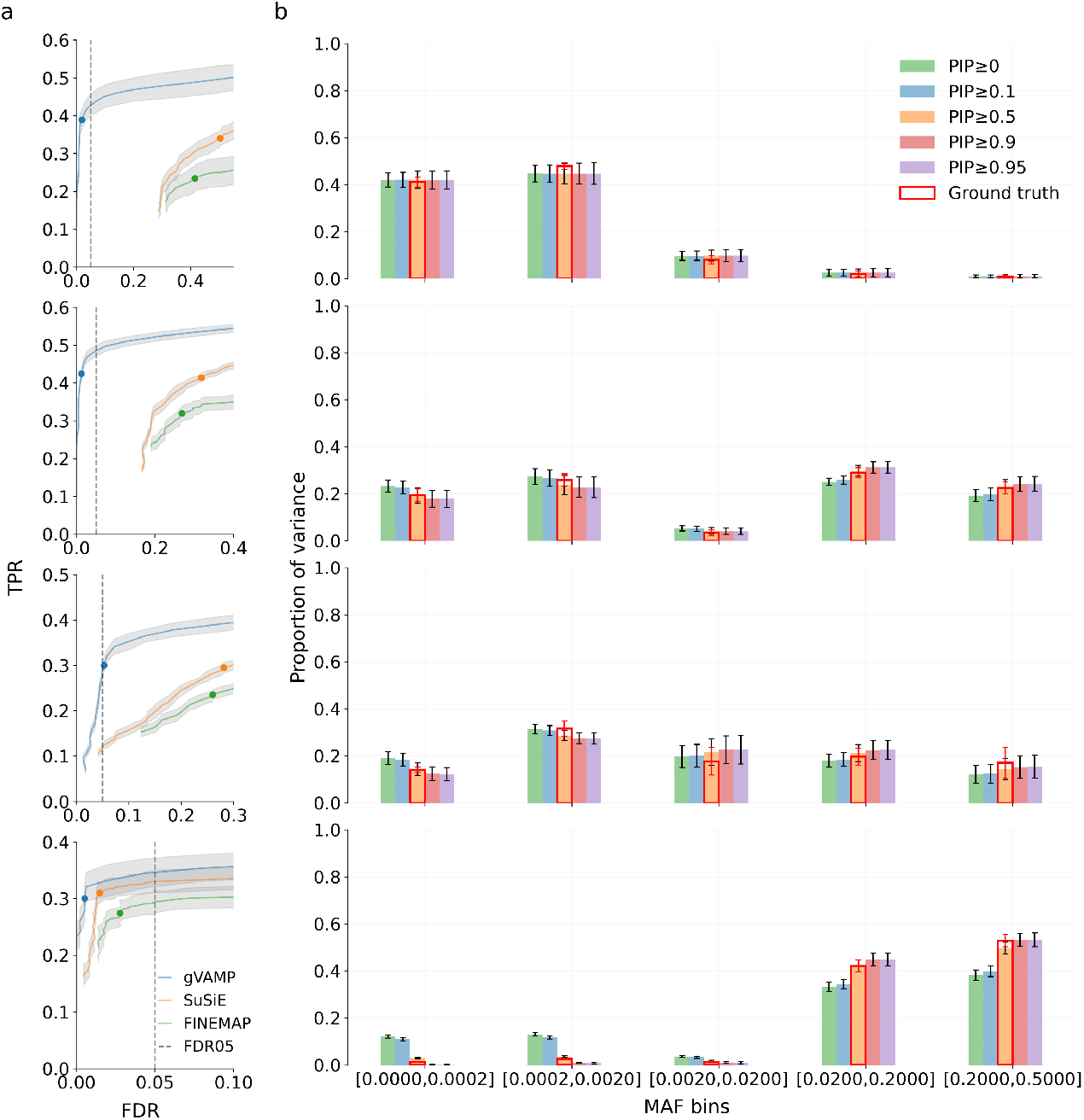
Simulation study of variable selection performance using whole genome sequence (WGS) data of chromosomes 11 to 15 in the UK Biobank. (a) Comparison of the true positive rate (TPR, *y*-axis) versus the false discovery rate (FDR, *x*-axis) between gVAMP (blue) and marginal GWAS association testing plus fine-mapping implemented in the FINEMAP (green) and SuSiE (orange) algorithms across four simulation scenarios. The rows of both (a) and (b) correspond to settings *a, b, e*, and *f*, which are described in the Method Details section of Star Methods, Supplementary Table S1 and Figures S1 to S3, alongside additional simulation settings. For gVAMP, variable selection is conducted from posterior inclusion probability (PIP) values obtained from a single model run genome-wide. For FINEMAP and SuSiE, we use a Bonferroni corrected p-value significance threshold for marginal GWAS testing and then, having identified associated regions, we use the algorithms to obtain the PIP values to conduct variable selection. Error bands give the standard deviation across 5 simulation replicates. The blue, green and orange dots give the TPR and FDR at PIP ≥ 0.95, which should correspond to an FDR of ≤ 0.05 if the model correctly conducts variable selection based on PIP values. (b) The proportion of variance attributable to DNA loci of different frequency. We plot the proportion of the sum of the squared regression coefficients attributable to variants of different frequency as estimated from gVAMP across different posterior inclusion probability (PIP) thresholds. The ground truth (red frame) is calculated as the sum of squared true effects across different frequency groups. The variance attributable to SNPs of different frequency matches the ground truth when selecting variants at posterior inclusion probability ≥ 0.5. Error bars give the standard deviation across simulation replicates.

The advantage of the gVAMP framework is confirmed in additional simulation comparisons. In particular, we find that infinitesimal versions of SuSiE and FINEMAP [28] that have been proposed in the literature also show only slight improvements on the standard versions in all WGS settings we explored (Figure S4). Additionally, we find that variants selected by gVAMP at posterior inclusion probability ≥ 0.5, have a frequency distribution that reflects the simulated ground truth across scenarios (Figure 1b and S5). Furthermore, we validate the part of our analysis which combines WGS variants and WES burden scores by performing an additional set of simulations on top of WGS variants from chromosomes 11 to 15 and WES burden scores corresponding to that region of the DNA. We simulated signals with a total heritability of *h*^2^ = 7.25% by allocating 75% of 500 causal markers to WGS variants and 25% to WES burden scores, with WES burden scores exhibiting effect sizes with variance three-fold larger than those of individual WGS variants. This means that both groups should explain half the variance in settings where the variance of effect sizes is independent of the minor allele frequency. We find that we can correctly recover the percentage of variance recovered by each of the groups (Figure S6). Finally, when we simulate causal effects in both coding and non-coding regions, with 0%, 50%, or 100% of the causal variants located in coding (exonic) regions, we find that as the percentage of causal variants allocated to coding regions increases so does the enrichment of the effects (Figure S7). Taken all together, these findings suggest that existing methods fall short of the performance required for variable selection in WGS data, while gVAMP both reliably localizes the signal to the correct genomic regions and it selects a variant of appropriate frequency.

### The genetic architecture of human height in WGS data

We then apply gVAMP to model the genetic architecture of human height in our 16,854,878 WGS variant dataset. We find significant amounts of common genetic variation detected and a considerable contribution of rare genetic variants (Figure 2). Generally, the regions identified are replicated in the most recent GWAS study of human height [26] within 100-500kb (Figure 2a). We find that for variants of frequency < 0.05 identified in the WGS analysis, variants discovered by Yengo *et al*. [26] of strongest correlation and most similar MAF, significant at *p* ≤ 5 · 10^−8^ within a 500kb region, are more common in frequency (Figure 2b). Variants of frequency < 0.005 discovered by gVAMP are likely not captured at all by Yengo *et al*. because these variants have lower average marginal Chi-squared test statistic values than the other variants we identify (Figure 2c) and Yengo *et al*. set a MAF threshold of 1% for their meta-analysis. We estimate the maximum correlation within the UK Biobank WGS data of each gVAMP discovery with Yengo *et al*. SNPs significant at *p* ≤ 5 · 10^−8^ within a 1 Mb window. We find that none of the variants of frequency < 0.005 identified by gVAMP have a strong correlation with those identified by Yengo *et al*. (Figure S8). Thus, very rare gVAMP-identified variants are likely independent of the colocalizing common variants from Yengo *et al*., being simply a randomly selected set with higher frequency (Figure 2b). In contrast, all gVAMP identified variants of frequency ≥ 0.05 are highly correlated to variants identified by Yengo *et al*. (Figure S8) with similar frequencies (Figure 2b). Thus, consistent with the literature [29], common variant-tagged rare variant associations likely make up a small fraction of common-variant associations in Yengo *et al*., and taken together, our results suggest that discovered WGS regions are broadly the same as those previously identified.

**Figure 2.**
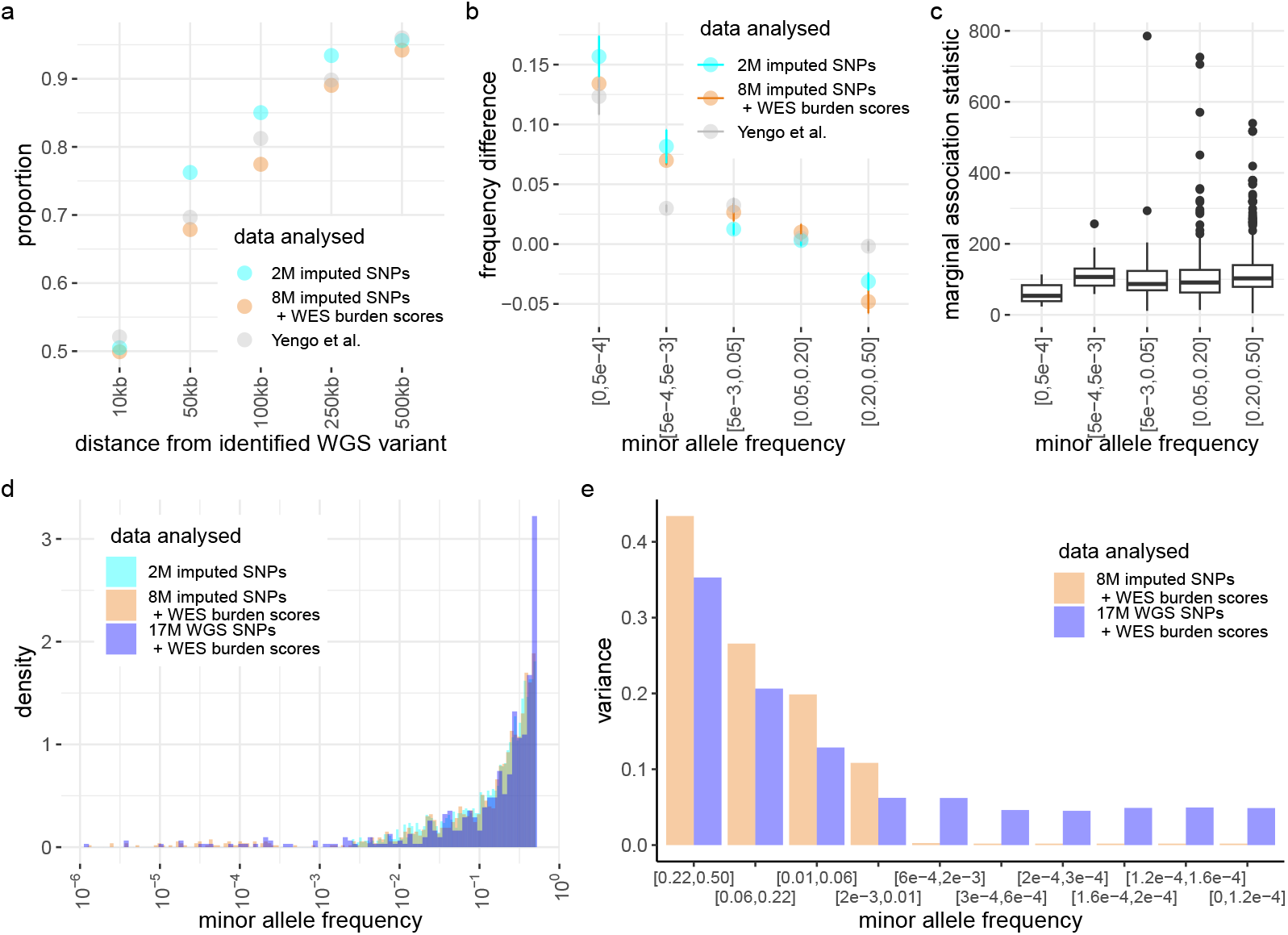
The genetic architecture of human height inferred by gVAMP from 16,854,878 whole genome sequence variants in the UK Biobank. *(a)* For each whole genome sequence (WGS) variant discovered at PIP ≥ 95%, we determine whether we also identify a variant within a given number of base pairs (distance, *x*-axis) from: *(i)* the latest height GWAS study by Yengo *et al*. [26], selecting variants at significance *p* ≤ 5 · 10^−8^ from the marginal summary statistics for European individuals excluding the UK Biobank; or *(ii)* an analysis of imputed SNP data with gVAMP, selecting variants with PIP ≥ 95%. The proportion overlap (*y*-axis) is calculated as the proportion of the identified WGS variants that are discovered within a certain base pair distance. *(b)* For each WGS variant discovered at PIP ≥ 95%, we determine: *(i)* the variant discovered by Yengo *et al*. of most similar MAF significant at *p* ≤ 5 · 10^−8^ within a 500kb region; or *(ii)* the imputed variant of most similar minor allele frequency (MAF) discovered at PIP ≥ 95% within a 500kb region. We then calculate the mean difference in MAF of the imputed/GWAS variant and the corresponding WGS variant, with error bars showing twice the standard error. *(c)* For each WGS variant discovered at PIP ≥ 95%, we determine the maximum marginal association test statistic of any variant with absolute correlation ≥0.6. Boxplots of these maximum Chi-squared test statistic values are then given for different MAF groups, showing a drop in average power to detect rare variants. *(d)* Density plot of the MAF distribution of all discovered variants at PIP ≥ 95% for 2,174,071 imputed SNPs (“2M imputed SNPs”), 8,430,446 imputed SNPs fit jointly alongside 17,852 whole exome sequencing (WES) gene burden scores (“8M imputed SNPs + WES burden scores”), and 16,854,878 WGS SNPs fit jointly alongside 17,852 WES gene burden scores (“17M imputed SNPs + WES burden scores”). Irrespective of the data analyzed, the frequency distributions are similar, with the exception that the WGS analysis identifies a greater number of variants (both SNPs and gene burden scores) at lower frequencies. *(e)* We calculate the proportional contribution to phenotypic variance across different MAF groups for 8,430,446 imputed SNPs fit jointly alongside 17,852 whole exome sequencing (WES) gene burden scores (“8M imputed SNPs + WES burden scores”), and 16,854,878 WGS SNPs fit jointly alongside 17,852 WES gene burden scores (“17M imputed SNPs + WES burden scores”). A larger proportion of variance is attributable to rare variation at the expense of more common variants in WGS data.

19 out of the 21 genome-wide significant gene burden scores are for genes that have previous GWAS height associations linked to them in Open Targets, but our analysis suggests that the effect is attributable to a rare protein coding variant rather than the common variants suggested by current genome-wide association studies. This includes insulin-like growth factor 2 genes IGF1R and IGF2BP2, and known height genes ACAN, ADAMTS10 and ADAMTS17, CRISPLD1, DDR2, FLNB, HMG20B, LTBP2, LCORL, NF1, and NPR2. Novel genes are COL27A1 which encodes a collagen type XXVII alpha 1 chain, and HECTD1. The top genes, where gVAMP attributes ≥ 0.04% of height variation in addition to those listed above are EFEMP1, ZBTB38, and ZFAT. We find that many genome-wide significant height-associated rare variants discovered in the WGS data were not found in previous UK Biobank analyses in Open Targets including: rs116467226, an intronic variant by TPRG1; rs766919361, an intronic variant by FGF18; rs141168133, an intergenic variant 19kb from ID4; rs150556786, an upstream gene variant for GRM4; rs574843917, a non-coding transcript exon variant in GPR21; rs532230290, an intergenic variant 42kb from SCYL1; rs543038394, intronic in OVOL1; rs1247942912, a non-coding transcript exon variant in AC024257.3; rs577630729, a regulatory region variant for ISG20; and rs140846043, a non-coding transcript exon variant of MIRLET7BHG. 10 out of our top 38 height-associated WGS variants of ≤ 1% MAF were not previously discovered, and only become height-associated when conditioning on the entire polygenic background captured by WGS data within our analysis.

Generally, marginal estimates made in WGS data for height have similar joint effect sizes when included within the gVAMP model (Figure S9). We find high replication for variants directly observed within the All of Us study (Figure S10) when we compare the jointly fitted gVAMP estimates to simple marginal OLS estimates made within All of Us, controlling for sex and 16 available PCs in (Figure S10). The slope of the regression coefficients is 0.8471, which is a previously used metric of replication [30].

We then compare our gVAMP WGS analysis to gVAMP run on two curated sets of 8,430,446 and 2,174,071 imputed SNPs for the same individuals (see Method Details section of Star Methods) but with an elevated MAF threshold, to contrast the findings obtained when setting different MAF thresholds, and to explore the benefit of using WGS data over imputed SNP data. gVAMP estimates the proportion of phenotypic variance in human height attributable to WGS data as 0.652, a value that is comparable to 0.63 for imputed SNP data, to the previously published REML estimate in a different WGS dataset [31] and to family-based estimates [32]. This is expected as the imputed data is a subset of the WGS data and very low frequency variants are expected to make little contribution to the population-level variance, so that the addition of large numbers of rare variants will have little impact, especially if they capture signal that was previously weakly tagged by more common variation.

gVAMP identifies 2,070 sets of variants when we apply the model to a set of 2,174,071 imputed SNPs at PIP ≥ 95%, which is consistent with findings from mixed-linear model association testing described in the next section within imputed data (Table 2). We note that in the WGS data however, gVAMP identifies fewer genome-wide significant effects, with only 501 found at PIP ≥ 95%. There are two potential explanations for this. The first is that some of the height associations are re-allocated to DNA variants of lower minor allele frequency, and we then have lower power to detected them above a PIP ≥ 95% threshold. The second explanation is that two variants can have very low correlation with each other in the imputed data, while still both being moderately correlated to a third SNP. If the third SNP is not in the imputed panel but is height associated, this would result in two quasi-independent associations in the imputed data, but only one in the WGS data. We find that our 501 findings are within 50kb of 1,042 of the 2,070 sets of variants identified by gVAMP in the 2,174,071 imputed SNP data and are within 500kb of 1,989 of the 2,070 variant sets. Thus, the WGS gVAMP findings are often neighbored by multiple variants discovered by gVAMP in the imputed SNP data. The WGS analysis identifies a greater proportion of variants (both SNPs and gene burden scores) at lower frequencies (Figure 2d) and finds a larger proportion of variance attributable to rare variation at the expense of slightly less variation attributable to common variants (Figure 2e).

We show that rare effects are larger than the effects of common variants when both are fit jointly (Figure 3a). We estimate the effects of 17,852 WES gene burden scores when fit alongside the WGS data, finding again that conditioning on rare variation enhances joint effect sizes (Figure 3a and Figure 3b). We find a general concordance of the estimated effect sizes for most but not all WES gene burden scores (Figure 3b). WES gene burden score effects can differ in the WGS analysis by being either: *(i)* more strongly associated because of improved conditioning on the genetic background, or *(ii)* attenuated because the variants that comprise the burden score are also included as variants in the analysis. We sum up the joint effects to determine the variation in height attributable to each gene, and we find a general concordance across markers annotated to each of the 17,852 genes, conditional on all other markers, across the imputed SNP and WGS analysis (Figure 3c). We see little relationship between the variance attributable to the burden score of a given gene and the SNPs annotated to the gene (Figure S11), with some notable exceptions of: ACAN, ADAMTS17, ADAMTS10, and LCORL. Additionally, across annotations, we find that the phenotypic variance attributable to different DNA regions is higher for intergenic and intronic variants (Figure 3d). However, when adjusting for the number of SNPs contained by each category by using the average effect size of the group relative to the average effect size genome-wide, we find that exonic variants that are nonsynonymous, splicing and stop-gain have the largest average effects of the WGS variants included within our model (Figure 3d). Finally, across annotations, we find similar patterns across both WGS and imputed SNP analyses (Figure 3d). Taken together, we find strong concordance across WGS and imputed data analyses performed on the same people, but we highlight MAF cut-offs influence the variants to which effects are assigned and thus where phenotypic variance is allocated. Therefore, we expect WGS data to provide a more accurate characterization of the genetic architecture of human traits, but very large sample sizes are required to incorporate more rare variation so that effects can be appropriately assigned.

**Figure 3.**
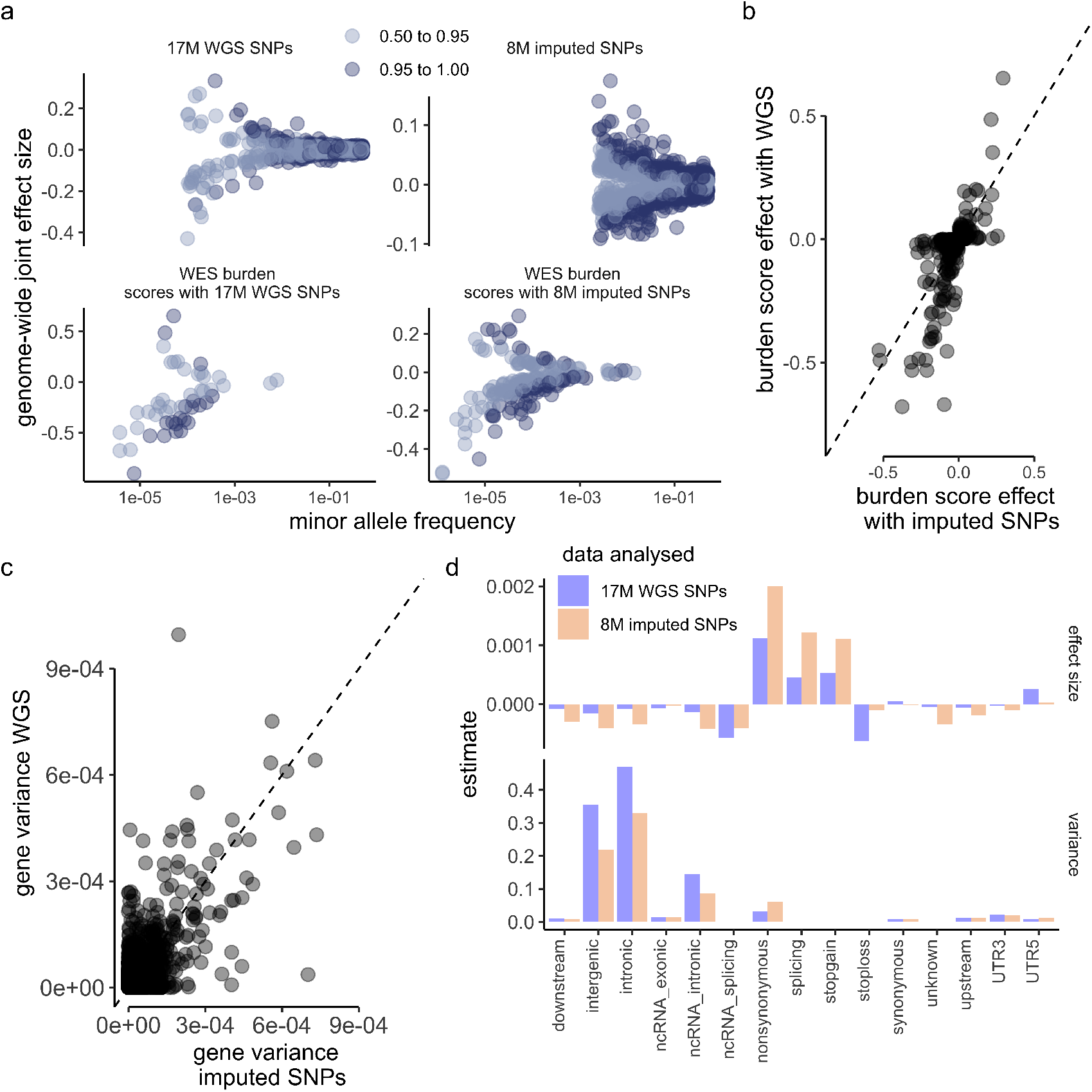
The relationship between effect size and minor allele frequency, burden score effects and the genetic architecture of human height inferred from 16,854,878 whole genome sequence (WGS) variants in the UK Biobank. *(a)* For different posterior inclusion probability thresholds (0.5 to 0.95 and ≥ 0.95), we plot the relationship between joint effect size and minor allele frequency (MAF) for 16,854,878 WGS variants (“17M WGS SNPs”), 8,430,446 imputed SNP variants (“8M imputed SNPs”) and 17,852 whole exome sequence (WES) gene burden scores fit alongside either the WGS or imputed variants. *(b)* Across 17,852 WES gene burden scores, we find general concordance of the estimated effect sizes when fit alongside either imputed or WGS data, for most but not all genes, with squared correlation 0.532 and regression slope 0.993 (0.007 SE). The dashed line gives *y* = *x. (c)* Likewise, we also find general concordance of the phenotypic variance attributable to markers annotated to each of the 17,852 genes, when fit conditional on either imputed or WGS variants, with squared correlation 0.494 and a regression slope of 0.698 (0.004 SE). The dashed line gives *y* = *x. (d)* Finally, we show similar patterns of enrichment when annotating markers to functional annotations in either the average effect size relative to the genome-wide average effect size (labeled “effect size”) or the proportion of variance attributable to each group (labeled “variance”), from joint estimation of either imputed or WGS data.

### Creating genetic predictors via gVAMP

We validate and benchmark gVAMP against state-of-the-art polygenic risk score prediction approaches in a number of alternative datasets. Applying our model to 13 traits in the UK Biobank imputed data (training data sample size and trait codes given in Table S2), we compare the prediction accuracy of gVAMP to the widely used summary statistic methods LDpred2 [9] and SBayesR [10], and to the individual-level methods GMRM [7] and LDAK [12]. gVAMP matches or outperforms all methods for most phenotypes (Figure 4a and Table 1 with the best prediction accuracy highlighted in bold). Specifically, for human height (HT), our prediction accuracy is the highest and individual-level approaches clearly outperform summary statistic ones: we obtain an accuracy of 45.7%, which is a 97.8% relative increase over LDpred2 and a 26.2% relative increase over SBayesR (Figure 4a and Table 1). We also note that our polygenic risk score has higher accuracy than that obtained from SBayesR using the latest height GWAS study of 3.5M people of 44.7% [26], despite our sample size of only 414,055.

**Table 1.**
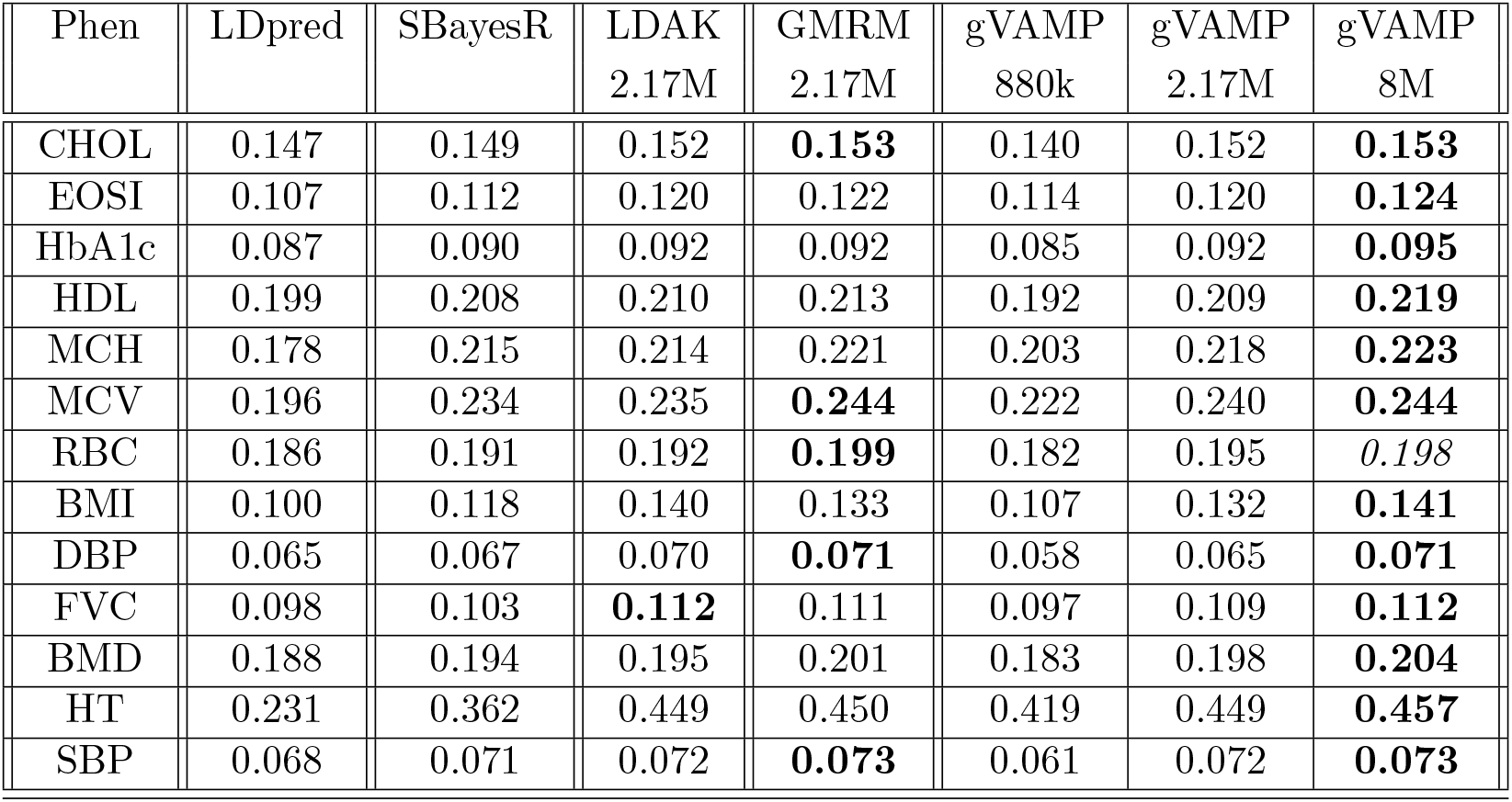
Polygenic risk score prediction accuracy *R*^2^ for 13 different traits from statistical models trained in the UK Biobank data and tested in a UK Biobank hold-out set. Training data sample size and trait codes are given in Table S2 for each trait. The sample size of the hold-out test set is 15, 000 for all phenotypes. LDpred2 and SBayesR give estimates obtained from the LDpred2 and SBayesR software respectively, using summary statistic data of 8,430,446 SNPs obtained from the REGENIE software. LDAK gives the estimates obtained by the LDAK software using the elastic net predictor and individual-level data of 2,174,071 SNP markers. GMRM denotes estimates obtained from a Bayesian mixture model at 2,174,071 SNP markers (“GMRM 2M”). gVAMP denotes estimates obtained from the adaptive EM Bayesian mixture model within a vector approximate message passing (VAMP) framework presented in this work, using either 887,060 (“gVAMP 880k”), 2,174,071 (“gVAMP 2M”), or 8,430,446 SNP markers (“gVAMP 8M”).

**Figure 4.**
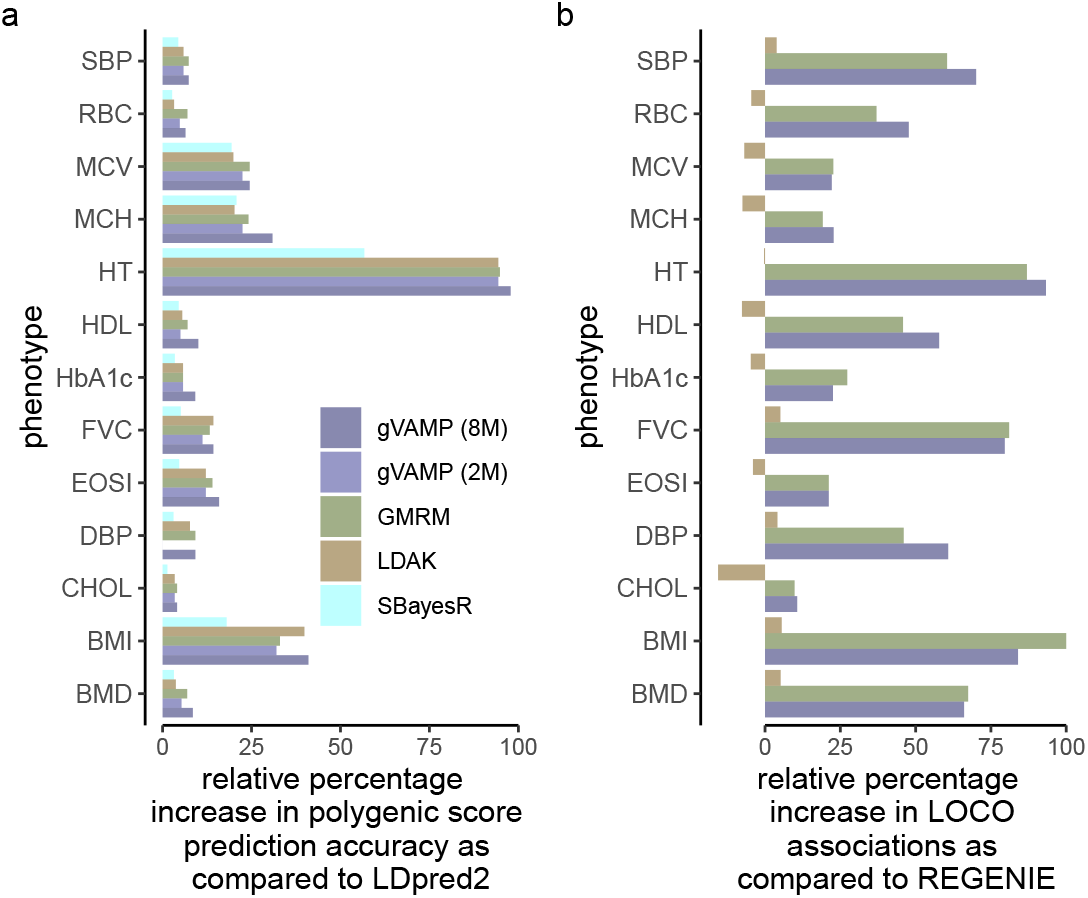
Validating gVAMP through polygenic risk score accuracy and association testing benchmarks in the UK Biobank within imputed SNP data. *(a)* Relative prediction accuracy of gVAMP in a hold-out set of the UK Biobank across 13 traits as compared to other approaches. *(b)* Relative number of leave-one-chromosome-out (LOCO) associations of gVAMP across 13 UK Biobank traits as compared to other approaches at 8,430,446 markers.

The polygenic risk score accuracy presented may represent the limit of what is achievable for these traits at the UK Biobank sample size. While modeling 2.17M SNPs improves accuracy over modeling 880,000 variants, we find rather small differences when going from 2.17M to 8M. This is because the 2.17M data set is an LD-grouped subset of the 8M data, and the addition of highly correlated variants captures little additional phenotypic variance. gVAMP estimates the 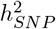 of each of the 13 traits at an average of 3.4% less than GMRM when using 2,174,071 SNP markers, but at an average of only 3.9% greater than GMRM when using 8,430,446 SNP markers. This implies that only slightly more of the phenotypic variance is captured by the SNPs when the full imputed SNP data are used (Figure S12). In summary, gVAMP (an adaptive mixture) performs similarly to LDAK (an elastic net) and GMRM (an MCMC sampling algorithm), improving over both when analyzing the full set of 8,430,446 imputed SNPs (Table 1 and Figure 4a).

Next, we compare gVAMP to REGENIE, GMRM and LDAK-KVIK for association testing of the 13 traits in the UK Biobank imputed data. gVAMP performs similarly to mixed-linear model association leave-one-chromosome-out (MLMA-LOCO) testing conducted using a predictor from GMRM, with the use of the full 8,430,446 imputed SNP markers improving performance (Table 2 and Figure 4b). REGENIE yields far fewer associations than either GMRM or gVAMP for all traits (Table 2 and Figure 4b), with LDAK-KVIK being equivalent to REGENIE. We additionally verify that placing the predictors obtained by gVAMP into the second step of the LDAK-KVIK algorithm, also yields improved performance as compared to the LDAK model (Table S3). Finally, using gVAMP to conduct variable selection, we find that hundreds of the LOCO marker associations for each trait can be localized (Table 2), with the obvious caveat that these results are for imputed SNP data and re-analysis of WGS data may yield different results as we show above for height. Using height as an example, we find that 3,367 of the 5,242 MLMA-LOCO associations can be localized directly to the 2070 imputed variant sets discovered by gVAMP via the PIP testing within 50kb. Thus, the gVAMP findings obtained by performing variable selection via the state evolution framework are often neighbored by multiple variants discovered by MLMA-LOCO in the imputed SNP data.

**Table 2.** Genome-wide significant associations for 13 UK Biobank traits from the software GMRM, gVAMP, LDAK-KVIK and REGENIE at 8,430,446 genetic variants from either mixed linear model association leave-one-chromosome-out (MLMA-LOCO) testing or gVAMP variable selection. “REGENIE 880k LOCO” denotes results obtained from MLMA-LOCO testing using the REGENIE software, with 882,727 SNP markers used for step 1 and 8,430,446 markers used for the LOCO testing of step 2. “GMRM 2M LOCO” refers to MLMA-LOCO testing at 8,430,446 SNPs, using a Bayesian MCMC mixture model in step 1, with 2,174,071 SNP markers. “LDAK 2M LOCO” refers to MLMA-LOCO testing at 8,430,446 SNPs, using the LDAK-KVIK software in step 1, with 2,174,071 SNP markers. “gVAMP 2M LOCO” refers to MLMA-LOCO testing at 8,430,446 SNPs, using the framework presented here, where in step 1 2,174,071 markers are used and the second step is identical to that of REGENIE. We then extend that to using all 8,430,446 SNP markers (“gVAMP 8M LOCO”) in step 1. We also present gVAMP posterior inclusion probability (PIP) testing when applying a correlation threshold on the markers prior to analysis of 0.9 to the SNP markers (2,174,071 markers). For LOCO testing, the values give the number of genome-wide significant linkage disequilibrium independent associations selected based upon a standard *p*-value threshold of less than 5 · 10^−8^ and *R*^2^ between SNPs in a 1 Mb genomic segment of less than 1%. For the PIP testing, values give the number of genome-wide significant associations selected based upon a ≥ 0.95 threshold.

| Trait | REGENIE<br>880k LOCO | GMRM<br>2M LOCO | LDAK<br>2M LOCO | gVAMP<br>2M LOCO | gVAMP<br>8M LOCO | gVAMP<br>PIP |
| --- | --- | --- | --- | --- | --- | --- |
| CHOL | 571 | 627 | 482 | 603 | **632** | 381 |
| EOSI | 572 | **693** | 549 | 630 | **693** | 527 |
| HbA1c | 337 | **429** | 321 | 385 | *413* | 227 |
| HDL | 692 | 1009 | 639 | 812 | **1092** | 536 |
| MCH | 746 | 889 | 690 | 810 | **916** | 691 |
| MCV | 970 | **1190** | 903 | 1062 | *1185* | 898 |
| RBC | 897 | 1229 | 856 | 1079 | **1325** | 651 |
| BMI | 688 | **1376** | 726 | 1175 | *1266* | 261 |
| DBP | 291 | 425 | 303 | 419 | **468** | 315 |
| FVC | 549 | **994** | 577 | 721 | *986* | 318 |
| BMD | 522 | **874** | 549 | 668 | *867* | 491 |
| HT | 2712 | 5070 | 2703 | 4452 | **5242** | 2070 |
| SBP | 311 | 499 | 323 | 388 | **529** | 245 |

The increase in MLMA-LOCO findings for gVAMP over LDAK and REGENIE within the UK Biobank is consistent with the extensive simulation study results presented in Supplementary Note 1. There, we show that MLMA-LOCO testing power scales with the out-of-sample prediction accuracy only for gVAMP and not for the other methods (Figures S13 and S14). Including more predictors in gVAMP to generate the polygenic risk scores used in the first stage of MLMA-LOCO generally improves out-of-sample prediction accuracy (Figure S13) and association testing power, whilst maintaining controlled type-I error (Figures S14, S15, S16). Additional simulations varying the number of markers included in the model lead to similar findings (Figures S17, S18, and S19). We also find that in practice our framework has similar performance to an MCMC approach, GMRM [7], that has a similar mixture prior (Figure S20).

In terms of compute time and resource use, gVAMP completes in less than 1/2 of the time given the same data and compute resources as compared to a single trait analysis with REGENIE (Figure S21a), it is dramatically faster than GMRM (12.5× speed-up, Figure S20c), and it is marginally faster than LDAK when run on our simulated data (Figure S21a). Across the 13 UK Biobank traits, gVAMP is faster than LDAK-KVIK, but it requires more RAM (roughly the plink file size on disk, Figure S21b). If run on cloud computing at the current DNAnexus rates, step 1 of REGENIE can accommodate all 13 UK Biobank traits together, requiring 75.6 hours with 30 threads, 15GB RAM and 421GB disk space, which costs 75 GBP for the whole analysis, or 5.77 GBP per trait. LDAK-KVIK analysis of 13 UK Biobank traits with 2.17M SNPs using 15 threads, requires up to 6 GB of RAM and 210 GB of storage, with an average runtime across the 13 traits of 26h, giving an average cost of 11.47 GBP per trait. Increasing the CPU given to LDAK to 60 as in Figure S21b reduces the average run time to 14.5h, but raises costs to 28.57 GBP per trait. GMRM analysis of 2.17M SNP markers requires 82 hours using 60 threads and 210GB RAM and disk space, which costs 163.59 GBP per trait. In comparison, gVAMP analysis of 2.17M SNP markers requires 16 hours on average with 32 threads and 210GB RAM and disk space, which costs 20.59 GBP per trait. Increasing the CPU to 60 as in Figure S21b reduces the average run time to 11h, but slightly raises the cost to 21.89 GBP per trait.

## Discussion

Here, we develop an adaptive form of Bayesian variable selection regression capable of fitting tens of millions of WGS variants jointly for hundreds of thousands of individuals. For human height, a joint analysis of a set of 17M WGS variants, localizes associations to numerous rare variants and gene burden scores as well as hundreds of regions containing common variation. gVAMP provides a point estimate from the posterior distribution and, as such, its performance is expected to match that of standard MCMC Gibbs sampling algorithms. We show that this is the case in imputed SNP data: gVAMP and GMRM have comparable out-of-sample prediction accuracy, comparable estimates of the proportion of variance attributable to the DNA, comparable number of MLMA-LOCO findings and comparable performance in our simulation study. However, unlike Gibbs sampling algorithms which cannot scale to current data sizes, gVAMP exhibits a remarkable speed-up which allows high-dimensional biobank data with full WGS observations to be analyzed. Taken together, gVAMP offers a flexible framework suitable to *(i)* conduct variable selection in WGS data, where SuSiE, FINEMAP and their variants are miscalibrated, and *(ii)* create polygenic risk scores with high accuracy that can also be used for downstream tasks such as MLMA-LOCO testing, with improved performance over LDAK-KVIK and REGENIE.

### Limitations of the Study

There are a number of remaining limitations and possible ways to expand and improve the analyses presented here. Our results suggest that the prioritization of relevant regions, genes and gene sets is feasible at current sample sizes, but the detection of specific height-associated mutations is expected to be a challenging task. There are a very large number of rare variants within the human population that are missing from our analysis of 16,854,878 WGS variants. While conducting variable selection genome-wide for these will require millions of WGS samples, it may be beneficial to include as many of these rare variants as possible. Our algorithm requires random-access memory (RAM) that is the same as the plink data file on disk, and although we evidence that this is not prohibitive to running the algorithm on the UK Biobank cloud computing, two alternative solutions could be to: *(i)* apply gVAMP region-by-region, or *(ii)* make slight modifications in the implementation, so that gVAMP streams data, at the cost of slightly increased run time. Our ongoing work focuses on exploring these options.

A further limitation is that we have exclusively tested and analyzed individuals of European ancestry. Combining data of different ancestries, or utilising admixed populations is of great importance [33], and previous work suggests that better modeling within a single large biobank can facilitate improved association testing in other global biobanks [34]. However, tackling this problem requires the development of a modeling framework that explicitly allows for differences in MAF, LD and effect size across the human population, which we leave to future work.

Additionally, we also removed first-degree relatives from all of our analyses to avoid confounding of shared environmental effects. While some relatedness still exists even after removing first-degree relatives, we show in simulations that gVAMP performs well in both WGS and imputed data when planting a signal on real observed genotype data. Our motivation is that genotype-phenotype data of close relatives is often utilized for follow-up studies and there are unique models (to which our framework could be adapted) employed for such tasks. Additionally, removing first-degree relatives loses 39,592 individuals from the UK Biobank, which does not alter the sample size in a significant way. We note that methods such as fastGWA [4] or the equivalent implementation in LDAK, could be used where a first step calculates the parameters needed to control for environmental confounding before the model is then run. Furthermore, our method currently requires LD pruning of highly correlated common variants to improve numerical stability and convergence. While this limits the ability of gVAMP to fine-map regions with extremely high correlations, we note that when rare variants are included in the analysis, we find that gVAMP demonstrates superior performance compared to FINEMAP, SuSiE, and their respective variants.

Finally, we also now seek to extend the model presented here to different outcome distributions (binary outcomes, time-to-event, count data, etc.), to have a grouped prior for annotations, to model multiple outcomes jointly, and to do all of this using summary statistic as well as individual-level data across different biobanks. Progress in these areas has been made in the AMP literature, see e.g. [35, 36]. Combining data sets together in a principled way without a loss of accuracy, whilst preserving privacy, is key to obtaining the sample sizes that are likely required to fully explore the genetic basis of complex traits.

## Summary

In summary, gVAMP is a different way to create genetic predictors and to conduct variable selection. With increasing sample sizes reducing standard errors, a vast number of genomic regions are being identified as significantly associated with trait outcomes by one-SNP-at-a-time association testing. Such large numbers of findings will make it increasingly difficult to determine the relative importance of a given mutation, especially in whole genome sequence data with dense, highly correlated variants.

Thus, it is crucial to develop statistical approaches fitting all variants jointly and asking whether, given the LD structure of the data, there is evidence for an effect at each locus.

## Data availability

This project uses the UK Biobank data under project number 35520. UK Biobank genotypic and phenotypic data is available through a formal request at (http://www.ukbiobank.ac.uk). It also uses genotypic and phenotypic data from the All of Us study, which is also available through a formal request at (https://www.researchallofus.org/data-tools/data-access/). All summary statistic estimates are released publicly on Dryad: https://doi.org/10.5061/dryad.cz8w9gjjc.

## Code availability

The gVAMP code developed in this work is open source and has been deposited on GitHub, where it is publicly available at https://github.com/medical-genomics-group/gVAMP. The URLs of other software used are listed in the Key Resources Table.

## Acknowledgments

We thank Malgorzata Borczyk for creating the gene burden scores. We thank Robin Beaumont, Amedeo Roberto Esposito, Gareth Hawkes, Philip Schniter, Matthew Stephens, Pragya Sur, Peter Visscher, Michael Weedon and Harry Wright for providing valuable suggestions and comments on earlier versions of the work. This project was funded by a Lopez-Loreta Prize to MM, by an SNSF Eccellenza Grant to MRR (PCEGP3-181181), by an ERC Starting Grant to MM (INF^2^, project number 101161364) and by core funding from ISTA. High-performance computing was supported by the Scientific Service Units (SSU) of ISTA through resources provided by Scientific Computing (SciComp). We would like to acknowledge the participants and investigators of the UK Biobank study. We gratefully acknowledge the All of Us participants for their contributions, without whom this research would not have been possible. We also thank the National Institutes of Health’s All of Us Research Program for making available the participant data [and/or samples and/or cohort] examined in this study.

## Author contribution

MM and MRR conceived the study. AD, MM and MRR designed the study. AD derived the model and the algorithm, with input from MM and MRR. AD wrote the software, with input from MM and MRR. AD, JB, MM, and MRR conducted the analysis and wrote the paper. All authors approved the final manuscript prior to submission.

## Author competing interests

MRR receives research funding from Boehringer Ingelheim for work that is unrelated to that presented here. All other authors declare no competing interests.

## Ethical approval declaration

This project uses UK Biobank data under project 35520. UK Biobank genotypic and phenotypic data is available through a formal request at http://www.ukbiobank.ac.uk. The UK Biobank has ethics approval from the North West Multi-centre Research Ethics Committee (MREC). Methods were carried out in accordance with the relevant guidelines and regulations, with informed consent obtained from all participants. 5]. Informed consent for all All of Us participants is conducted in person or through an eConsent platform that includes primary consent, HIPAA Authorization for Research use of electronic health records and other external health data, and Consent for Return of Genomic Results. The protocol was reviewed by the Institutional Review Board (IRB) of the All of Us Research Program. The All of Us IRB follows the regulations and guidance of the NIH Office for Human Research Protections for all studies, ensuring that the rights and welfare of research participants are overseen and protected uniformly.

## STAR⋆ METHODS

### KEY RESOURCES TABLE

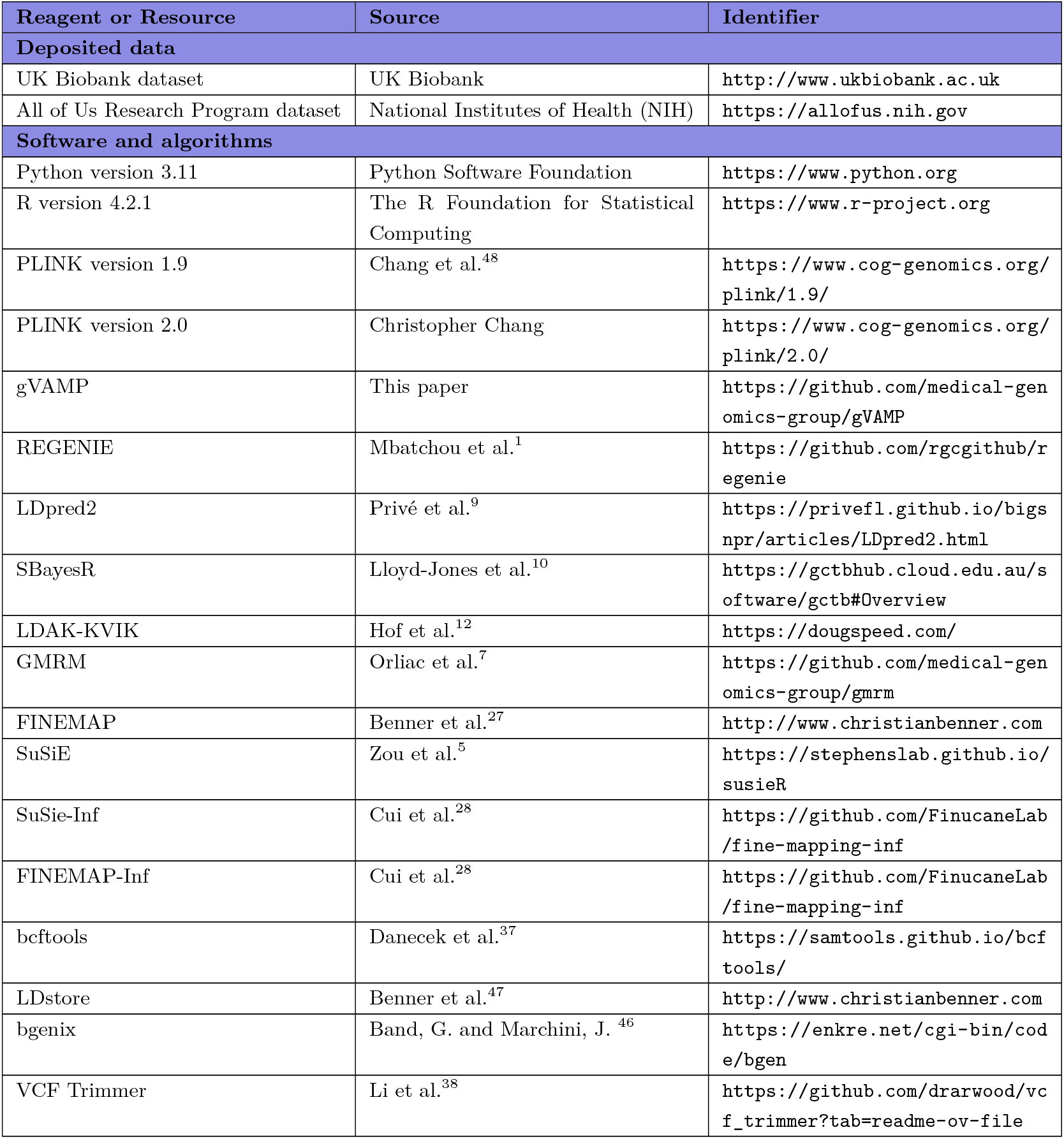

### EXPERIMENTAL MODEL AND SUBJECT DETAILS

In this section, we discuss the data preparation setup. A more detailed description that also motivates our choices further is available in Supplementary Note 2.

#### Participant Inclusion in UK Biobank

The study utilized data from the UK Biobank, and all procedures were covered by ethical approval from the North-West Multicenter Research Ethics Committee, and all participants provided written informed consent. The analysis focused on a sample of European-ancestry UK Biobank individuals. Ancestry inference was based on two criteria: self-reported ethnic background (UK Biobank field 21000-0, selecting coding 1) and genetic ethnicity (UK Biobank field 22006-0, selecting coding 1). Participants were projected onto the first two genotypic principal components (PCs) derived from 2,504 individuals in the 1,000 Genomes project. Individuals were retained if their projections were within PC1 ≤ absolute value 4 and PC2 ≤ absolute value 3. Samples were also excluded based on several UK Biobank quality control procedures:

- Individuals identified as extreme heterozygosity or missing genotype outliers,
- Individuals whose genetically inferred gender did not match their self-reported gender,
- Individuals with putative sex chromosome aneuploidy,
- Individuals excluded from kinship inference,
- Individuals who had withdrawn consent.

Furthermore, 39,592 individuals identified as first-degree relatives were removed from the analyses to avoid potential confounding from shared environmental effects, leaving 419,155 individuals for the whole genome sequence analysis.

#### Whole Genome Sequence (WGS) Data

The study used population-level WGS variants from the UK Biobank DNAnexus platform. Thousands of pVCF files were processed using bcftools [37] and VCF Trimmer [38]. A series of elementary filters were applied to the WGS data:

- Genotype Quality (GQ) ≤ 10.
- Local allele depth (smpl_sum LAD) < 8.
- Missing genotype (F_MISSING) > 0.1.
- Minor Allele Frequency (MAF) < 0.0001.

Additionally, indels were normalized to the most recent reference, redundant data fields were removed, and any duplicate variants sharing the same base pair position were excluded. Due to RAM limitations of 2TBs on the compute nodes, the variant set was reduced. Variants were ranked by MAF, and a PLINK clumping approach was used to remove variants in high linkage disequilibrium (LD) with the most common variants. This was set with a 1000 kb radius and an *R*^2^ threshold of 0.36. The final WGS dataset consisted of 16,854,878 variants.

### Imputed SNP Data

We use version 3 of the imputed autosomal genotype data from the UK Biobank. Genotype probabilities were hard-called for variants with an imputation quality score above 0.3, using a hard-call threshold of 0.1 (genotypes with probability ≤ 0.9 were set to missing). Markers were filtered for missingness (< 5%) and Hardy-Weinberg test *p*-value (> 10^−6^), and only those with an rs identifier were kept, resulting in 23,609,048 SNPs. A primary subset of 8,430,446 autosomal markers was created by selecting markers with MAF ≥ 0.002. Two smaller subsets were also created from the 8.4M set for comparative analyses:

- 2,174,071 SNPs by taking one marker from variants with LD *R*^2^ ≥ 0.8 within a 1MB window,
- 882,727 SNPs by further subsetting markers based on *R*^2^ ≥ 0.5 criteria.

### Whole Exome Sequence (WES) Data Burden Scores

The UK Biobank final release of population-level exome variant call files was processed. High-quality, biallelic sites were retained based on: individual and variant missingness < 10%, Hardy-Weinberg Equilibrium *p*-value > 10^−15^, minimum read coverage depth of 7, and at least one sample per site passing the allele balance threshold > 0.15. Genomic variants in canonical, protein-coding transcripts were annotated using the Ensembl Variant Effect Predictor (VEP) tool. The LOFTEE plugin was used to identify high-confidence (HC) loss-of-function (LoF) variants. For each of the 17,852 genes, a burden score was calculated:

- Score of 2: Individuals homozygous or multiple heterozygous for LoF variants.
- Score of 1: Individuals with a single heterozygous LoF variant.
- Score of 0: All other individuals.

### Phenotypic Records

The prepared DNA datasets were linked to phenotypic measurements from the UK Biobank. For the imputed SNP data analyses, 13 quantitative traits (including 7 blood-based biomarkers and 6 other quantitative measures) were selected. These traits were chosen because they showed ≥ 15% SNP heritability and ≥ 5% out-of-sample prediction accuracy in previous work. The sample was divided into a training set and a testing set. The test set consisted of 15,000 individuals who were unrelated (SNP marker relatedness < 0.05) to the individuals in the training set. Prior to analysis, all phenotypic values were adjusted for several covariates: participant’s recruitment age, genetic sex, north-south and east-west location coordinates, measurement center, genotype batch, and the leading 20 genetic principal components.

## METHOD DETAILS

### gVAMP installation

The primary method used for analysis in this paper is gVAMP. Its source code is available at https://github.com/medical-genomics-group/gVAMP and can be downloaded using the `git` clone command. Compilation in a High-Performance Computing (HPC) environment requires `gcc, openmpi` and `boost` libraries. Upon loading the required modules (e.g., `module load gcc openmpi boost`), the executable (e.g., `main_real.exe`) is built from the C++ source files using an `mpic++` command with relevant flags (such as `-march=native, -Ofast`, and `-fopenmp`).

To execute an analysis, gVAMP requires specific inputs passed as command-line arguments, primarily the genotype data in a standard PLINK `.bed` file (specified with `–bed-file`), the phenotype data in a `.phen` file (`–phen-files`), and other model parameters like the number of samples (`–N`) and total markers (`–Mt`). A typical execution command integrating these inputs is shown below:

~~~
# Example with 30 processes
time mpirun -np 30 /path/to/main_real.exe \
  --bed-file /path/to/genotypes.bed \
  --phen-files /path/to/phenotypes.phen \
  --N 401452 \
  --Mt 2174071 \
  … [other arguments]
~~~

### gVAMP algorithm overview

gVAMP extends EM-VAMP, introduced in [14, 24, 25], in which the prior parameters are adaptively learnt from the data via EM, and it is an iterative procedure consisting of two steps: *(i)* denoising, and *(ii)* linear minimum mean square error estimation (LMMSE). The denoising step accounts for the prior structure given a noisy estimate of the signal ***β***, while the LMMSE step utilizes phenotype values to further refine the estimate by accounting for the LD structure of the data. Further information on historical development and an informal description of AMP-based approaches for Bayesian regressions are presented in Supplementary Note 3.

A key feature of the algorithm is the so called *Onsager correction*: this is added to ensure the asymptotic normality of the noise corrupting the estimates of ***β*** at every iteration. Here, in contrast to MCMC or other iterative approaches, the normality is guaranteed under mild assumptions on the normalized genotype matrix. This property allows a precise performance analysis via state evolution and, consequently, the optimization of the method.

In particular, the quantity *γ*_1,*t*_ in line 7 of Algorithm 1 is the state evolution parameter tracking the error incurred by ***r***_1,*t*_ in estimating ***β*** at iteration *t*. The state evolution result gives that ***r***_1,*t*_ is asymptotically Gaussian, i.e., for sufficiently large *N* and *P*, ***r***_1,*t*_ is approximately distributed as 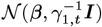. Here, ***β*** represents the signal to be estimated, with the prior learned via EM steps at iteration *t*:

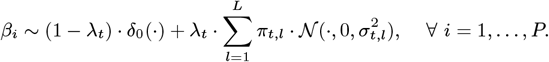

Compared to Equation (2), the subscript *t* in *λ*_*t*_, *π*_*t,l*_, *σ*_*t,l*_ indicates that these parameters change through iterations, as they are adaptively learned by the algorithm. Similarly, ***r***_2,*t*_ is approximately distributed as 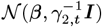. The Gaussianity of ***r***_1,*t*_, ***r***_2,*t*_ is enforced by the presence of the Onsager coefficients *α*_1,*t*_ and *α*_2,*t*_, see lines 17 and 22 of Algorithm 1, respectively. We also note that *α*_1,*t*_ (resp. *α*_2,*t*_) is the state evolution parameter linked to the error incurred by 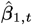 (resp. 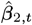).

The vectors ***r***_1,*t*_, ***r***_2,*t*_ are obtained after the LMMSE step, and they are further improved via the denoising step, which respectively gives 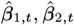. In the denoising step, we exploit our estimate of the approximated posterior by computing the conditional expectation of ***β*** with respect to ***r***_1,*t*_, ***r***_2,*t*_ in order to minimize the mean square error of the estimated effects. For example, let us focus on the pair 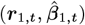 (analogous considerations hold for 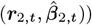. Then, we have that

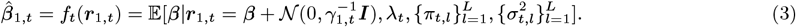

Here, *f*_*t*_ : ℝ → ℝ denotes the denoiser at iteration *t* and the notation *f*_*t*_(***r***_1,*t*_) assumes that the denoiser *f*_*t*_ is applied component-wise to elements of ***r***_1,*t*_. We give explicit formulas for denoisers *f*_*t*_ as well as the Onsager corrections in the Supplementary Note 4. Moreover, note that, in line 15 of Algorithm 1, we take this approach one step further by performing an additional step of damping, see “gVAMP algorithm stability” below.

If one has access to the singular value decomposition (SVD) of the data matrix ***X***, the per-iteration complexity is of order 𝒪 (*NP*). However, at biobank scales, performing the SVD is computationally infeasible. Thus, the linear system (*γ*_*ϵ,t*_***X***^*T*^ ***X*** + *γ*_2,*t*_***I***)^−1^(*γ*_*ϵ,t*_***X***^*T*^ ***y*** + *γ*_2,*t*_***r***_2,*t*_) (see line 21 of Algorithm 1) needs to be solved using an iterative method, in contrast to having an analytic solution in terms of the elements of the singular value decomposition of ***X***. We approximate the solution of the linear system (*γ*_*ϵ,t*_***X***^*T*^ ***X*** + *γ*_2,*t*_***I***)^−1^(*γ*_*ϵ,t*_***X***^*T*^ ***y*** + *γ*_2,*t*_***r***_2,*t*_) with a symmetric and positive-definite matrix via the *conjugate gradient method* (CG), see Algorithm 2 in Supplementary Note 5. If *κ* is the condition number of *γ*_*ϵ,t*_***X***^*T*^ ***X*** + *γ*_2,*t*_***I***, the method requires 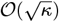 iterations to return a reliable approximation.

Additionally, inspired by [40], we initialize the CG iteration with an estimate of the signal from the previous iteration of gVAMP. This warm-starting technique leads to a reduced number of CG steps that need to be performed and, therefore, to a computational speed-up. However, this comes at the expense of potentially introducing spurious correlations between the signal estimate and the Gaussian error from the state evolution. Such spurious correlations may lead to algorithm instability when run for a large number of iterations (also extensively discussed below). This effect is prevented by simply stopping the algorithm as soon as the *R*^2^ measure on the training data starts decreasing.

In order to calculate the Onsager correction in the LMMSE step of gVAMP (see line 22 of Algorithm 1), we use the Hutchinson estimator [41] to estimate the quantity Tr[(*γ*_*ϵ,t*_***X***^*T*^ ***X*** + *γ*_2,*t*_***I***)^−1^]*/P* . We recall that this estimator is unbiased, in the sense that, if ***u*** has i.i.d. entries equal to −1 and +1 with the same probability, then

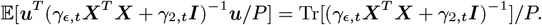

Furthermore, in order to perform an EM update for the noise precision *γ*_*ϵ*_, one has to calculate the trace of a matrix which is closely connected to the one we have seen in the previous paragraph. In order to do so efficiently, i.e., to avoid solving another large-dimensional linear system, we store the inverted vector (*γ*_*ϵ,t*_***X***^*T*^ ***X*** + *γ*_2,*t*_***I***)^−1^***u*** and reuse it again in the EM update step.

The VAMP approach in [14] assumes exact knowledge of the prior on the signal ***β***, which deviates from the setting in which genome-wide association studies are performed. Hence, we adaptively learn the signal prior from the data using expectation-maximization (EM) steps, see lines 8 and 28 of Algorithm 1. This leverages the variational characterization of EM-VAMP [24], and its rigorous theoretical analysis presented in [25]. In the Supplementary Note 6, we summarize the hyperparameter estimation results derived based upon [42] in the context of our model.

### gVAMP implementation

Our open-source gVAMP software (https://github.com/medical-genomics-group/gVAMP) is implemented in C++, and it incorporates parallelization using the OpenMP and MPI libraries. MPI parallelization is implemented in a way that the columns of the normalized genotype matrix are approximately equally split between the workers. OpenMP parallelization is done on top of that and used to further boost performance within each worker by simultaneously performing operations such as summations within matrix vector product calculations. Moreover, data streaming is employed using a lookup table, enabling byte-by-byte processing of the genotype matrix stored in PLINK format with entries encoded to a set {0, 1, 2}:

#### Algorithm 1

gVAMP

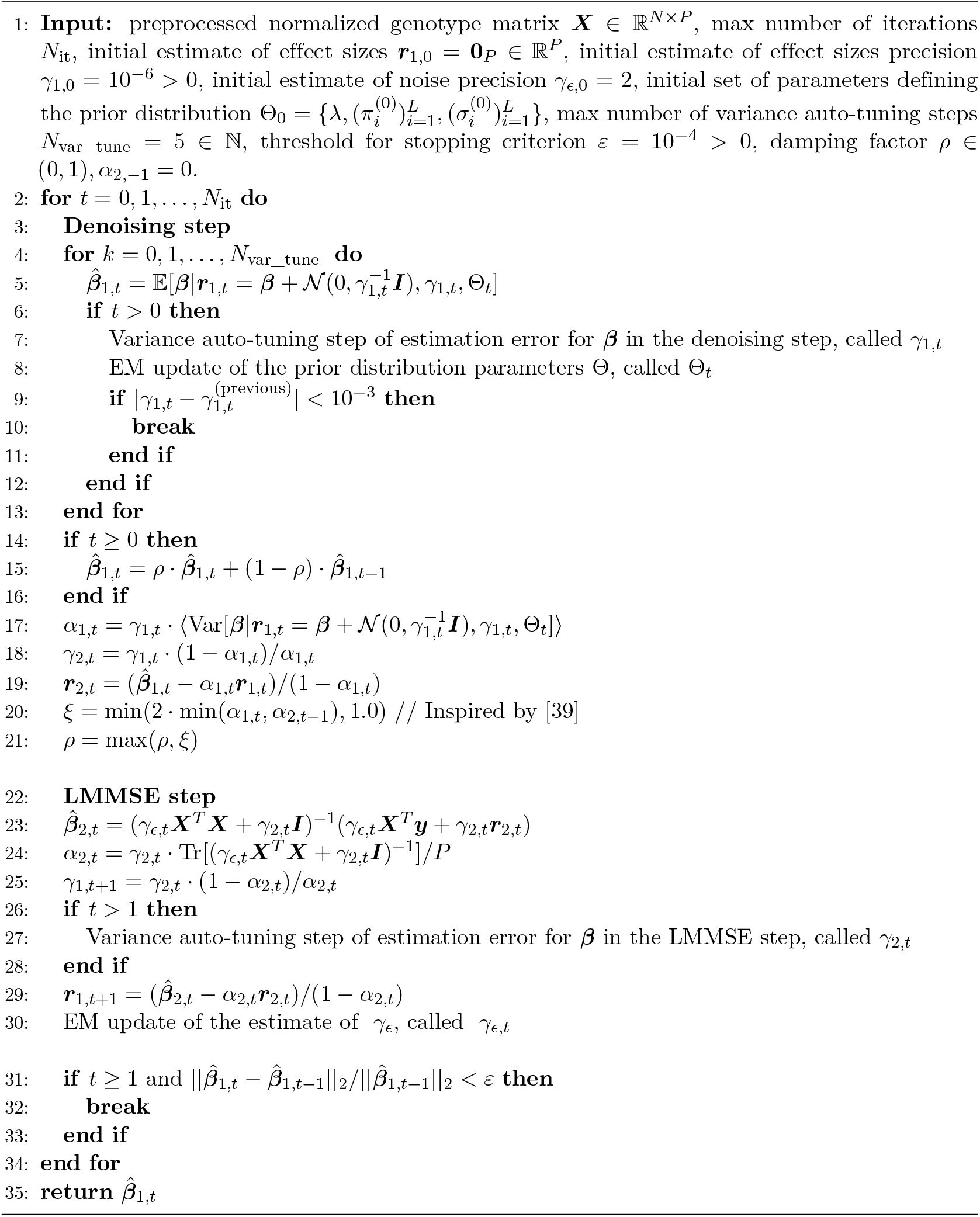

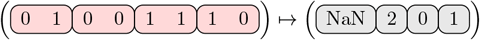

The lookup table enables streaming in the data in bytes, where every byte (8 bits) encodes the information of 4 individuals. This reduces the amount of memory needed to load the genotype matrix. In addition, given a suitable computer architecture, our implementation supports SIMD instructions which allow handling four consecutive entries of the genotype matrix simultaneously. To make the comparisons between different methods fair, the results presented in the paper do not assume usage of SIMD instructions. Additionally, we emphasize that all calculations take un-standardized values of the genotype matrix in the form of standard PLINK binary files, but are conducted in a manner that yields the parameter estimates one would obtain if each column of the genotype matrix was standardized.

### gVAMP algorithm stability

We find that the application of existing EM-VAMP algorithms to the UK Biobank dataset leads to diverging estimates of the signal. This is due to the fact that the data matrix (the SNP data) might not conform to the properties required in [14], especially that of right-rotational invariance. Furthermore, incorrect estimation of the noise precision in line 28 of Algorithm 1 may also lead to instability of the algorithm, as previous applications of EM-VAMP generally do not leave many hyperparameters to estimate.

To mitigate these issues, different approaches have been proposed including row or/and column normalization, damping (i.e., doing convex combinations of new and previous estimates) [43], and variance auto-tuning [25]. In particular, to prevent EM-VAMP from diverging and ensure it follows its state evolution, we empirically observe that the combination of the following techniques is crucial:

1. We perform *damping* in the space of denoised signals. Thus, line 15 of Algorithm 1 reads as

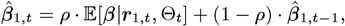

in place of 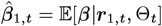. Here, *ρ* ∈ (0, 1) denotes the damping factor. This ensures that the algorithm is making smaller steps when updating a signal estimate.
2. We perform *auto-tuning* of *γ*_1,*t*_ via the approach from [25]. Namely, in the auto-tuning step, one refines the estimate of *γ*_1,*t*_ and the prior distribution of the effect size vector ***β*** by jointly re-estimating them. If we denote the previous estimates of *γ*_1,*t*_ and Θ with 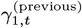 and Θ^(previous)^, then this is achieved by setting up an expectation-maximization procedure whose aim is to maximize

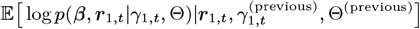

with respect to *γ*_1,*t*_ and Θ.
3. We *filter* the design matrix for first-degree relatives to reduce the correlation between rows, which has the additional advantage of avoiding potential confounding of shared-environmental effects among relatives.

### gVAMP model parameters for the analysis on Imputed data

For the analyses of the 13 UK Biobank phenotypes using the 887,060 and 2,174,071 imputed SNP sets, a damping factor *ρ* of 0.1 was used. The prior was initialized with a spike at zero and 22 slab mixtures. The variances of these mixtures followed a geometric progression in the interval [10^−6^, 10], and their probabilities followed a geometric progression with a factor of 1/2. The initial probability of being assigned to the zero mixture was 97%. This configuration works well for all phenotypes. We also note that our inference of the number of mixtures, their probabilities, their variances and the SNP marker effects is not dependent upon specific starting parameters for the analyses of the 2,174,071 and 882,727 SNP datasets, and the algorithm is rather stable for a range of initialization choices. Similarly, the algorithm is stable for different choices of the damping *ρ*, as long as said value is not too large.

Generally, appropriate starting parameters are not known in advance and this is why we learn them from the data within the EM steps of our algorithm. However, it is known that EM can be sensitive to the starting values given and, thus, we recommend initialising a series of models at different values to check that this is not the case (similar to starting multiple Monte Carlo Markov chains in standard Bayesian methods). The feasibility of this recommendation is guaranteed by the significant speed-up of our algorithm compared to existing approaches (see Supplementary Note 1, Figure S21).

Given the same data, gVAMP yields estimates that more closely match GMRM when convergence in the training *R*^2^, SNP heritability, residual variance, and out-of-sample test *R*^2^ are smoothly monotonic within around 10-40 iterations. Following this, training *R*^2^, SNP heritability, residual variance, and out-of-sample test *R*^2^ may then begin to slightly decrease as the number of iterations becomes large. Thus, as a stopping criterion for the 2,174,071 and 882,727 SNP datasets, we choose the iteration that maximizes the training *R*^2^.

We highlight the iterative nature of our method. Thus, improved computational speed and more rapid convergence is achieved by providing better starting values for the SNP marker effects. Specifically, when moving from 2,174,071 to 8,430,446 SNPs, only columns with correlation *R*^2^ ≥ 0.8 are being added back into the data. Thus, for the 8,430,446 SNP set, we initialise the model with the converged SNP marker and prior estimates obtained from the 2,174,071 SNP runs, setting to 0 the missing markers. Furthermore, we lower the value of the damping factor *ρ*, with typical values being 0.05 and 0.01. We experiment both with using the noise precision from the initial 2,174,071 SNP runs and with setting it to 2. We then choose the model that leads to a smoothly monotonic curve in the training *R*^2^. We observe that SNP heritability, residual variance, and out-of-sample test *R*^2^ are also smoothly monotonic within 25 iterations. Thus, as a stopping criterion for the 8,430,446 SNP dataset, we choose the estimates obtained after 25 iterations for all the 13 traits. We follow the same process when extending the analyses to include the WES rare burden gene scores.

### gVAMP model parameters for the analysis of human height in WGS data

We apply gVAMP to the WGS data to analyse human height using the largest computational instance currently available on the DNAnexus platform, employing 128 cores and 1921.4 GB total memory. Efficient C++ gVAMP implementation allows for parallel computing, utilizing OpenMP and MPI libraries. Here, we split the memory requirements and computational workload between 2 OpenMP threads and 64 MPI workers. The prior initialization of gVAMP is similar to that used in the simulation study, except that we opt for a sparser prior with probability 99.5% of variants being assigned to the 0 mixture. As before, the SNP marker effect sizes are initialised with 0. Based on the experiments in the UK Biobank imputed dataset, in WGS we set the initial damping factor *ρ* to 0.1, and adjust it to 0.05 from iteration 4 onward, which stabilizes the algorithm and leads us to stop it at iteration 18. We also note that, after the first few iterations, the marker inclusion probabilities are stable.

In the simulation study on WGS data from chromosomes 11-15, gVAMP was run with a damping factor *ρ* of 0.05. The prior was initialized with a spike at zero and 22 slab mixtures. The mixture variances followed a geometric progression in the interval [10^−7^, 1], and the mixture probabilities followed a geometric progression with a factor of 1/2. The initial probability of a variant being assigned to the zero mixture was 99%. All variant effect sizes were initialized to 0, and the method was run until iteration 30.

### Polygenic risk scores and SNP heritability

gVAMP produces SNP effect estimates that can be directly used to create polygenic risk scores. The estimated effect sizes are on the scale of normalised SNP values, i.e., 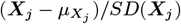, with the column mean and *SD*(***X***_*j*_) the standard deviation, and thus SNPs in the out-of-sample prediction data must also be normalized. We provide an option within the gVAMP software to do phenotypic prediction, returning the adjusted prediction *R*^2^ value when given input data of a PLINK file and a corresponding file of phenotypic values. gVAMP estimates the SNP heritability as the phenotypic variance (equal to 1 due to normalization) minus 1 divided by the estimate of the noise precision, i.e., 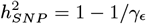.

### Comparison to polygenic risk scores obtained by other methods

We compare gVAMP to an MCMC sampler approach (GMRM) with a similar prior (the same number of starting mixtures) as presented in [7]. We select this comparison as the MCMC sampler was demonstrated to exhibit the highest genomic prediction accuracy up to date [7]. We run GMRM for 2000 iterations, taking the last 1800 iterations as the posterior. We calculate the posterior means for the SNP effects and the posterior inclusion probabilities of the SNPs belonging to the non-zero mixture group. GMRM estimates the SNP heritability in each iteration by sampling from an inverse *χ*^2^ distribution using the sum of the squared regression coefficient estimates. We also compare to the LDAK-KVIK elastic net software[12], run using their suggested parameters and default options.

Finally, we also compare gVAMP to the summary statistics prediction methods LDpred2 [9] and SBayesR [10] using summary statistic data of 8,430,446 SNPs obtained from the REGENIE software. For SBayesR, following the recommendation on the software webpage (https://cnsgenomics.com/software/gctb/#SummaryBayesianAlphabet), after splitting the genomic data per chromosomes, we calculate the so-called *shrunk* LD matrix, which use the method proposed by [44] to shrink the off-diagonal entries of the sample LD matrix toward zero based on a provided genetic map. We make use of all the default values: <monospace>--genmap-n</monospace> 183, <monospace>--ne</monospace> 11400 and <monospace>--shrunk-cutoff</monospace> 10^−5^. Following that, we run the SBayesR software using summary statistics generated via the REGENIE software (see “Mixed linear association testing” below) by grouping several chromosomes in one run. Namely, we run the inference jointly on the following groups of chromosomes: {1}, {2}, {3}, {4}, {5, 6}, {7, 8}, {9, 10, 11}, {12, 13, 14} and {15, 16, 17, 18, 19, 20, 21, 22}. This allows to have locally joint inference, while keeping the memory requirements reasonable. All the traits except for Blood cholesterol (CHOL) and Heel bone mineral density T-score (BMD) give non-negative *R*^2^; CHOL and BMD are then re-run using the option to remove SNPs based on their GWAS *p*-values (threshold set to 0.4) and the option to filter SNPs based on LD R-Squared (threshold set to 0.64). For more details on why one would take such an approach, one can check https://cnsgenomics.com/software/gctb/#FAQ. As the obtained test *R*^2^ values are still similar, as a final remedy, we run standard linear regression over the per-group predictors obtained from SBayesR on the training dataset. Following that, using the learned parameters, we make a linear combination of the per-group predictors in the test dataset to obtain the prediction accuracy given in the table.

### Replication of results in All of Us

The analysis used whole genome sequence (WGS) data from the All of Us cohort [45]. Informed consent for all participants is conducted in person or through an eConsent platform that includes primary consent, HIPAA Authorization for Research use of electronic health records and other external health data, and Consent for Return of Genomic Results. The protocol was reviewed by the Institutional Review Board (IRB) of the All of Us Research Program. The All of Us IRB follows the regulations and guidance of the NIH Office for Human Research Protections for all studies, ensuring that the rights and welfare of research participants are overseen and protected uniformly.

A cohort of 226,147 individuals of European genetic ancestry with height data was selected for the replication analysis. The phenotypic values for height were adjusted for sex and 16 available principal components (PCs) within the All of Us dataset. A marginal regression was performed on the standardized WGS dosage data. This analysis was conducted using the Hail Python library. The results of this marginal regression (estimating one variant at a time) from All of Us were then compared to the joint-effect estimates generated by gVAMP from the WGS runs in the UK Biobank data for standing height as a trait.

### Simulation study for WGS data

To demonstrate the usefulness of gVAMP as a signal localization tool, we investigate a series of 12 simulation scenarios in which the signal is planted on top of the observed whole genome sequence data of chromosomes 11-15, which consists of 3,285,117 variants and 411,536 individuals. At a scale of over 3 million WGS variants and over 400,000 individuals, cloud computing costs naturally restrict the number of scenarios we can explore, especially as we have to benchmark against a range of other fine-mapping approaches. The simulation details for each of the 12 scenarios we considered are reported in Table S1 and described below.

The signal itself is simulated from a Bernoulli-Gaussian prior in which the variance of the slab part is determined as the ratio of the phenotypic variance explained by the causal variants and the number of causal variants. We fix the phenotypic variance explained by the causal variants to be 7.25% and vary the number of causal variants as either 500 or 1000. Given that the length of the DNA of chromosomes 11-15 is ∼ 18.8% of the total autosomal DNA length, the settings we use are equivalent to a trait with a proportion of variance explained of ∼ 39% spread over ∼ 2700 to ∼ 5300 causal variants genome-wide. Our objective in the simulation study is to access the performance of our model at scale in settings similar to our WGS analysis of human height, but where power is sufficient to accurately compare variable selection methods. 39%/5300 = 0.0074% is a value similar to that expected for human height 70%/10000 = 0.007% suggested by recent results[26]. In the settings we explore, we vary *γ*, the percentage of non-zero effects attributed to common variants (those with minor allele frequency ≥ 0.05). In simulation scenarios with *γ* = 0%, we spread the signals uniformly across variants maintaining our minimal distance. *γ* = 50% allows common variants in the interval [0.05, 0.5] to be causal with the probability 50%. When selecting causal variants within these two settings, we do so at random, but to spread variants across the DNA we enforce a minimal distance between consecutive causal variants of size 100,000 base pairs. This minimal distance facilitates a simple direct comparison of variable selection methods to localize a single signal within a given genomic region, but restricts us to a smaller absolute number of causal variants than may be expected for some traits genome-wide. However, as noted above, the average expected contribution to variance is in-line with that expected for human height and, thus, our simulation represents a realistic potential architecture.

We vary *α*, the power relationship between effect size and minor allele frequency. For each SNP position *j*, upon randomly choosing a mixture group *l* it belongs to, according to the true effect probability distribution, we draw its effect size from 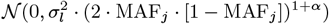. Given a desired heritability *h*^2^ of the simulated trait, we normalize the estimates so that the sum of squares of all simulated effects is equal to *h*^2^. This model conforms exactly to that described in [46] and we set values of −1, −0.5, and −0.25 to cover those reported for complex traits [46]. As described in [46], when *α* = 0 the unscaled effect-size variance is invariant to MAF, indicative of selective neutrality. The uniform heritability model (*α* = −1) implies that the unscaled effect-size variance increases as MAF decreases, which is interpreted as a signal of negative (or purifying) selection.

All simulated traits are adjusted by the first 20 principal components following standard practice. For running SuSiE and SuSiE-inf (v1.4), we use the standard fine-mapping with infinitesimal effects v1.2 Python framework provided by the authors [28]. The framework allows to initialize FINEMAP-inf using tau-squared and sigma-squared parameters generated by SuSiE-inf to improve accuracy. When running FINEMAP, we use a shotgun stochastic search subprogram. We perform genome-wide marginal testing and define fine-mapping blocks of size 3Mbp around a marginal discovery, passing the Bonferroni-adjusted p-value threshold of 0.05. When the 3Mbp contained or neighboured multiple causal variants, we algorithmically allow for merging the overlapping regions, allowing FINEMAP and SuSiE to run on region sizes of 6Mbp. In fact, we found that this gave favorable performance to minimize the probability of counting an association as false, when it is simply correlated to a neighboring true causal variant. For each region, we create an index of variants using bgenix [47], compute LD matrices using LDstore v2.0 [48], and marginal estimates using PLINK 1.9 [49].

We run the gVAMP method using as prior initialization a spike at zero and 22 slab mixtures. We let the variance of those mixtures follow a geometric progression in the interval [10^−7^, 1], and we let the probabilities follow a geometric progression with factor 1/2. The prior probability 1 − *λ* of variants being assigned to the 0 mixture is initialized to 99%. Variant effect sizes are initialised with 0. Based on the experiments in the UK Biobank imputed dataset, in WGS we use damping factor *ρ* = 0.05 and run the method until iteration 30 when the hyper-parameters stabilize.

We then conduct an additional simulation study focusing on the inclusion of WES burden scores alongside WGS variants. For the settings of (*α, γ*) ∈ {(−1, 0), (−1, 50), (−0.5, 0), (−0.5, 50)}, we used the same whole genome sequence data of chromosomes 11-15 described above. We simulated 500 underlying causal variants with a total heritability of *h*^2^ = 7.25%, allocating 75% of the causal markers (CM) to randomly selected WGS variants and 25% to randomly selected WES burden scores from the same chromosomes. The WES burden scores exhibit effect sizes drawn from the normal distribution with variance three-fold larger than those of individual WGS variants. This gives the expectation that the variance attributable to WGS variants should equal that of the WES burden scores. We then ran gVAMP using the same initialization described above.

Finally, we conduct a further simulation study where we vary effects sizes across genomic annotations. Specifically, we simulated 500 causal variants with a total heritability of *h*^2^ = 7.25%, stratifying them into coding and non-coding regions of the genome. We vary the parameter *γ*_annot_ ∈ {0, 50, 100} which specifies the percentage of causal variants assigned to coding (exonic) regions. This simulation was performed only for settings *a, b, e*, and *f*, described in Supplementary Table S1. We then ran gVAMP using the same initialization described above.

### QUANTIFICATION AND STATISTICAL ANALYSIS

#### gVAMP association testing

In line with the standard outputs of MCMC methods, the AMP framework offers the possibility of calculating posterior inclusion probabilities (PIPs) jointly for each of the SNPs. Once the prior information in a particular iteration is updated, we use Equation (4) below to approximate the posterior inclusion probability of each SNP. To conduct variable selection, we apply a threshold of PIP ≥ 0.95.

The gVAMP framework provides posterior inclusion probability testing for detecting associations. The calculation uses the output of the state evolution, specifically the noisy estimate of the effects ***r***_1,*t*_ and its associated precision *γ*_1,*t*_ at a given iteration *t*. The posterior inclusion probability for a single SNP *j*, denoted as *ζ*_*j*_, is calculated using the following equation from the Supplementary Note 6:

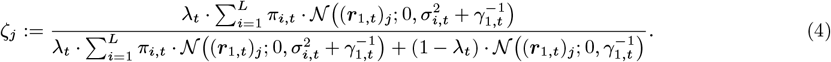

In this formula, the terms represent the following theoretical quantities:

- *ζ*_*j*_ : The posterior probability that SNP *j* has a non-zero effect,
- (***r***_1,*t*_)_*j*_ : The noisy estimate of the effect for SNP *j* at iteration *t*,
- *γ*_1,*t*_: The precision of the estimate (***r***_1,*t*_)_*j*_, tracked by state evolution,
- *λ*_*t*_: The adaptively learned sparsity rate, which is the prior probability of a non-zero effect,
- *L*: The number of Gaussian components in the “slab” part of the prior.
- *π*_*i,t*_: The prior probability of being in the *i*-th Gaussian mixture, given that the effect is non-zero,
- 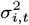: The variance of the *i*-th Gaussian mixture component.
- 𝒩 (·; *µ, ν*): The probability density function of a Gaussian distribution with mean *µ* and variance *ν*.

### Mixed linear model association testing

We conduct mixed linear model association testing using a leave-one-chromosome-out (LOCO) estimation approach on the 8,430,446 and 2,174,071 imputed SNP markers. LOCO association testing approaches have become the field standard and they are two-stage: a subset of markers is selected for the first stage to create genetic predictors; then, statistical testing is conducted in the second stage for all markers one-at-a-time. We consider REGENIE [1], as it is a recent commonly applied approach and LDAK-KVIK[12]. We also compare to GMRM [7], a Bayesian linear mixture of regressions model that has been shown to outperform REGENIE for LOCO testing. For the first stage of LOCO, REGENIE is given 887,060 markers to create the LOCO genetic predictors, even if it is recommended to use 0.5 million genetic markers. We compare the number of significant loci obtained from REGENIE to those obtained if one were to replace the LOCO predictors with: *(i)* those obtained from GMRM using the LD pruned sets of 2,174,071 and 887,060 markers; and *(ii)* those obtained from gVAMP at all 8,430,446 markers and the LD pruned sets of 2,174,071 and 887,060 markers. We note that obtaining predictors from GMRM at all 8,430,446 markers is computationally infeasible, as using the LD pruned set of 2,174,071 markers already takes GMRM several days. In contrast, gVAMP is able to use all 8,430,446 markers and still be faster than GMRM with the LD pruned set of 2,174,071 markers. LDAK-KVIK is given 2,174,071 markers to facilitate a direct comparison with gVAMP and we compare *(i)* placing the predictors of gVAMP directly within the LDAK-KVIK step 2; *(ii)* using gVAMP step 2 procedure which is identical to that of REGENIE.

LOCO testing does not control for linkage disequilibrium within a chromosome. Thus, to facilitate a simple, fair comparison across methods, we clump the LOCO results obtained with the following PLINK commands: `--clump-kb` 1000 `--clump-r2` 0.01 `--clump-p1` 0.00000005. Therefore, within 1Mb windows of the DNA, we calculate the number of independent associations (squared correlation ≤ 0.01) identified by each approach that pass the genome-wide significance testing threshold of 5 · 10^−8^. As LOCO can only detect regions of the DNA associated with the phenotype and not specific SNPs, given that it does not control for the surrounding linkage disequilibrium, a comparison of the number of uncorrelated genome-wide significance findings is conservative. A full benchmark of approaches in a simulation study is also provided in Supplementary Note 1, where we consider a wider range of metrics, scenarios and comparisons.

## Supplementary information

## Supplementary Tables

**Table S1.** Simulation settings using whole genome sequence (WGS) data of chromosomes 11 to 15 in the UK Biobank. Each setting varies in the number of causal variants (CV), the frequency-based enrichment parameter (*γ*), and the effect-size–MAF relationship (*α*, see Methods for the description of the parameters). All scenarios shown use *h*^2^ = 7.25%.

| ID | Number of causal variants | $\gamma$ | $\alpha$ |
| --- | --- | --- | --- |
| a | 500 | 0 | -1 |
| b | 500 | 50 | -1 |
| c | 1000 | 0 | -1 |
| d | 1000 | 50 | -1 |
| e | 500 | 0 | -0.5 |
| f | 500 | 50 | -0.5 |
| g | 1000 | 0 | -0.5 |
| h | 1000 | 50 | -0.5 |
| i | 500 | 0 | -0.25 |
| j | 500 | 50 | -0.25 |
| k | 1000 | 0 | -0.25 |
| l | 1000 | 50 | -0.25 |

**Table S2.** The 13 UK Biobank traits used within the study. Phenotypic names and their codes used in the study. The sample size, *N*, gives the number of individuals with training data measures.

| Phenotype | Code | Sample size, $N$ |
| --- | --- | --- |
| Blood: cholesterol | CHOL | 395,025 |
| Blood: eosinophill count | EOSI | 401,452 |
| Blood: glycated haemoglobin | HbA1c | 394,912 |
| Blood: High density lipoprotein | HDL | 360,286 |
| Blood: mean corpuscular haemoglobin | MCH | 402,201 |
| Blood mean corpuscular volume | MCV | 402,202 |
| Red blood cell count | RBC | 402,204 |
| Body mass index | BMI | 413,595 |
| Diastolic blood pressure | DBP | 377,358 |
| Forced vital capacity | FVC | 376,724 |
| Heel bone mineral density | BMD | 231,693 |
| Standing height | HT | 414,055 |
| Systolic blood pressure | SBP | 377,347 |

**Table S3.** Genome-wide significant associations for 13 UK Biobank traits from the software gVAMP and LDAK-KVIK at 8,430,446 genetic variants from mixed linear model association leave-one-chromosome-out (MLMA-LOCO) testing. “LDAK 2M LOCO” refers to MLMA-LOCO testing at 8,430,446 SNPs, using the LDAK-KVIK software in step 1, with 2,174,071 SNP markers. “gVAMP 2M LOCO 1” refers to MLMA-LOCO testing at 8,430,446 SNPs, using the framework presented here, where in step 1 2,174,071 markers are used and the second step is identical to that of LDAK-KVIK. “gVAMP 2M LOCO 2” is the same as “gVAMP 2M LOCO 1” but with the second step identical to REGENIE. The values are the number of genome-wide significant linkage disequilibrium independent associations selected based upon a standard *p*-value threshold of less than 5 · 10^−8^ and *R*^2^ between SNPs in a 1 Mb genomic segment of less than 1%.

| Trait | LDAK<br>2M LOCO | gVAMP<br>2M LOCO<br>1 | gVAMP<br>2M LOCO<br>2 |
| --- | --- | --- | --- |
| CHOL | 482 | 625 | 603 |
| EOSI | 549 | 666 | 630 |
| HbA1c | 321 | 400 | 385 |
| HDL | 639 | 855 | 812 |
| MCH | 690 | 858 | 810 |
| MCV | 903 | 1128 | 1062 |
| RBC | 856 | 1140 | 1079 |
| BMI | 726 | 1202 | 1175 |
| DBP | 303 | 357 | 419 |
| FVC | 577 | 759 | 721 |
| BMD | 549 | 681 | 668 |
| HT | 2703 | 4747 | 4452 |
| SBP | 323 | 410 | 388 |

## Supplementary Figures

**Figure S1.**
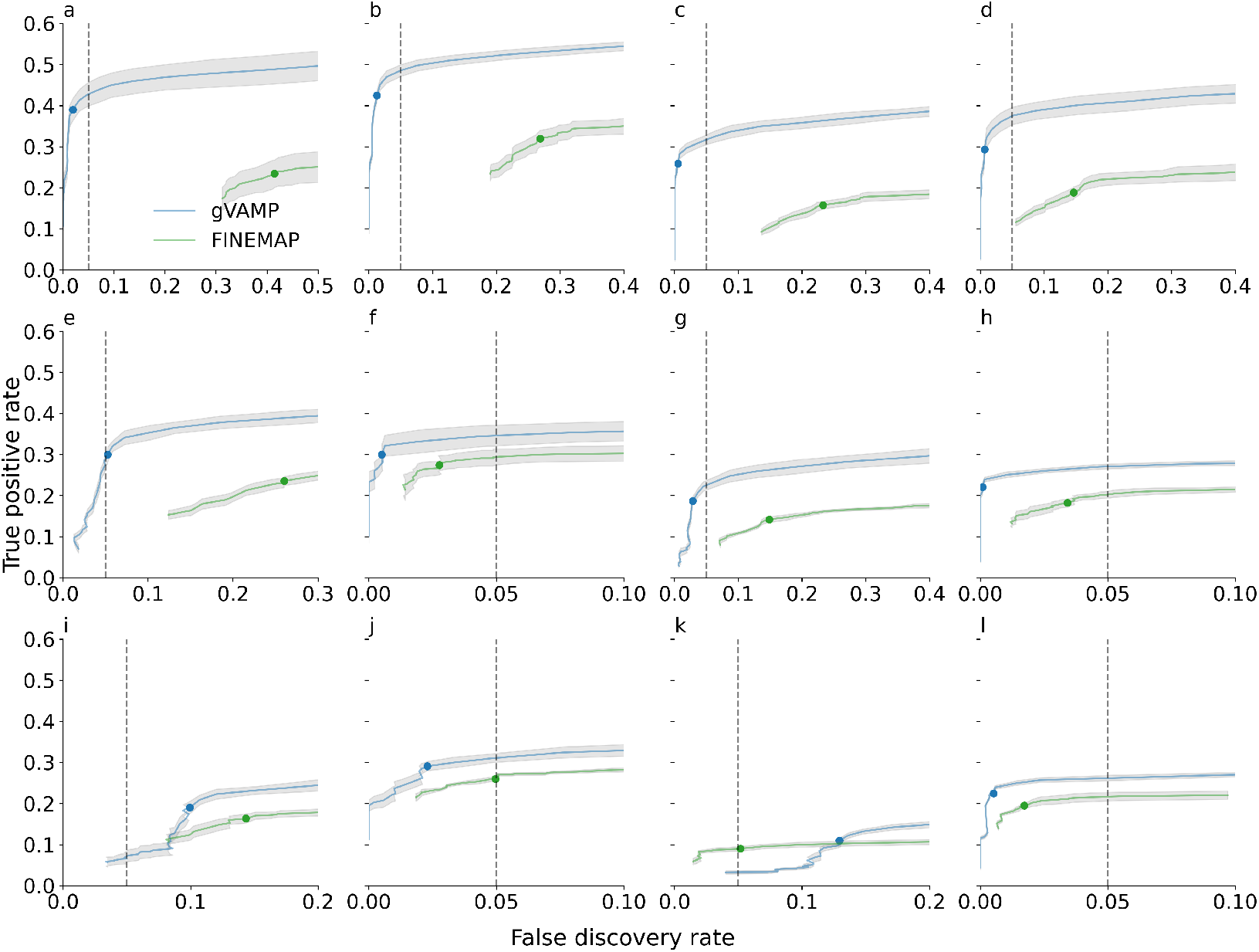
Comparison of the true positive rate (TPR, *y*-axis) versus the false discovery rate (FDR, *x*-axis) between gVAMP and marginal GWAS association testing plus fine-mapping implemented in the FINEMAP algorithm across 12 simulation scenarios using whole genome sequence (WGS) data of chromosomes 11 to 15 in the UK Biobank. For the detailed description of different simulation settings, see Methods and Supplementary Table S1. We use a Bonferroni corrected p-value significance threshold for marginal GWAS testing and then having identified associated regions we examine the ability of FINEMAP (green) to conduct variable selection across posterior inclusion probability (PIP) thresholds. For gVAMP (blue), variable selection is conducted from PIP values obtained from a single model run genome-wide. Error bands give the standard deviation across 5 simulation replicates. The blue and green dots give the TPR and FDR at PIP ≥ 0.95, which should correspond to an FDR of ≤ 0.05 if the model correctly conducts variable selection based on PIP values.

**Figure S2.**
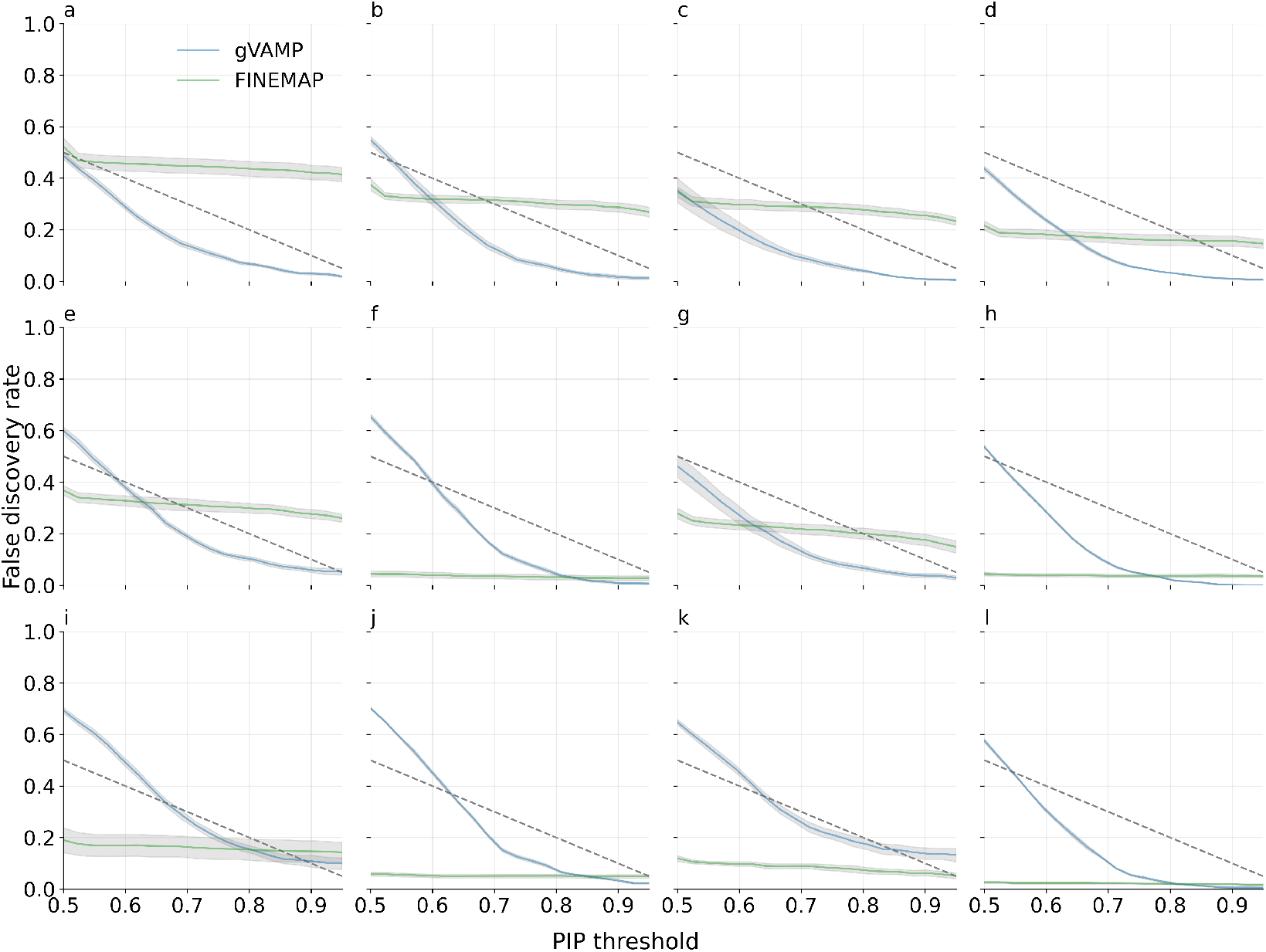
Comparison of the posterior inclusion probability threshold (PIP threshold, *x*-axis) versus the false discovery rate (FDR, *y*-axis) between gVAMP and marginal GWAS association testing plus fine-mapping implemented in the FINEMAP algorithm across 12 simulation scenarios using whole genome sequence (WGS) data of chromosomes 11 to 15 in the UK Biobank. For the detailed description of different simulation settings, see Methods and Supplementary Table S1. We use a Bonferroni corrected p-value significance threshold for marginal GWAS testing and then having identified associated regions we examine the ability of FINEMAP (green) to conduct variable selection across PIP thresholds. For gVAMP (blue), variable selection is conducted from PIP values obtained from a single model run genome-wide. Error bands give the standard deviation across 5 simulation replicates. Dashed lines gives the maximum FDR expected at different PIP thresholds for a model that correctly conducts variable selection based on PIP values.

**Figure S3.**
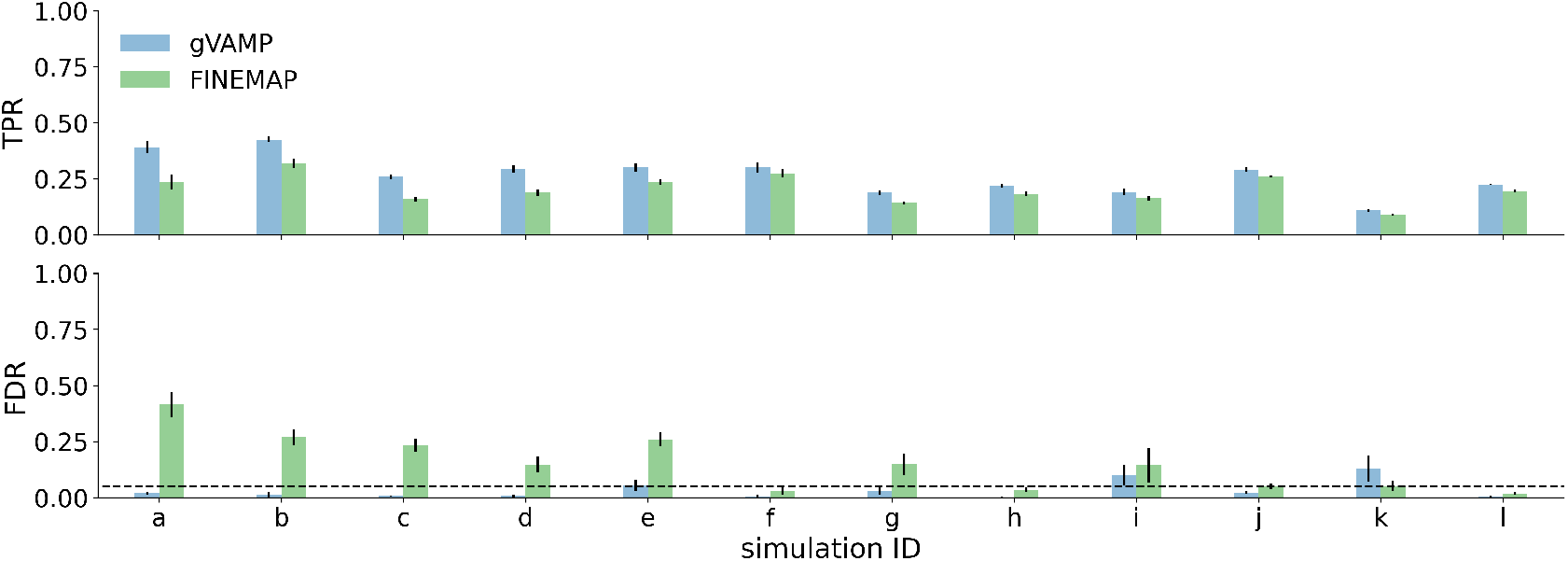
Performance in terms of the true positive rate (TPR) and the false discovery rate (FDR) of marginal GWAS testing plus fine-mapping with FINEMAP, and gVAMP at posterior inclusion probability ≥ 0.95 in a whole genome sequence (WGS) data simulation study of chromosomes 11 to 15 in the UK Biobank. We use a Bonferroni corrected p-value significance threshold for marginal GWAS and PIP > 0.95 for FINEMAP (green) applied for regions identified as significant in the marginal analysis. FINEMAP controls the FDR in only 5 out of 12 WGS data simulation settings (see Methods and Supplementary Table S1). In contrast, using a PIP > 0.95 threshold for the gVAMP testing (blue) controls FDR in 10 out of 12 settings, whilst yielding consistently higher TPR across all settings. Error bars give the standard deviation across 5 simulation replicates.

**Figure S4.**
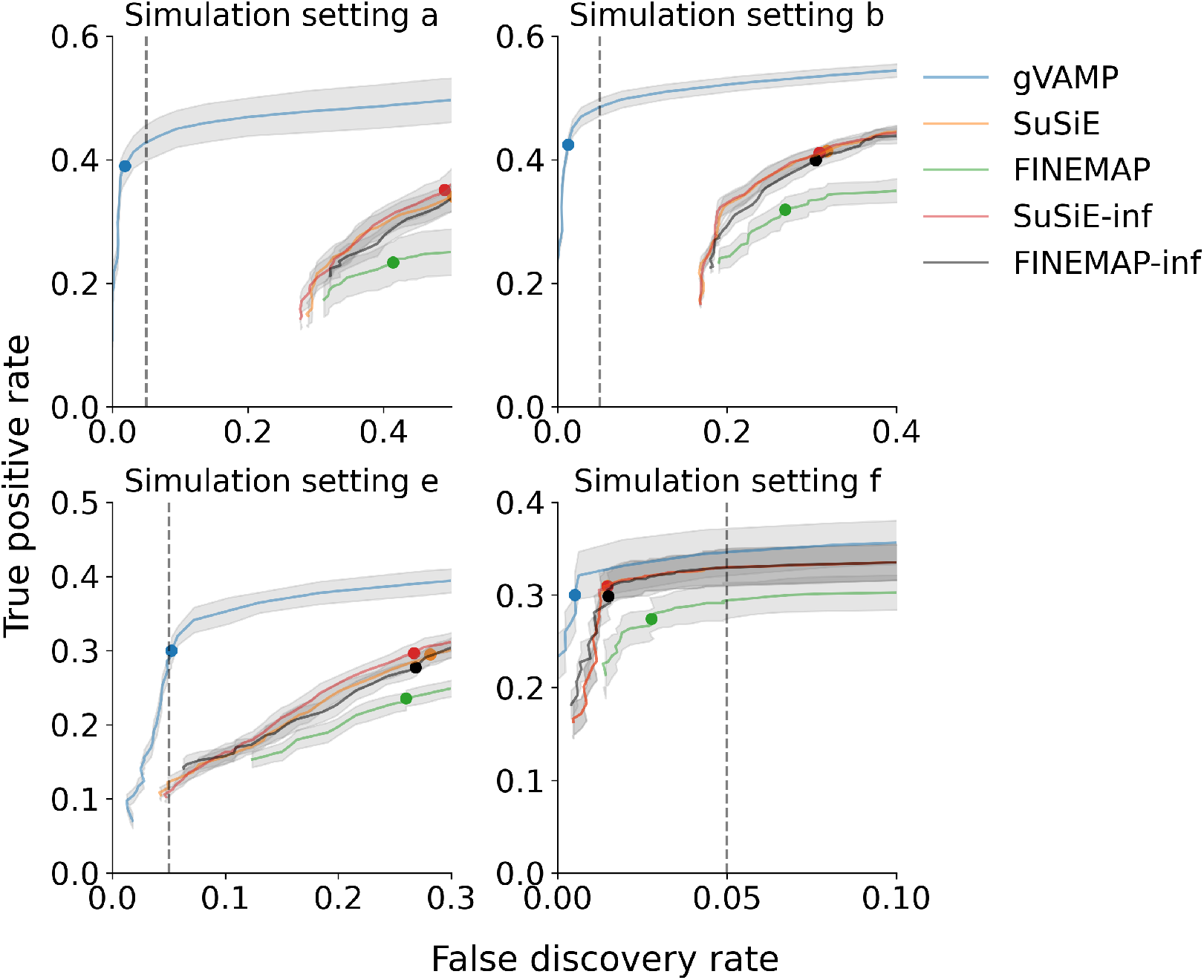
Benchmarking of gVAMP to a wide range of fine-mapping methods applied to marginal GWAS summary data in a whole genome sequence (WGS) data simulation study of chromosomes 11 to 15 in the UK Biobank. For the detailed description of different simulation settings, see Methods and Supplementary Table S1. We use a Bonferroni corrected p-value significance threshold for marginal GWAS testing and then having identified associated regions we apply FINEMAP (green) and SuSiE (orange) to conduct variable selection across PIP thresholds. We also compare to SuSiE-inf (red) and FINEMAP-inf (black) run genome-wide. We compare the true positive rate (TPR) and false discovery rate (FDR) of the methods, with coloured dots indicating the TPR and FDR at ≥ 0.95 PIP, lines giving the TPR and FDR across PIP thresholds and error bands giving the standard deviation across 5 simulation replicates. SuSiE-inf and FINEMAP-inf improve the TPR, but generally not the FDR. gVAMP gives consistently higher TPR at FDR ≤ 5%.

**Figure S5.**
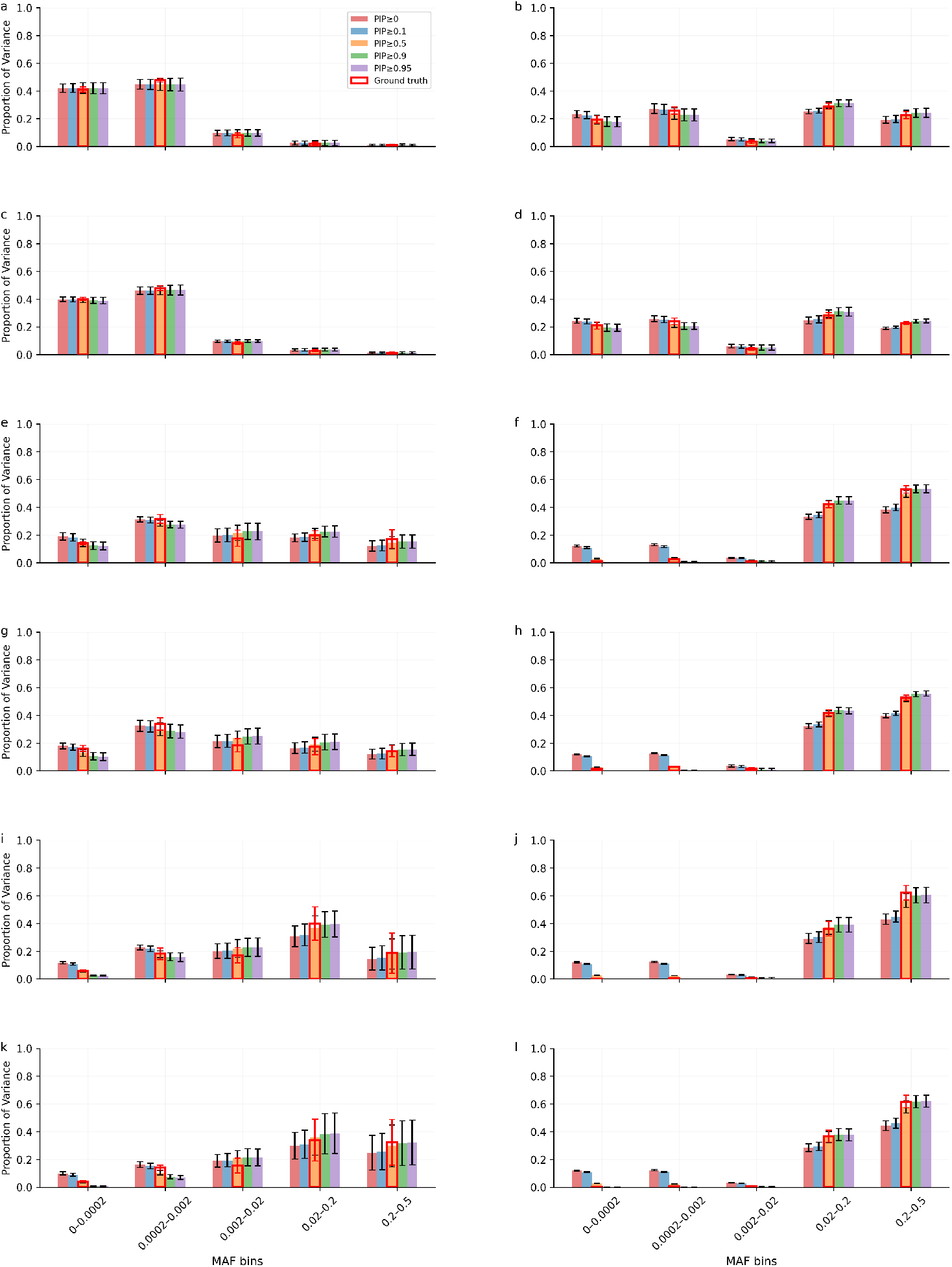
The proportion of variance attributable to DNA loci of different frequency in a whole genome sequence (WGS) data simulation study using chromosomes 11 to 15 of the UK Biobank. We plot the proportion of the sum of the squared regression coefficients attributable to variants of different frequency as estimated from gVAMP across different posterior inclusion probability (PIP) thresholds. The ground truth (red frame) is calculated as the sum of squared true effects across different frequency groups. The variance attributable to SNPs of different frequency matches the ground truth when selecting variants at posterior inclusion probability ≥ 0.5. Error bars give the standard deviation across 5 simulation replicates.

**Figure S6.**
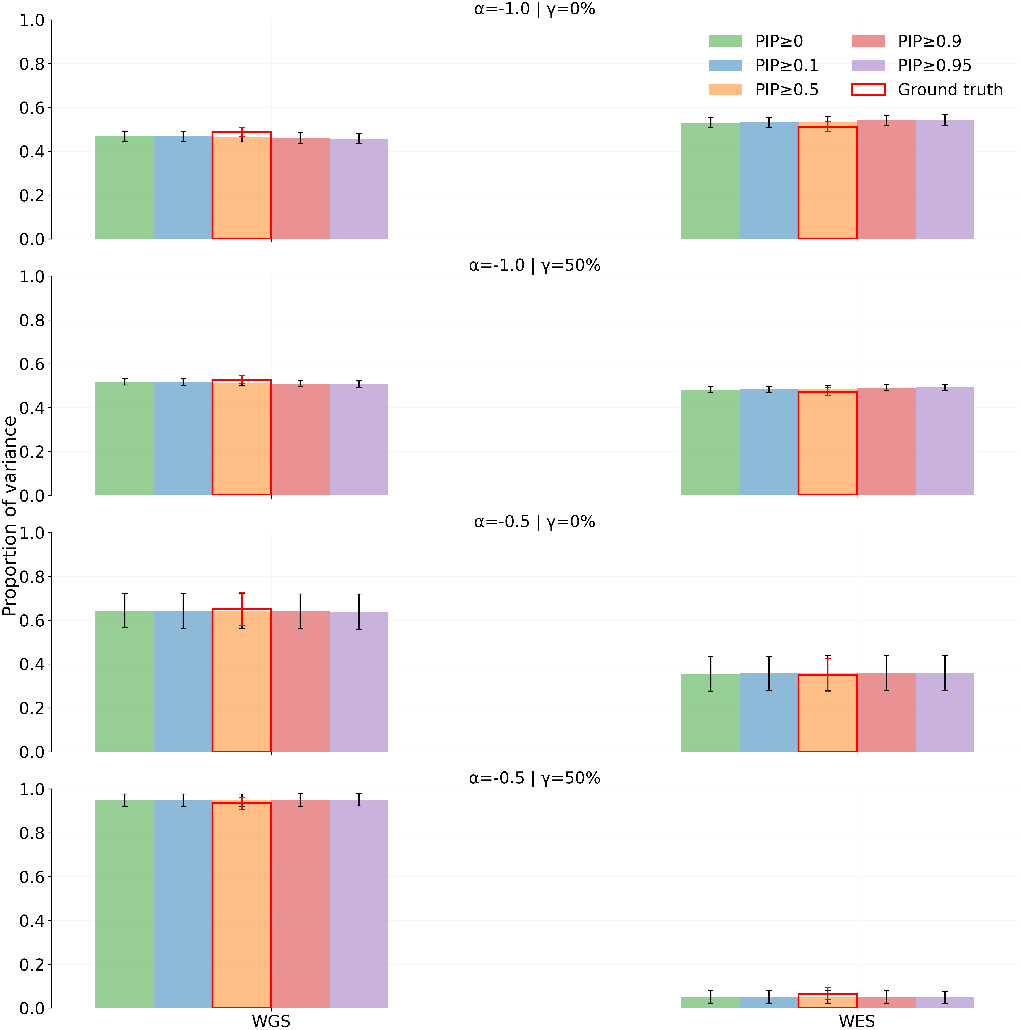
Simulation study including WES burden scores alongside WGS variants from chromosomes 11 to 15 of the UK Biobank. We simulate 500 underlying causal signals with a total heritability of *h*^2^ = 7.25%, allocating 75% of signals to WGS variants and 25% to WES burden scores, with WES burden scores exhibiting effect sizes three-fold larger than those of individual WGS variants. For the detailed description of *α* and *γ* parameters, see Methods. We find that the variance explained by WES and WGS, as inferred by gVAMP, aligns well with the ground truth across a range of PIP thresholds.

**Figure S7.**
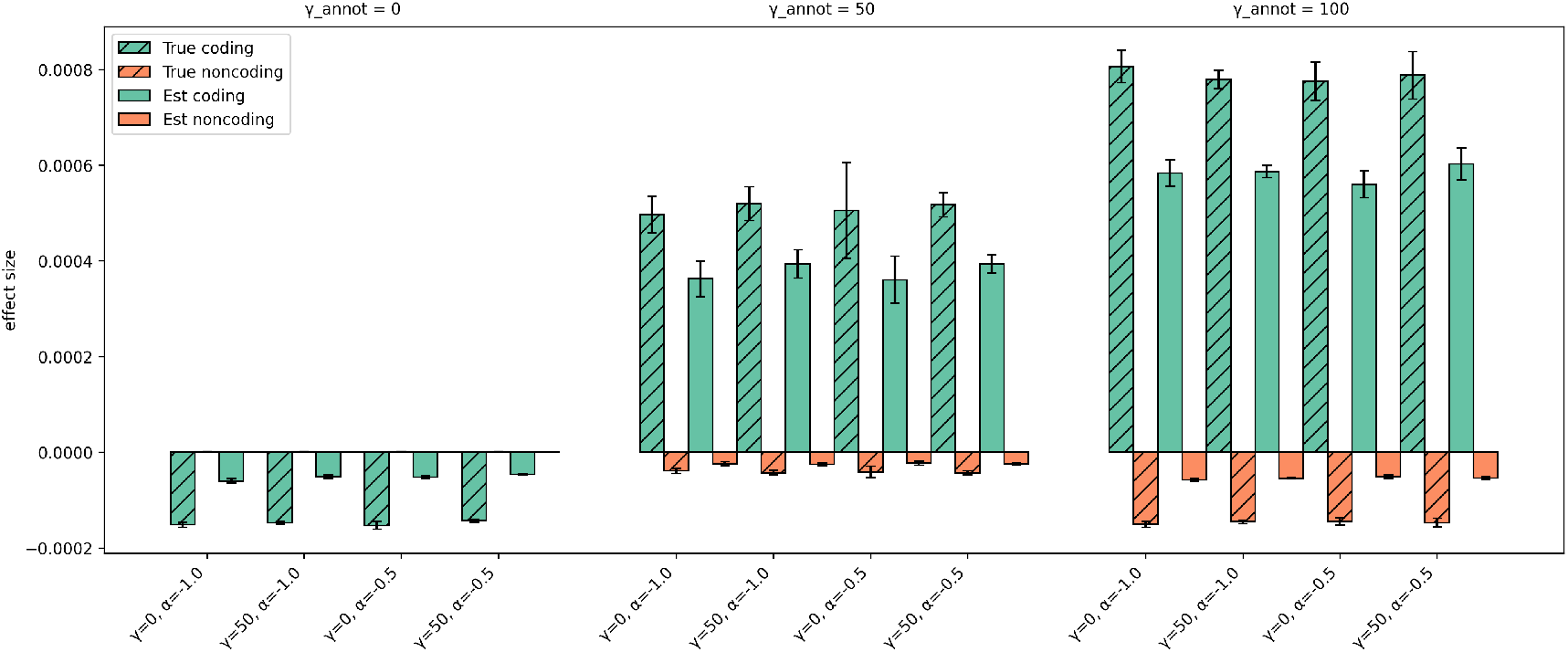
Simulation study stratifying causal effects to coding and non-coding regions. We simulate 500 signals with a total heritability of *h*^2^ = 7.25% across a variety of settings for power law *α* and percentage of common variants *γ*, also allocating *γ*_annot_ ∈ { 0, 50, 100} % percentage of causal variants to the coding region of the DNA. Effect sizes (y-axis) are calculated as the difference of the group average absolute value of the effects from the total group average absolute value of the effect (enrichment). The true simulated values are given by boxes with diagonal lines.

**Figure S8.**
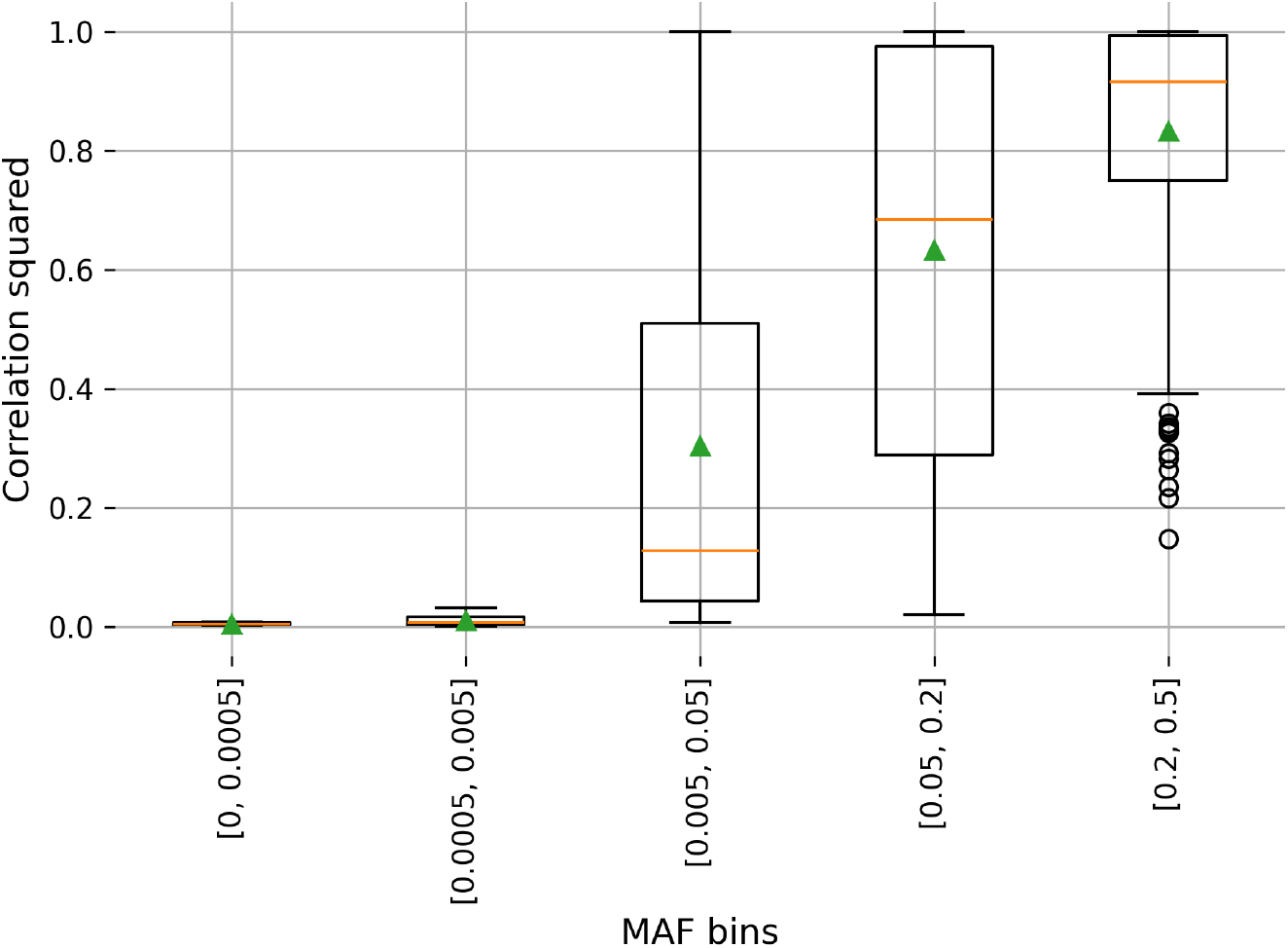
Distribution of the maximum squared correlation between each height-associated variant found by gVAMP and variants reported by Yengo *et al*. [1] within a 1Mb window across MAF groups. We estimate the maximum correlation within the UK Biobank WGS data of each gVAMP discovery at PIP≥ 95% with Yengo *et al*. SNPs significant at *p* ≤ 5 · 10^−8^ within a 1 Mb window and stratify the squared correlations by the MAF of the gVAMP discovered variant. The orange line depicts the median, while the green triangle indicates the mean value. In total, we find 432 SNPs with correlation squared greater than 0.1. The results suggest that very rare variants (MAF ≤ 0.005) are independent of the colocalizing common variants from [1].

**Figure S9.**
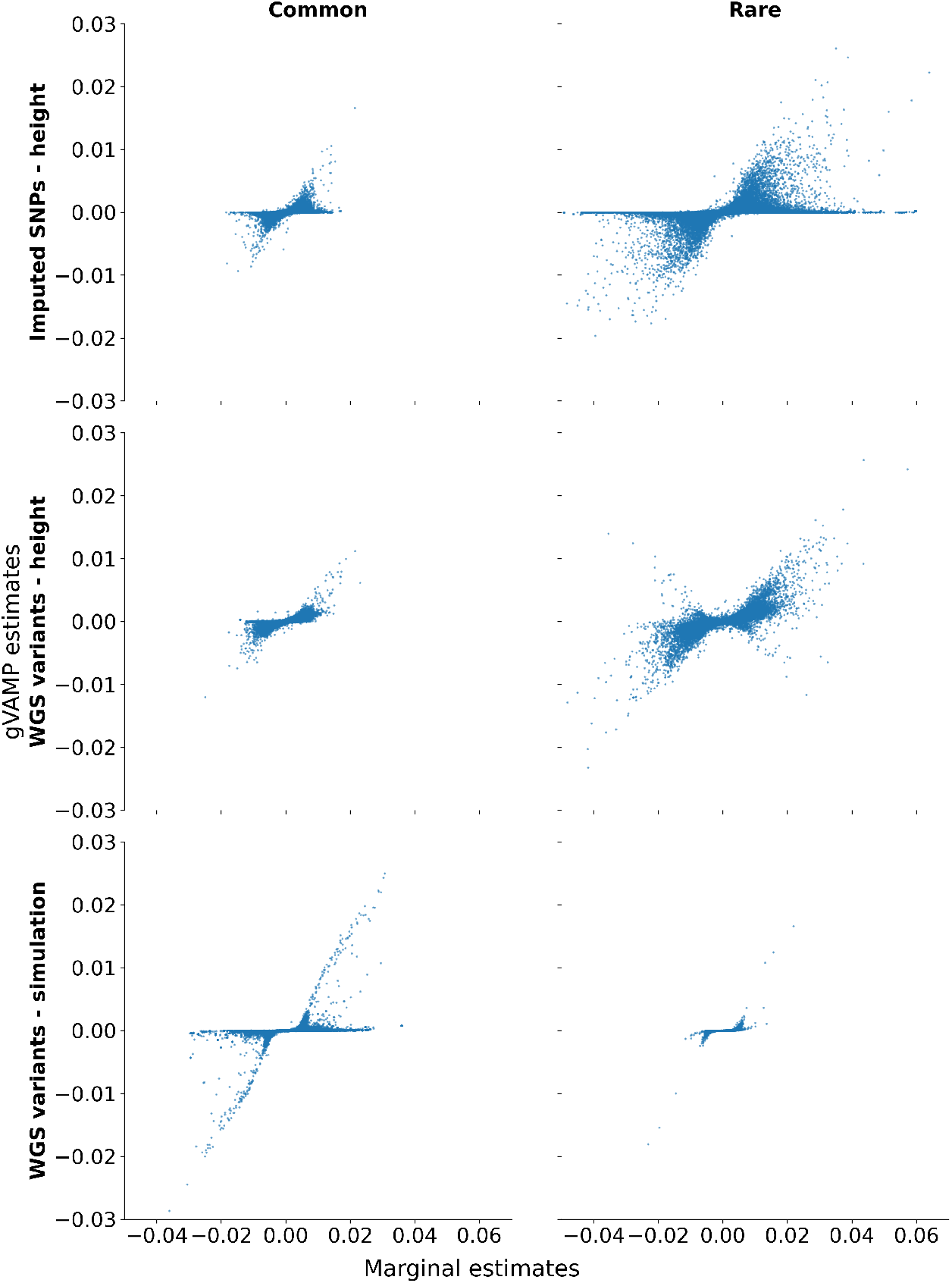
Marginal regression coefficient estimates of the relationship of 16,854,878 whole genome sequence (WGS) variants with human height, plotted against the regression coefficient values obtained from gVAMP. “WGS” refers to the gVAMP run on the WGS variants, “Imputed” refers to the gVAMP run on 8,430,446 imputed SNP markers, and “Simulation” gives the same comparison of marginal and gVAMP estimates in setting *a*. The comparison is plotted for common (minor allele frequency, MAF ≥ 1%) and rare variants (MAF < 1%).

**Figure S10.**
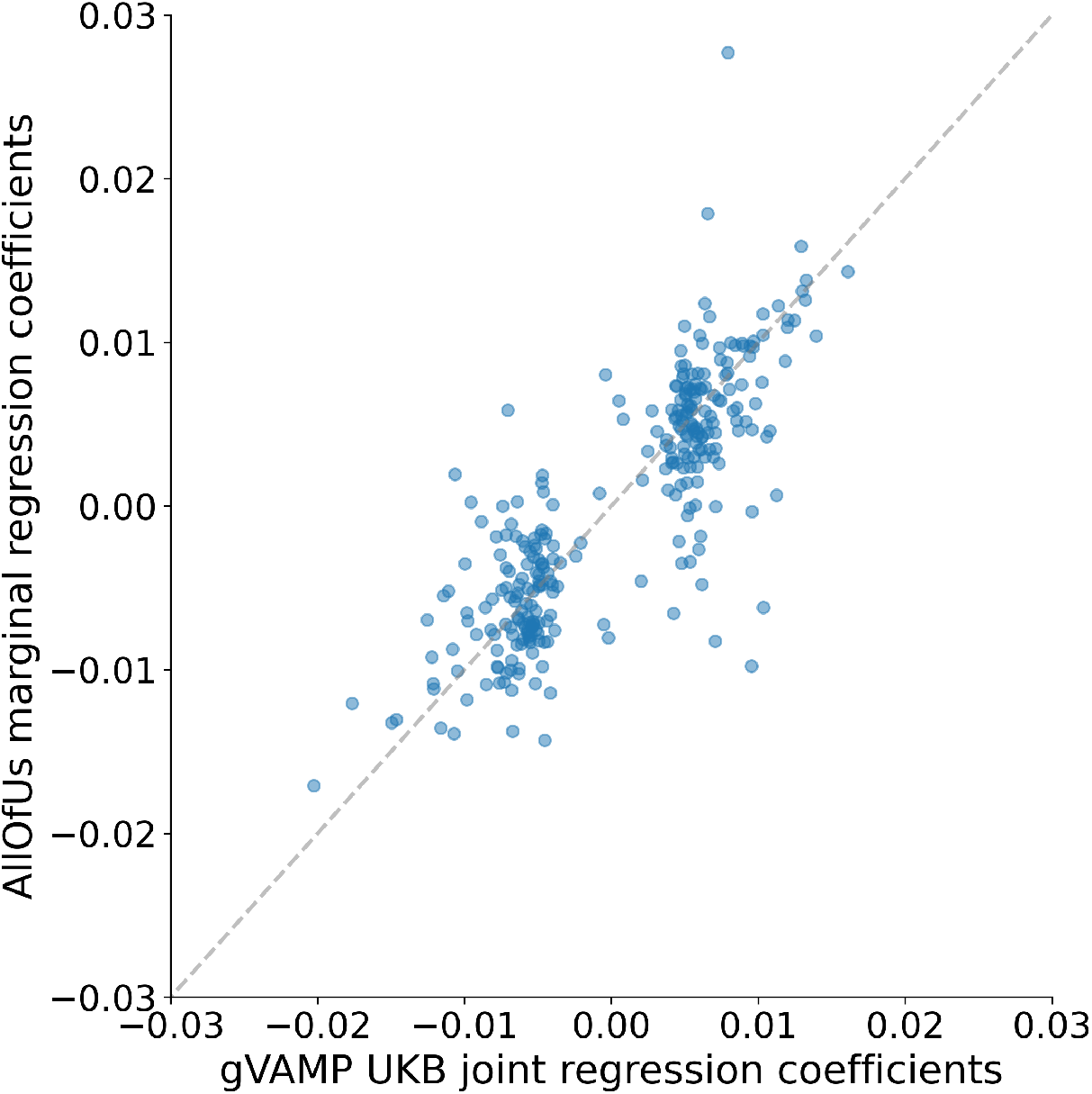
Replication of UK Biobank gVAMP findings in the All of Us dataset. On the x-axis, we plot gVAMP regression coefficient estimates for human height in WGS data discovered at PIP≥ 0.95. On the y-axis, we plot the marginal regression coefficient estimates of the same SNPs in WGS data in the All of Us dataset, controlling for sex and 16 available PCs. When the major and minor alleles are swapped between two datasets, we change the sign of the effect to match them. The slope of the fitted regression line is 0.8471.

**Figure S11.**
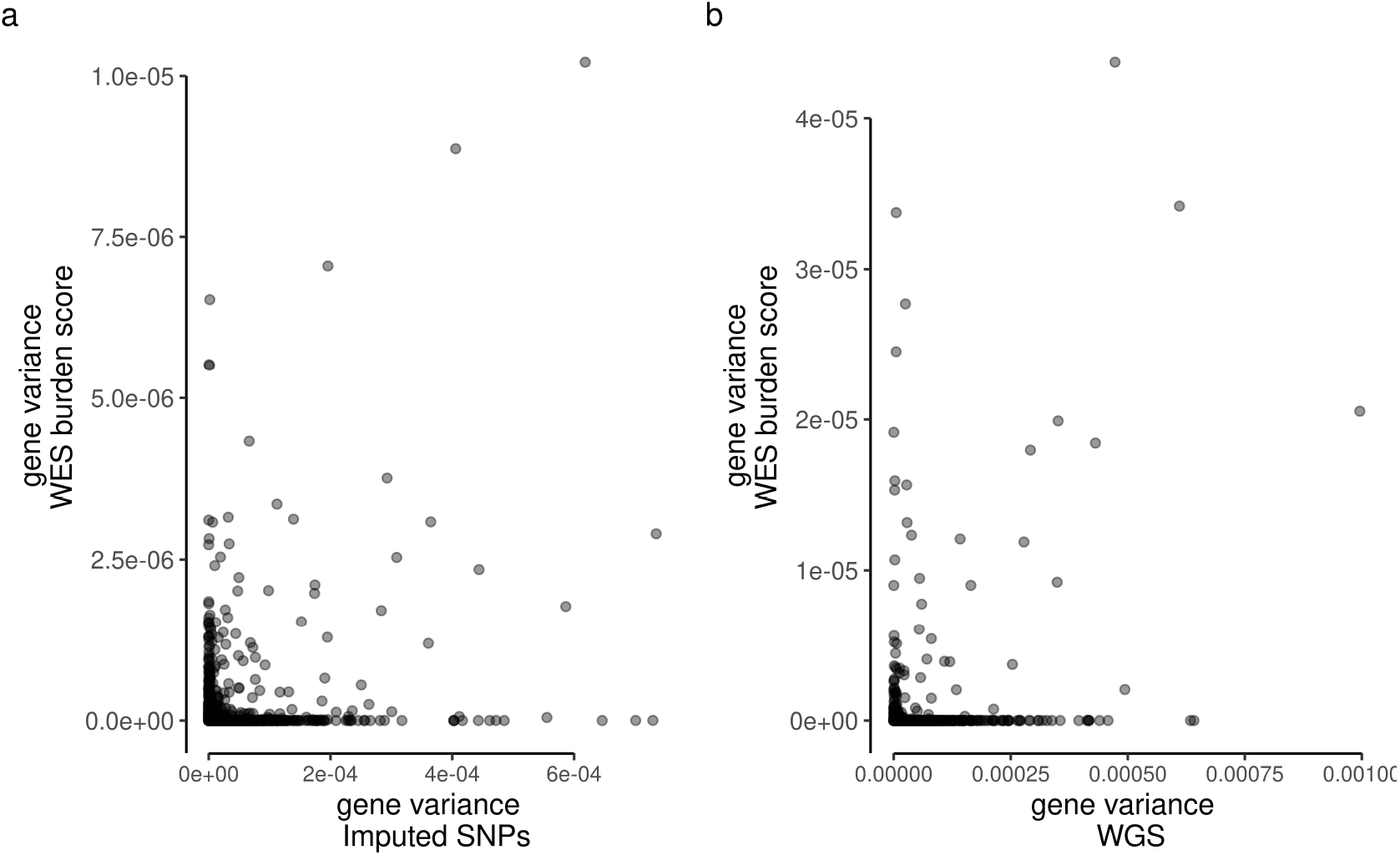
The variance attributable to gene burden scores calculated from whole exome sequence (WES) data (*y*-axis) shows no relationship to the variance attributable to DNA variants within and around the gene (*x*-axis) estimated from a gVAMP analysis of either 8,430,446 imputed SNP markers or 16,854,878 whole genome sequence (WGS) variants.

**Figure S12.**
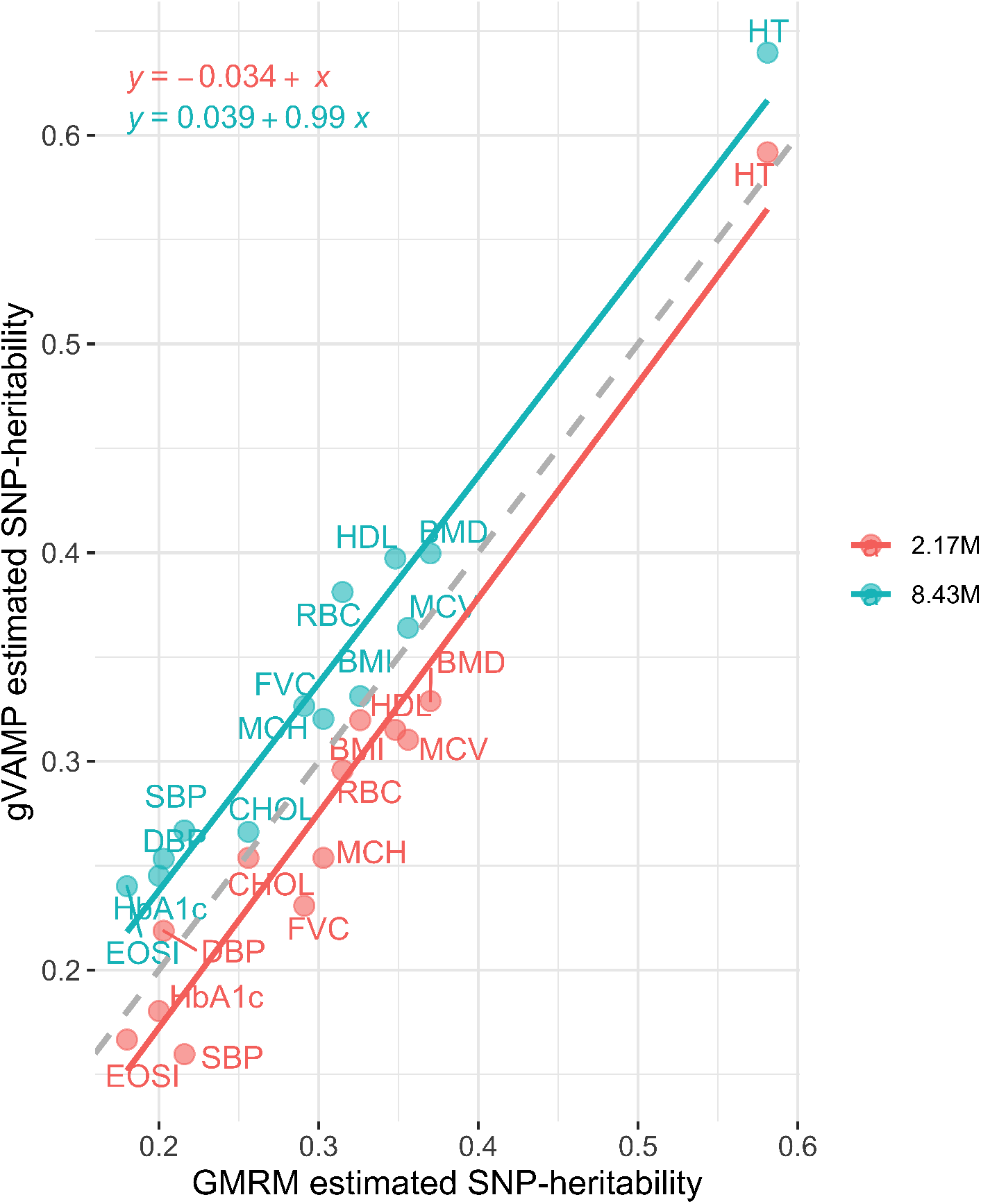
SNP heritability estimation of GMRM versus gVAMP with different numbers of SNP markers across 13 traits in the UK Biobank. Comparison of the proportion of phenotypic variation attributable to 2,174,071 autosomal SNP genetic markers (SNP heritability) estimated by GMRM (*x*-axis) to the SNP heritability estimated by gVAMP (*y*-axis) at either the same 2,174,071 SNPs (red) or 8,430,446 SNP markers (blue). The slope of the lines shows a 1-to-1 relationship of gVAMP to GMRM, but with an average of 3.4% lower estimate for gVAMP at 2.17M SNPs. Analyzing 8.4M SNPs with gVAMP increases the heritability estimate over GMRM by 3.9%, which is consistent with an increase in phenotypic variance captured by the full imputed sequence data, as opposed to a selected subset of SNP markers. The dashed grey line gives *y* = *x*.

## Supplementary Note 1

**Figure S13.**
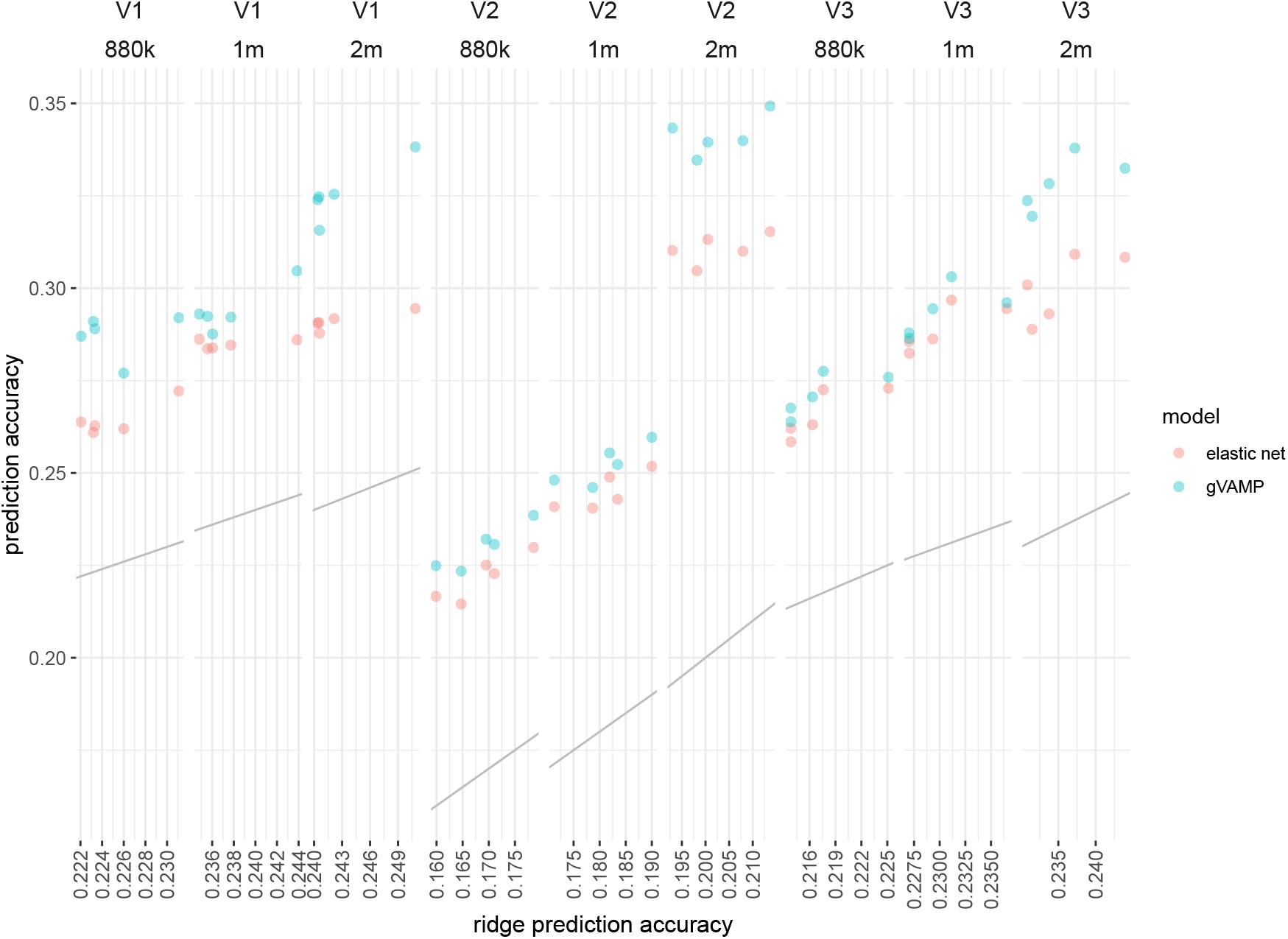
Simulation study of out-of-sample prediction accuracy in UK Biobank imputed genotype data. We compare out-of-sample prediction accuracy (*y*-axis) for polygenic risk scores (PRS) created at different sets of markers (“880k” = 887,060, “1m” = 1,410,525, “2m” = 2,174,071 SNPs) from gVAMP (blue) and an elastic net model implemented in the LDAK software (red), as compared to the prediction accuracy obtained at the same sets of SNPs from a ridge regression model implemented in the LDAK software (*x*-axis). Grey lines give the 1:1 line. Comparisons are made across three settings: “V1” where 40,000 causal variants are randomly selected from 8,430,446 SNP markers and effects are drawn from a Gaussian, “V2” where 40,000 causal variants are randomly selected from 8,430,446 SNP markers, with effects that have a power relationship to the minor allele frequency of -0.5, and “V3” where 40,000 causal variants are randomly selected from 8,430,446 SNP markers, with effects drawn from an elastic net distribution.

### Simulation study in imputed SNP data

To support our empirical analyses we also conduct a simulation study in imputed SNP data using the 8,430,446 UK Biobank genetic marker data with 414,055 individuals. Our WGS simulation focused on variable selection of exact SNP positions and thus to give power to select variables and to enable clear comparisons of fine-mapping methods (where the objective was the selection of a single variable out of a segment) we simulated only 500 or 1000 causal variables on chromosomes 11-15. Here, we sought to simulate the other extreme of a very highly polygenic genetic basis and thus we randomly sample 40,000 causal variants genome-wide (effectively 4 times the density of causal variants simulated in the WGS simulation). We then simulate three scenarios. First, in *V1*, we allocate effect sizes from a Gaussian with mean zero and variance 0.5/40000, where 0.5 is the proportion of variance attributable to the SNP markers. Multiplying the simulated SNP effects by normalized values of the 40,000 causal markers, gives a vector of genetic values of length *N* = 414055 with variance 0.5. To this we add a vector of noise, drawn from a Gaussian with mean zero and variance 0.5, to produce a response variable of length *N*, with zero mean and unit variance. Second, in *V2*, we simulate an *α*-power relationship between the causal variant effect size and the minor allele frequency of *α* = −0.5 (see Methods). Third, in *V3*, we simulate under an elastic net prior, where with probability 0.8 we sample effect sizes from a standard Gaussian, with probability 0.2 we sample from a double exponential distribution, and we then scale the effects vector so that multiplying them with normalized values of the 40,000 causal markers, gives a vector of genetic values of length *N* = 414055 with variance 0.5.

**Figure S14.**
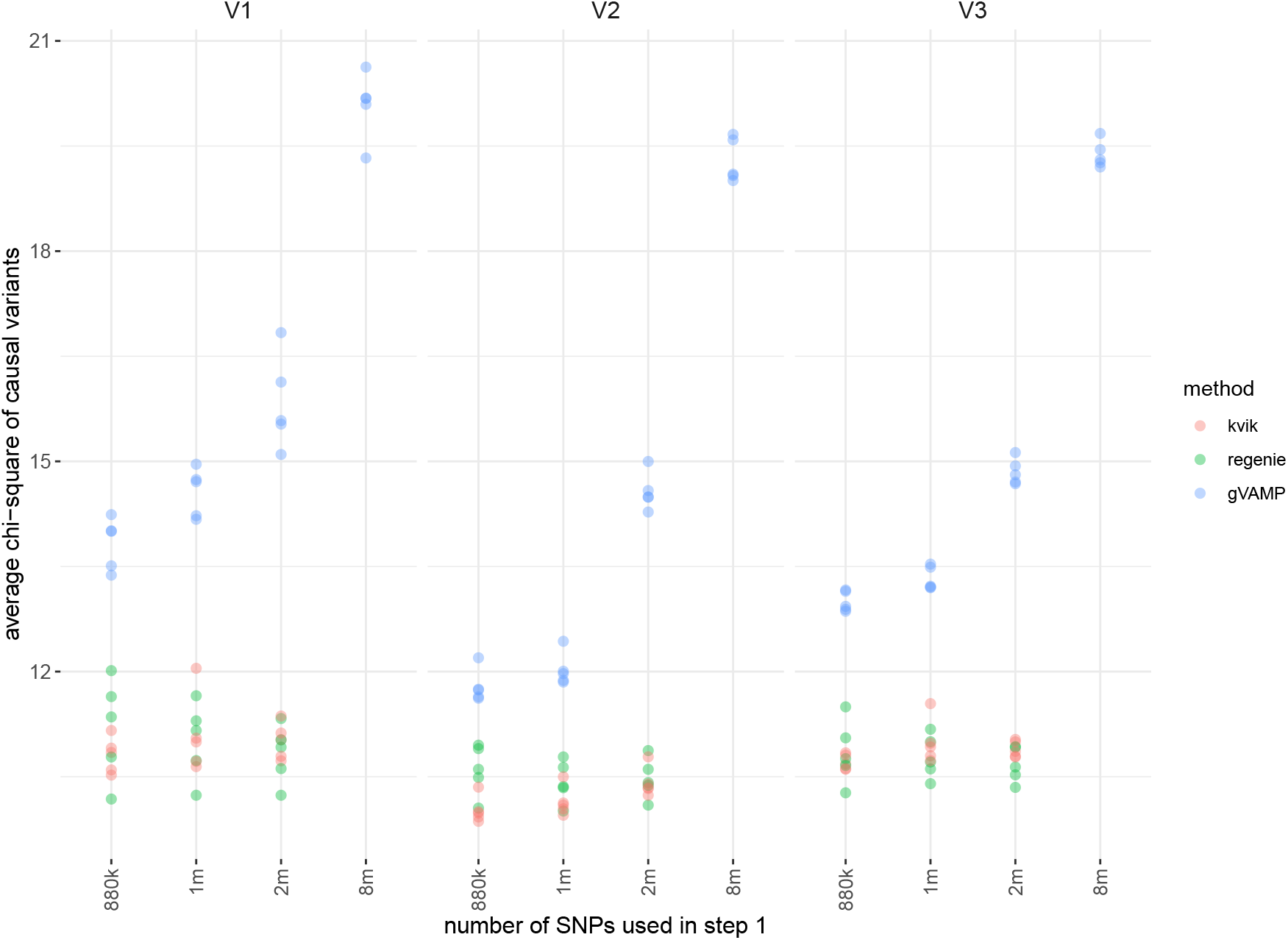
Simulation study of power for mixed linear model leave-one-chromosome out association testing in UK Biobank imputed genotype data. We compare the average chi-squared value at the causal variants (*y*-axis) for leave-one-chromosome-out association testing (LOCO) when the first step LOCO predictors are created at different sets of markers (“880k” = 887,060, “1m” = 1,410,525, “2m” = 2,174,071 SNPs) from gVAMP (blue), an elastic net model implemented in the LDAK software (red), and Regenie (green). Comparisons are made across three settings: “V1” where 40,000 causal variants are randomly selected from 8,430,446 SNP markers and effects are drawn from a Gaussian, “V2” where 40,000 causal variants are randomly selected from 8,430,446 SNP markers, with effects that have a power relationship to the minor allele frequency of -0.5, and “V3” where 40,000 causal variants are randomly selected from 8,430,446 SNP markers, with effects drawn from an elastic net distribution.

**Figure S15.**
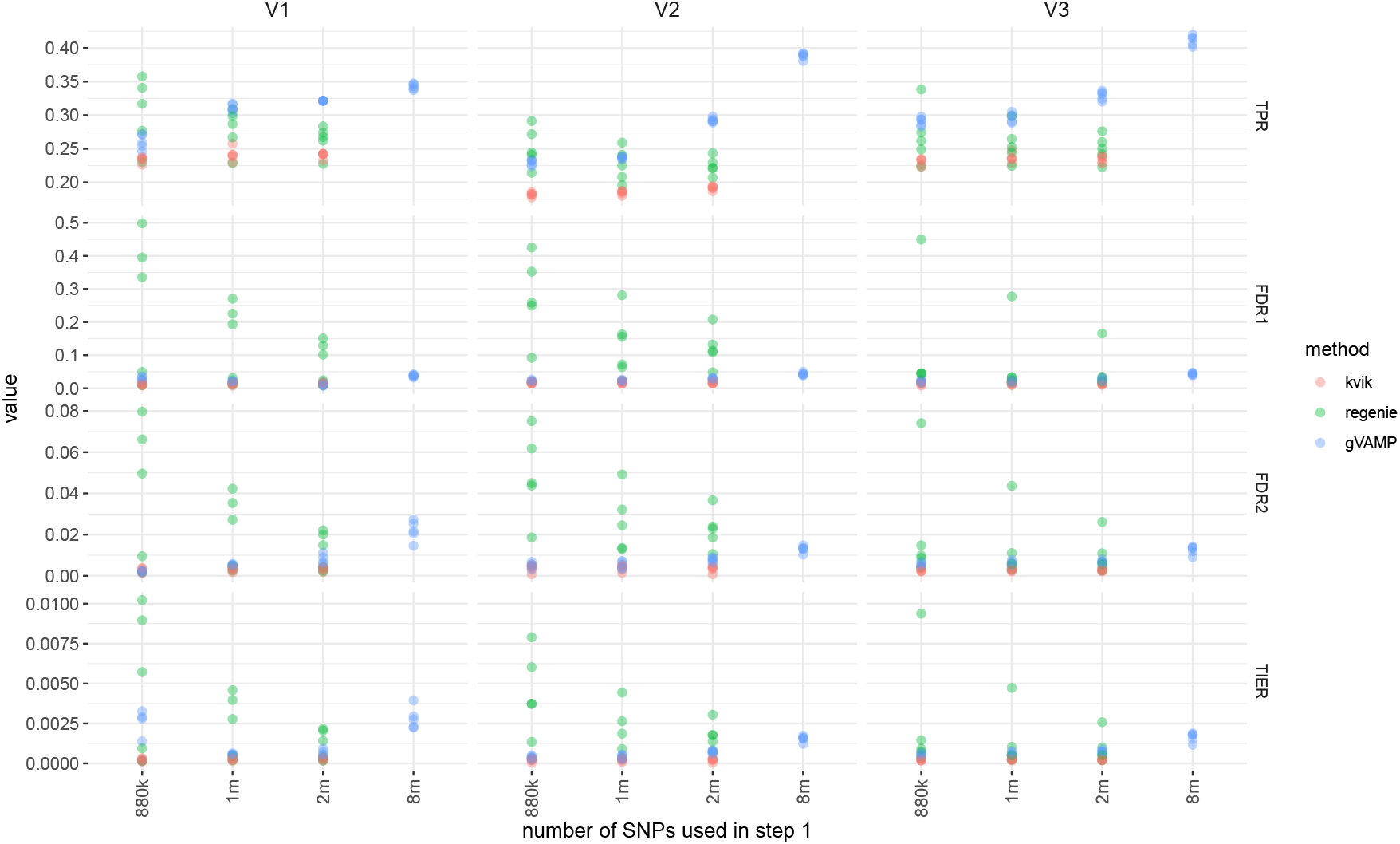
Simulation study of true positive rate (TPR), false discovery rate (FDR) and type-I error for mixed linear model leave-one-chromosome out association testing in UK Biobank imputed genotype data. We compare performance of leave-one-chromosome-out association testing (LOCO) when the first step LOCO predictors are created at different sets of markers (“880k” = 887,060, “1m” = 1,410,525, “2m” = 2,174,071 SNPs) from gVAMP (blue), an elastic net model implemented in the LDAK software (red), and Regenie (green). Comparisons are made across three settings: “V1” where 40,000 causal variants are randomly selected from 8,430,446 SNP markers and effects are drawn from a Gaussian, “V2” where 40,000 causal variants are randomly selected from 8,430,446 SNP markers, with effects that have a power relationship to the minor allele frequency of -0.5, and “V3” where 40,000 causal variants are randomly selected from 8,430,446 SNP markers, with effects drawn from an elastic net distribution. For each 1Mb window that contains an identified genome-wide significant association, we then ask if the identified variants are correlated (squared correlation ≥ 0.01) to a causal variant: if so, we classify the window as a true positive; otherwise, we classify it as a false positive. The true positive rate is calculated as the number of true positive 1Mb windows divided by the total number of simulated causal variants, and it is also known as the recall, or sensitivity, reflecting the power of a statistical test. We provide two measures of the false discovery rate: FDR1 is calculated as the number of false positive 1Mb windows divided by the number of 1Mb windows that contain genome-wide significant associations, and it is a measure of the proportion of independent discoveries that are false. FDR2 is calculated as the total number of false positive SNPs (squared correlation ≤ 0.01 with a causal variant) identified divided by the total number of identified genome-wide significant associations. We calculate the type-I error rate (TIER) as the number of SNPs with no correlation to a causal variant (squared correlation ≤ 0.01 with a causal variant) that pass the Bonferroni genome-wide significance testing threshold divided by the total number of SNPs that have no correlation to a causal variant (squared correlation ≤ 0.01 with a causal variant).

We analyze the simulated response variable with gVAMP, using either 8,430,446, 2,174,071, 1,410,525 or 887,060 SNP markers with identical initialization to that described in the Methods for the empirical UK Biobank study. We also analyze the data with REGENIE using 2,174,071, 1,410,525 or 887,060 SNP markers for the first stage and 8,430,446 SNP markers for the second stage 887,060 SNP markers for the first stage and 8,430,446 SNP markers for the second stage LOCO testing. This creates three simulation scenarios where the heritability and number of causal variants is fixed, but the effect size distributions vary. We note that because the 2,174,071, 1,410,525 or 887,060 SNP marker sets are LD reduced subsets of the larger 8,430,446 SNP markers, the simulated causal variants are in LD with markers used in step 1, but they may not be present within the data. This is in contrast to the simulations presented in the LDAK-KVIK[2] and REGENIE[3] papers, where the causal variants are always within the subset of SNPs used in the first stage of the analysis.

We begin by comparing the out-of-sample prediction accuracy obtained by each approach for the different marker sets, following the same procedures outlined in the Methods for the empirical UK Biobank analysis. Note that it is not possible to obtain SNP marker effects for the model used in LDAK software. For the out-of-sample prediction, we use a hold-out set of 15,000 individuals that are unrelated (SNP marker relatedness < 0.05) to the training individuals. We present these results in Figure S13.

**Figure S16.**
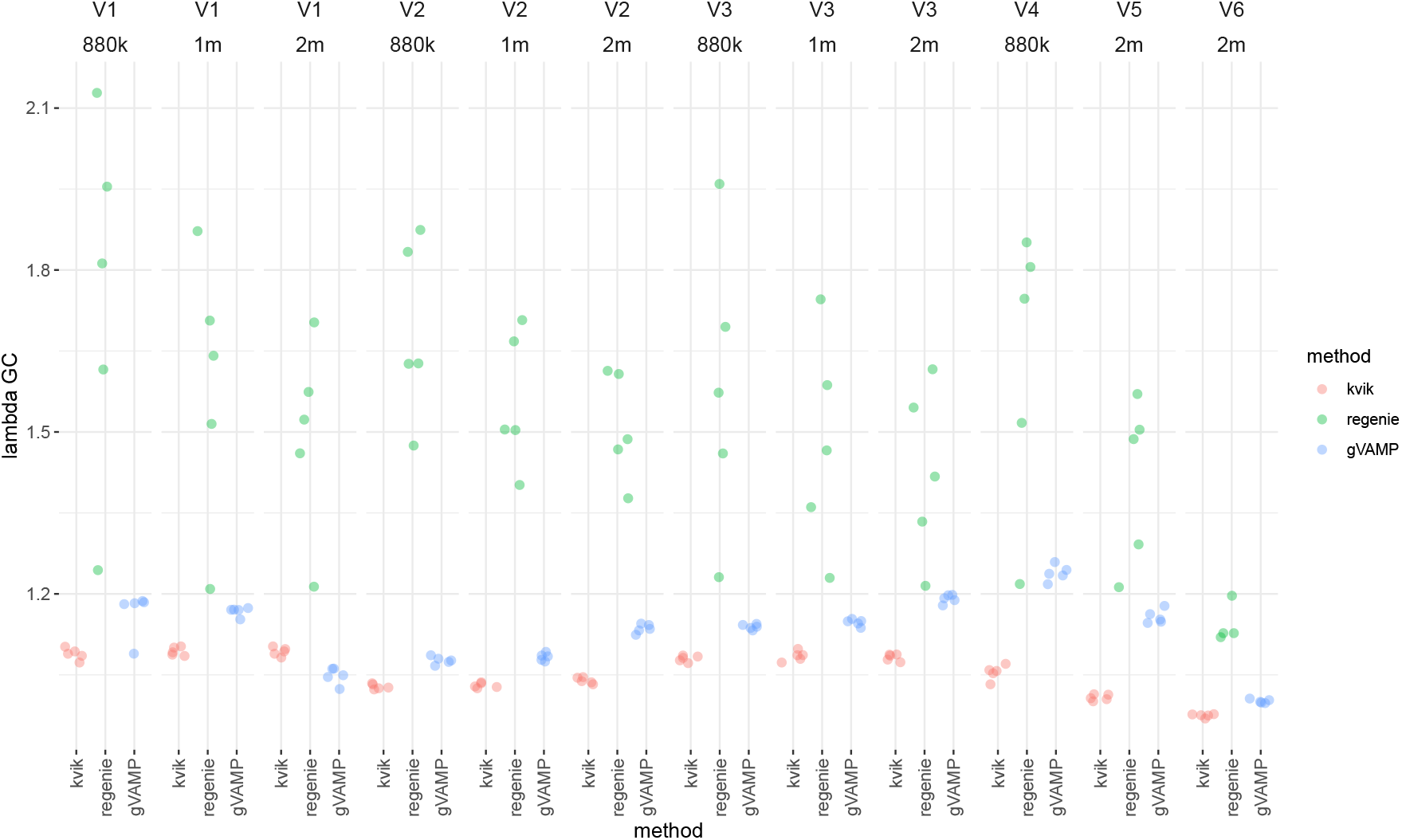
Simulation study of lambda GC (*λ*_*GC*_) values of mixed linear model leave-one-chromosome out association testing in UK Biobank imputed genotype data. We compare performance of leave-one-chromosome-out association testing (LOCO) when the first step LOCO predictors are created at different sets of markers, from gVAMP (blue), an elastic net model implemented in the LDAK software (red), and Regenie (green). Comparisons are made across six settings: “V1” where 40,000 causal variants are randomly selected from 8,430,446 SNP markers and effects are drawn from a Gaussian, “V2” where 40,000 causal variants are randomly selected from 8,430,446 SNP markers, with effects that have a power relationship to the minor allele frequency of -0.5, “V3” where 40,000 causal variants are randomly selected from 8,430,446 SNP markers, with effects drawn from an elastic net distribution, “V4” where 40,000 causal variants are randomly selected from 887,060 SNPs, with effects that contribute 50% to the phenotypic variance and have a power relationship to the minor allele frequency of -0.5, “V5” where 40,000 causal variants are randomly selected from 2,174,071 SNPs, with effects that contribute 50% to the phenotypic variance and have a power relationship to the minor allele frequency of -0.5, and “V6” where 5,000 causal variant effects are randomly selected from 2,174,071 SNPs, with effects that contribute 25% to the phenotypic variance and have a power relationship to the minor allele frequency of -0.5. For each replicate, we calculate the ratio of the median chi-square to the expected median chi-square values (*λ*_*GC*_).

**Figure S17.**
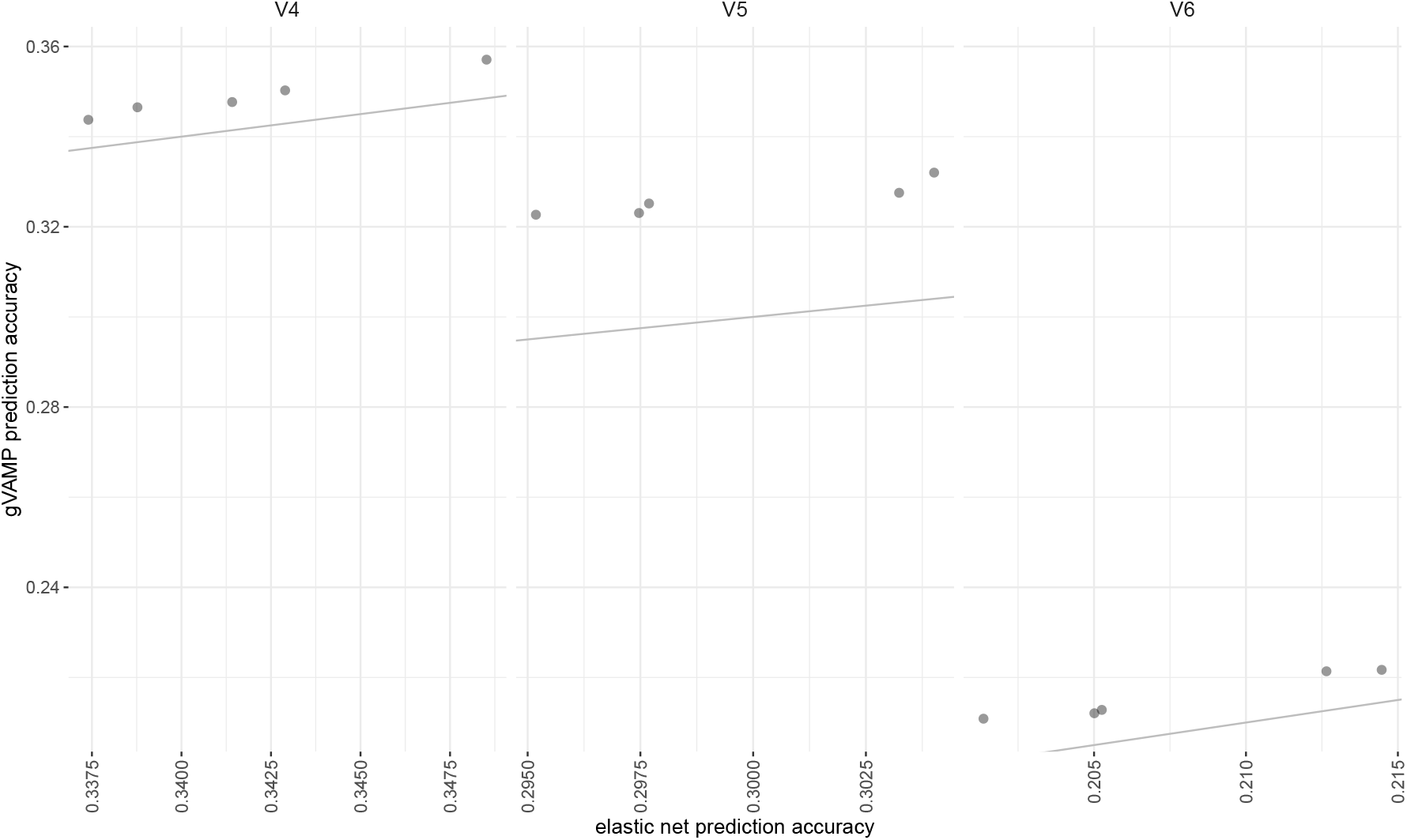
Simulation study of out-of-sample prediction accuracy in UK Biobank imputed genotype data. We compare out-of-sample prediction accuracy (*y*-axis) for polygenic risk scores (PRS) created at different sets of markers from gVAMP and an elastic net model implemented in the LDAK software. Grey lines give the 1:1 line. Comparisons are made across three settings: “V4” where 40,000 causal variants are randomly selected from 887,060 SNPs, with effects that contribute 50% to the phenotypic variance and have a power relationship to the minor allele frequency of -0.5, “V5” where 40,000 causal variants are randomly selected from 2,174,071 SNPs, with effects that contribute 50% to the phenotypic variance and have a power relationship to the minor allele frequency of -0.5, and “V6” where 5,000 causal variant effects are randomly selected from 2,174,071 SNPs, with effects that contribute 25% to the phenotypic variance and have a power relationship to the minor allele frequency of -0.5.

We then compare the LOCO association testing results obtained by REGENIE, LDAK-KVIK, and gVAMP. We estimate the average chi-squared value at the causal variants and present these results in Figure S14. We give a simple, fair comparison of the true positive rate (TPR) and false discovery rate (FDR) across methods controlling for LD, where we calculate the number of independent associations (squared correlation ≤ 0.01) identified by each approach that pass a Bonferroni genome-wide significance testing threshold of 0.05/8430446 within 1Mb windows of the DNA. Specifically, for each 1Mb window that contains an identified genome-wide significant association, we then ask if it is correlated (squared correlation ≥ 0.01) to a causal variant: if so, we classify it as a true positive; otherwise, we classify it as a false positive. The true positive rate is calculated as the number of true positive 1Mb windows divided by the total number of simulated causal variants, and it is also known as the recall, or sensitivity, reflecting the power of a statistical test. We provide two measures of the false discovery rate: FDR1 is calculated as the number of false positive 1Mb windows divided by the number of 1Mb windows that contain genome-wide significant associations, and it is a measure of the proportion of independent discoveries that are false. FDR2 is calculated as the total number of false positive SNPs (squared correlation ≤ 0.01 with a causal variant) identified divided by the total number of identified genome-wide significant associations. As genome-wide association studies aim to detect regions of the DNA associated with the phenotype, the definition of a false discovery as the detection of a variant at genome-wide significance when that variant has squared correlation ≤ 0.01 with a causal variant within 1Mb is a very conservative one. Additionally, we calculate the type-I error rate as the number of SNPs with no correlation to a causal variant (squared correlation ≤ 0.01 with a causal variant) that pass the Bonferroni genome-wide significance testing threshold divided by the total number of SNPs that have no correlation to a causal variant (squared correlation ≤ 0.01 with a causal variant). We present these results in Figure S15. Finally, we also calculate the ratio of the median chi-square to the expected median chi-square values (*λ*_*GC*_) and we present these results in Figure S16. We conduct five simulation replicates for each simulation scenarios, as we find that this is more than sufficient to contrast methods, as the results were very consistent across replicates.

**Figure S18.**
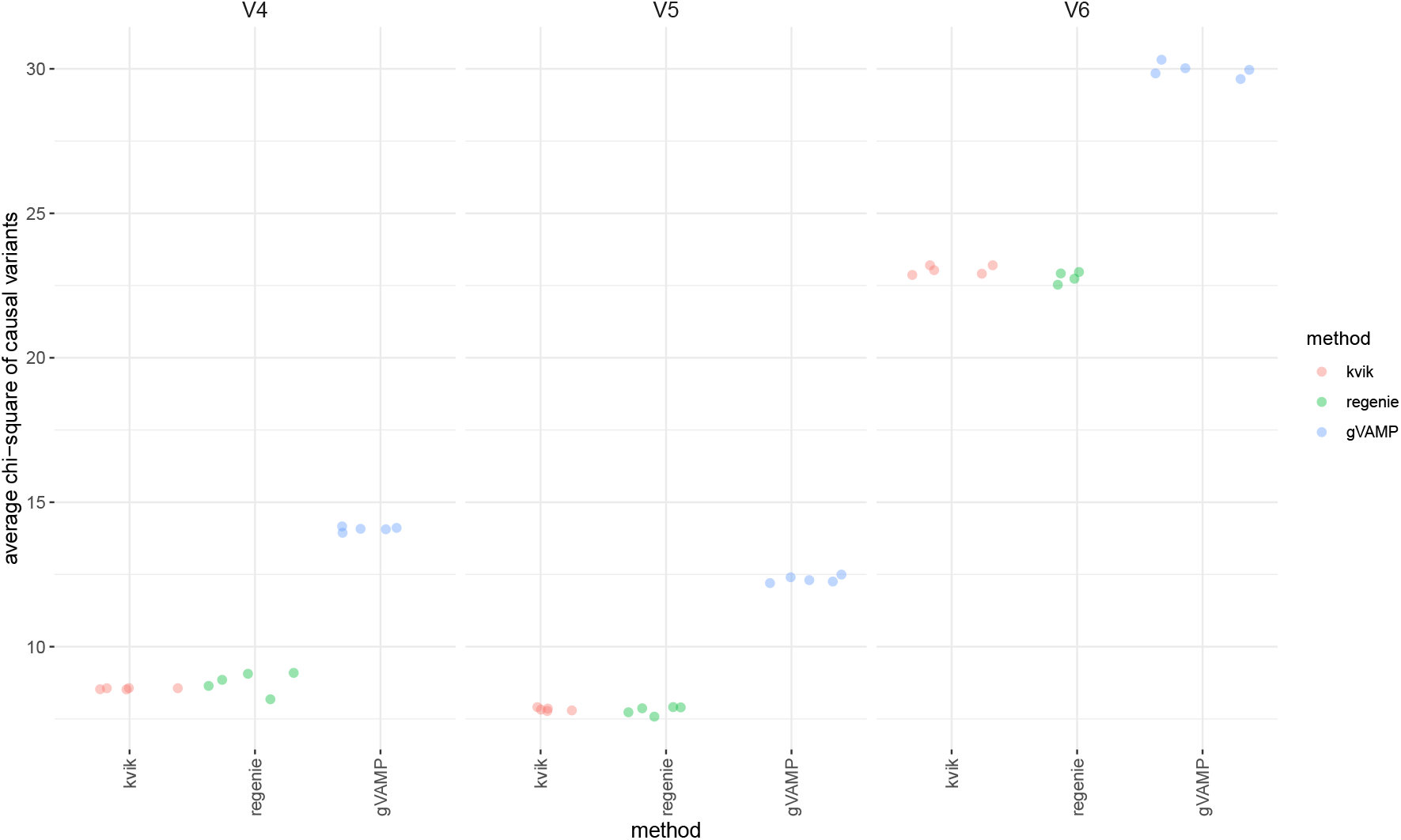
Simulation study of power for mixed linear model leave-one-chromosome out association testing in UK Biobank imputed genotype data. We compare the average chi-squared value at the causal variants (*y*-axis) for leave-one-chromosome-out association testing (LOCO) when the first step LOCO predictors are created at different sets of markers, from gVAMP (blue), an elastic net model implemented in the LDAK software (red), and Regenie (green). Comparisons are made across three settings: “V4” where 40,000 causal variants are randomly selected from 887,060 SNPs, with effects that contribute 50% to the phenotypic variance and have a power relationship to the minor allele frequency of -0.5, “V5” where 40,000 causal variants are randomly selected from 2,174,071 SNPs, with effects that contribute 50% to the phenotypic variance and have a power relationship to the minor allele frequency of -0.5, and “V6” where 5,000 causal variant effects are randomly selected from 2,174,071 SNPs, with effects that contribute 25% to the phenotypic variance and have a power relationship to the minor allele frequency of -0.5.

**Figure S19.**
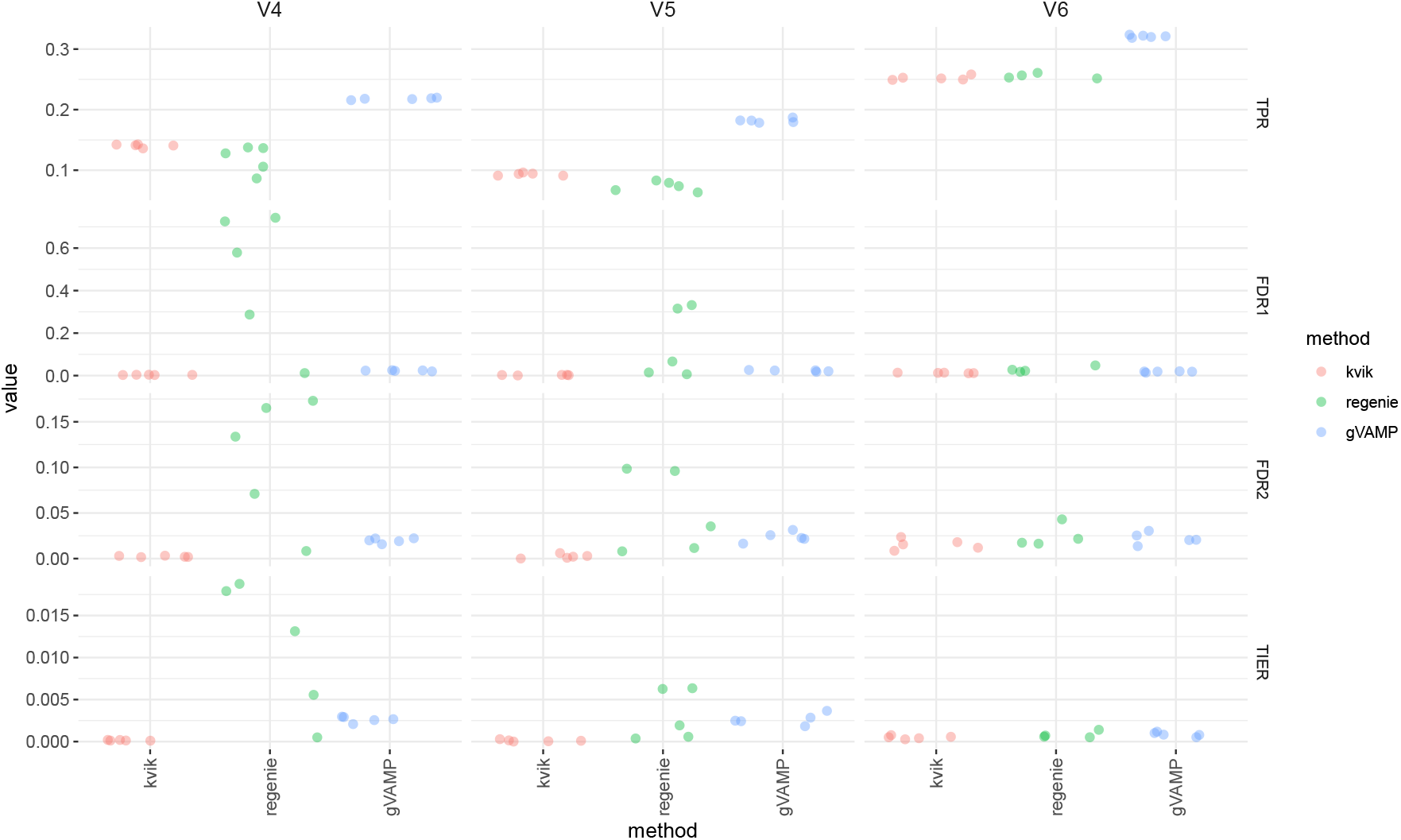
Simulation study of true positive rate (TPR), false discovery rate (FDR) and type-I error for mixed linear model leave-one-chromosome out association testing in UK Biobank imputed genotype data. We compare performance of leave-one-chromosome-out association testing (LOCO) when the first step LOCO predictors are created at different sets of markers, from gVAMP (blue), an elastic net model implemented in the LDAK software (red), and Regenie (green). Comparisons are made across three settings: “V4” where 40,000 causal variants are randomly selected from 887,060 SNPs, with effects that contribute 50% to the phenotypic variance and have a power relationship to the minor allele frequency of -0.5, “V5” where 40,000 causal variants are randomly selected from 2,174,071 SNPs, with effects that contribute 50% to the phenotypic variance and have a power relationship to the minor allele frequency of -0.5, and “V6” where 5,000 causal variant effects are randomly selected from 2,174,071 SNPs, with effects that contribute 25% to the phenotypic variance and have a power relationship to the minor allele frequency of -0.5. For each 1Mb window that contains an identified genome-wide significant association, we then ask if the identified variants are correlated (squared correlation ≥ 0.01) to a causal variant: if so, we classify the window as a true positive; otherwise, we classify it as a false positive. The true positive rate is calculated as the number of true positive 1Mb windows divided by the total number of simulated causal variants, and it is also known as the recall, or sensitivity, reflecting the power of a statistical test. We provide two measures of the false discovery rate: FDR1 is calculated as the number of false positive 1Mb windows divided by the number of 1Mb windows that contain genome-wide significant associations, and it is a measure of the proportion of independent discoveries that are false. FDR2 is calculated as the total number of false positive SNPs (squared correlation ≤ 0.01 with a causal variant) identified divided by the total number of identified genome-wide significant associations. We calculate the type-I error rate (TIER) as the number of SNPs with no correlation to a causal variant (squared correlation ≤ 0.01 with a causal variant) that pass the Bonferroni genome-wide significance testing threshold divided by the total number of SNPs that have no correlation to a causal variant (squared correlation ≤ 0.01 with a causal variant).

We then further conduct another three simulation scenarios: *V4* where the 40,000 causal variants are drawn from the 2,174,071 SNP subset and we simulate a *α*-power relationship between the causal variant effect size and the minor allele frequency of *α* = −0.5; *V5* where the 40,000 causal variants are drawn from the 887,060 SNP subset and we simulate a *α*-power relationship between the causal variant effect size and the minor allele frequency of *α* = −0.5; and *V6* where the 5,000 causal variants are drawn from the 2,174,071 SNP subset with SNP-heritability = 0.25 and we simulate a *α*-power relationship between the causal variant effect size and the minor allele frequency of *α* = −0.5. Here, our objective in *V4, V5*, and *V6* is to compare REGENIE, LDAK-KVIK and gVAMP in the scenario where the causal variants are present in the data used to create the predictors for the first step of LOCO as to date simulations for these methods have assumed this. This ensures that our findings are not just driven by only having SNPs correlated with the causal variants in step 1. We present these results, comparing prediction accuracy of LDAK to gVAMP in Figure S17, comparing the chi-squared values at the causal variants from REGENIE, LDAK, and gVAMP in Figure S18, and comparing the TPR and FDR of REGENIE, LDAK, and gVAMP in Figure S19.

**Figure S20.**
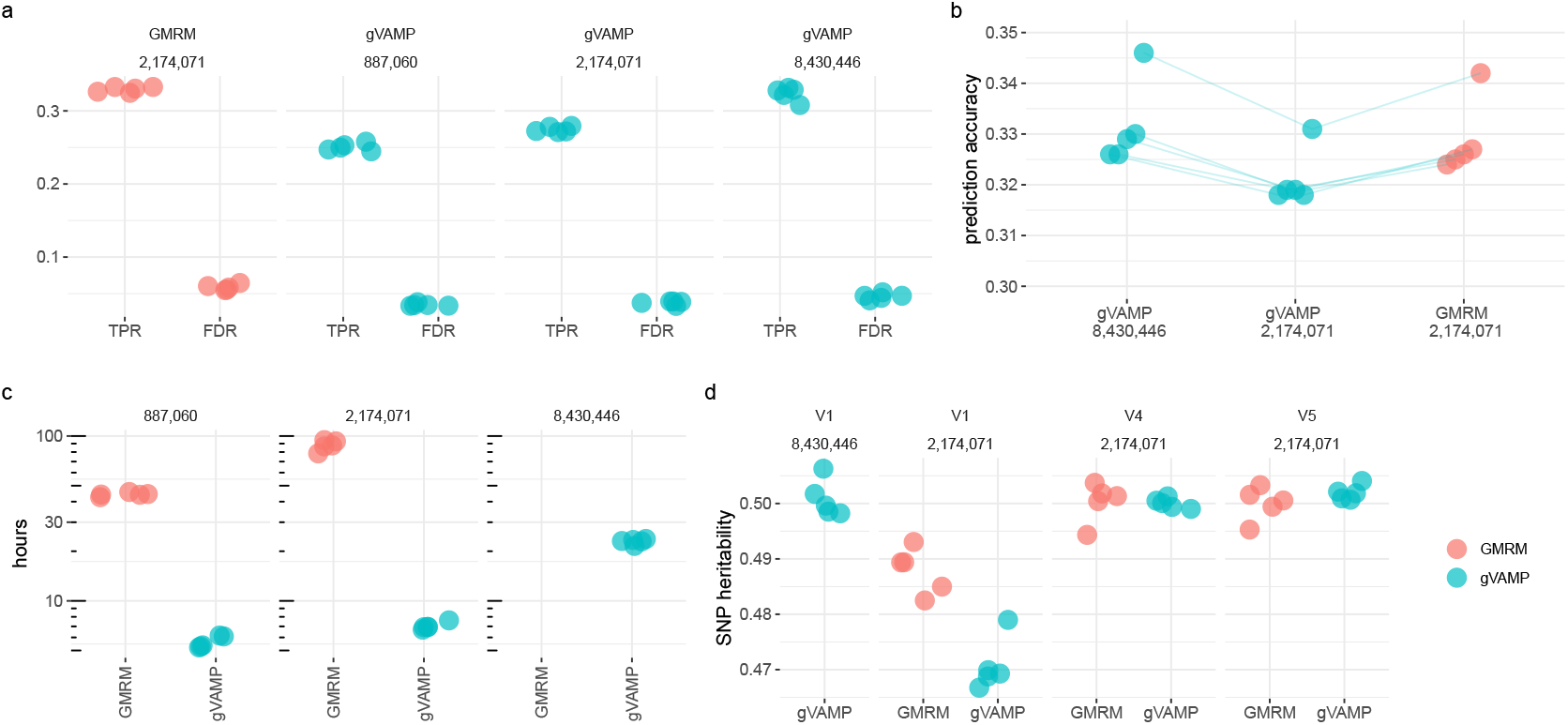
Comparison of MCMC Gibbs sampling algorithm GMRM to gVAMP in an imputed UK Biobank data simulation study. For a simulation setting where 40,000 causal variants are randomly selected from 8,430,446 SNP markers and effects are drawn from a Gaussian (“V1”), in *(a)*, we show results from standard leave-one-chromosome-out (LOCO) association testing where we select either all markers, 2,174,071 markers, or 887,060 markers for the first stage and then use all markers for the second stage LOCO testing. GMRM utilises 2,174,071 markers, whilst gVAMP can utilise the full range. We calculate the true positive rate (TPR) and the false discovery rate (FDR) and find that the FDR is well controlled at 5% or less for both gVAMP and GMRM. For *(b)*, we compare out-of-sample prediction accuracy for polygenic risk scores created at different sets of markers from gVAMP (8,430,446 and 2,174,071) and GMRM (2,174,071). *(c)* gives the run time in hours for the first stage analysis of gVAMP and GMRM, across different marker sets using 50 CPU from a single compute node. *(d)* gives a comparison of the proportion of phenotypic variation attributable to either 8,430,446 or a subset of 2,174,071 autosomal single nucleotide polymorphism (SNP) genetic markers (SNP heritability) estimated by GMRM and gVAMP. We consider three simulation scenarios: “V1” where 40,000 causal variants are randomly selected from 8,430,446 SNP markers and effects are drawn from a Gaussian, “V4” where 40,000 causal variants are randomly selected from 887,060 SNPs, with effects that contribute 50% to the phenotypic variance and have a power relationship to the minor allele frequency of -0.5, “V5” where 40,000 causal variants are randomly selected from 2,174,071 SNPs, with effects that contribute 50% to the phenotypic variance and have a power relationship to the minor allele frequency of -0.5. Points give the posterior means for GMRM and the convergence of gVAMP from five simulation replicates. Analysing 8,430,446 SNPs with gVAMP increases the heritability estimate over GMRM. This is consistent with an increase in phenotypic variance captured by the full imputed sequence data, as opposed to analyzing a selected subset of SNP markers, in which case gVAMP estimates are lower than those obtained from GMRM. Given the same data containing all the causal variants, the algorithms perform similarly irrespective of the underlying effect size distributions (“V4” and “V5”).

**Figure S21.**
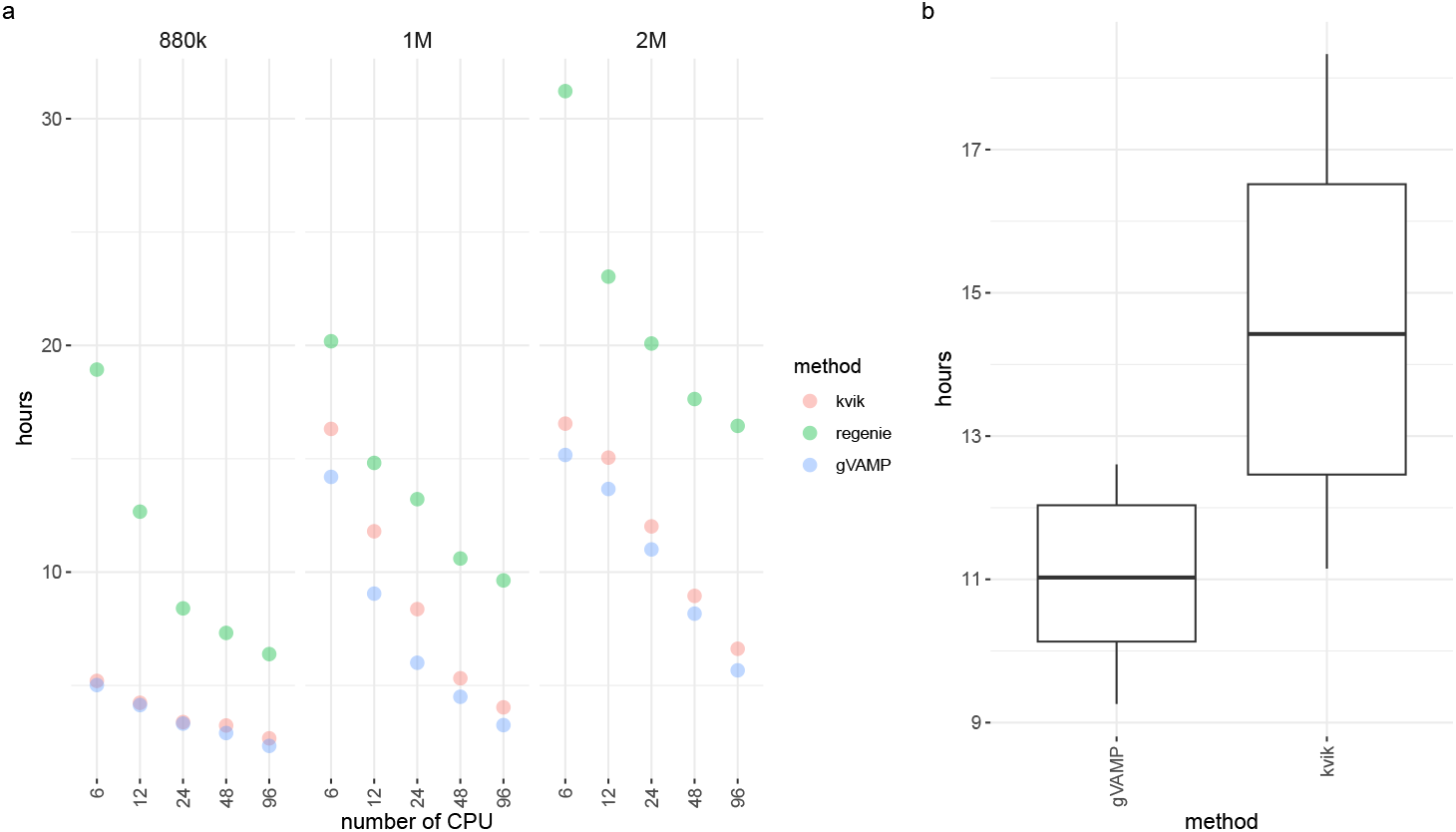
Timing comparison of LDAK elastic net prediction model and gVAMP in (a) an imputed UK Biobank data simulation study and (b) across 13 UK Biobank traits. *(a)* For simulation setting “V3” where 40,000 causal variants are randomly selected from 8,430,446 SNP markers, with effects drawn from an elastic net distribution, we show the total run time of the algorithms when run on different numbers of imputed SNP markers and given different numbers of CPU resources from a single AMD EPYC 7763 (64 cores, 2.45 GHz base-frequency) compute machine. *(b)* A boxplot of the run time of the algorithms when run on 13 different UK Biobank traits and 2,174,071 SNPs, when given 60 CPU. Training data sample size is given in Table S2 for each trait.

Additionally, we compare to GMRM using 2,174,071 SNP markers (as this completes within reasonable compute time and resource use), running for 2,500 iterations with 500 iteration burn in. Our objective is to compare GMRM and gVAMP to empirically assess the Bayes optimality of gVAMP when applied to genomic data. We compare the LOCO association testing, the out-of-sample prediction accuracy, the run time, and the SNP heritability of the two methods under the scenarios “V1”, “V4” and “V5”. We present these results in Figure S20. Finally, we compare the run time of REGENIE, LDAK and gVAMP both in a controlled simulation setting “V3” and across the 13 UK Biobank traits. We present these results in Figure S21.

### Imputed data simulation study results

For mixed linear model association testing (MLMA), association testing power (TPR) depends upon the sample size and the out-of-sample prediction accuracy of the predictors obtained from the first step [4]. Polygenic risk score prediction accuracy depends upon 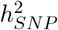, the number of true underlying causal variants and the sample size [5], which are fixed in our simulation allowing a comparison of prediction accuracy obtained by different approaches across different genetic architectures. We find that prediction accuracy increases as marker number increases for all approaches (Figure S13). The prediction accuracy obtained by gVAMP is always slightly higher than that obtained by the LDAK model in every scenario and this difference becomes larger as more markers are used (Figure S13).

When these predictors are used for leave-one-chromosome-out MLMA testing, we find that the average chi-square values of the causal variants increases as marker number increases only for gVAMP (Figure S14). The values obtained by gVAMP are always higher than those obtained by LDAK or REGENIE and this difference becomes larger as more markers are used (Figure S14). We find that gVAMP has better LOCO association testing performance as compared to REGENIE or LDAK in terms of the TPR and FDR (Figure S15). In the simulation settings presented here, REGENIE does not control the FDR unless larger numbers of markers are used in step 1 (Figure S15). LDAK controls the FDR and type-I error well, comparative to gVAMP, but has lower TPR (Figure S15). gVAMP with larger numbers of SNPs used for step 1 gives higher TPR, whilst controlling the FDR below 5% (Figure S15). Across all simulation settings, we find that the median chi-square value obtained from an MLMA-LOCO analysis with gVAMP is well controlled, showing the moderate elevation expected under polygenicity (Figure S16). This is in sharp contrast to REGENIE, which is elevated in all settings (Figure S16). The values obtained by LDAK rely on their proposal for adjusting the chi-square statistics and were generally lower than those obtained by MLMA-LOCO with gVAMP (Figure S16), even if we use the gVAMP LOCO predictors in place of the LDAK-KVIK ones and using LDAK-KVIK’s proposed step 2 (Table S3).

Similar patterns are also observed in the scenarios where the causal variants are present in the data used to create the predictors for the first step of LOCO, where gVAMP shows improved prediction accuracy (Figure S17), higher average-chi square values of the causal variants (Figure S18), higher TPR with controlled FDR (Figure S19).

Finally, we also show that gVAMP has comparative performance to a similar MCMC implementation of a Bayesian mixture of Gaussians variable selection regression model (Figure S20). Analysing 8,430,446 SNPs with gVAMP increases the SNP-heritability estimate over GMRM. This is consistent with an increase in phenotypic variance captured by the full imputed sequence data, as opposed to analyzing a selected subset of SNP markers, in which case gVAMP estimates are lower than those obtained from GMRM (Figure S20). Given the same data containing all the causal variants, the algorithms perform similarly irrespective of the scenario (Figure S20).

## Supplementary Note 2

### Participant inclusion in UK Biobank

UK Biobank has approval from the North-West Multicenter Research Ethics Committee (MREC) to obtain and disseminate data and samples from the participants (https://www.ukbiobank.ac.uk/ethics/), and these ethical regulations cover the work in this study. Written informed consent was obtained from all participants.

Our objective is to use the UK Biobank to provide proof of principle of our approach and to compare to state-of-the-art methods in applications to biobank data. We first restrict our analysis to a sample of European-ancestry UK Biobank individuals to provide a large sample size and more uniform genetic background with which to compare methods. To infer ancestry, we use both self-reported ethnic background (UK Biobank field 21000-0), selecting coding 1, and genetic ethnicity (UK Biobank field 22006-0), selecting coding 1. We project the 488,377 genotyped participants onto the first two genotypic principal components (PC) calculated from 2,504 individuals of the 1,000 Genomes project. Using the obtained PC loadings, we then assign each participant to the closest 1,000 Genomes project population, selecting individuals with PC1 projection ≤ absolute value 4 and PC2 projection ≤ absolute value 3. We apply this ancestry restriction as we wish to provide the first application of our approach, and to replicate our results, within a sample that is as genetically homogeneous as possible. Our approach can be applied within different human groups (by age, genetic sex, ethnicity, etc.). However, combining inference across different human groups requires a model that is capable of accounting for differences in minor allele frequency and linkage disequilibrium patterns across human populations. Here, the focus is to first demonstrate that our approach provides an optimal choice for biobank analyses, and ongoing work focuses on exploring differences in inference across a diverse range of human populations. Secondly, samples are also excluded based on UK Biobank quality control procedures with individuals removed of *(i)* extreme heterozygosity and missing genotype outliers; *(ii)* a genetically inferred gender that did not match the self-reported gender; *(iii)* putative sex chromosome aneuploidy; *(iv)* exclusion from kinship inference; *(v)* withdrawn consent.

### Whole genome sequence data

We process the population-level WGS variants, recently released on the UK Biobank DNAnexus platform. We use bcftools [6] and VCF Trimmer [7] to process thousands of pVCF files storing the chunks of DNA sequences, applying elementary filters on genotype quality (GQ ≤ 10), local allele depth (smpl_sum LAD < 8), missing genotype (F_MISSING > 0.1), and minor allele frequency (MAF < 0.0001). We select this MAF threshold as it means that on average about 80 people will have a genotype that is non-zero, which was the lowest frequency for which we felt that there was adequate power in the data to detect the variants and estimate the regression coefficients. Simultaneously, we normalize the indels to the most recent reference, removing redundant data fields to reduce the size of the files. Finally, we remove the variants sharing the same base pair position, not keeping any of these duplicates. For all chromosomes separately, we then concatenate all the pre-processed VCF files and convert them into PLINK format.

The compute nodes on the DNAnexus system are quite RAM limited, and it is not possible to analyze the WGS data outside of this system, which restricts the number of variants that can be analyzed jointly. To reduce the number of variants to the scale which can be fit in the largest computational instance available on the DNAnexus platform, we rank variants by minor allele frequency and remove the variants in high LD with the most common variants using the PLINK clumping approach, setting a 1000 kb radius, and *R*^2^ threshold to 0.36. This selects a focal common variant from a group of other common variants with correlation ≥ 0.6, which serves to capture the common variant signal into groups, whilst keeping all rare variation within the data. Finally, we merge all the chromosomes into a large data instance, giving the final 16,854,878 WGS variants.

### Imputed SNP data

We use genotype probabilities from version 3 of the imputed autosomal genotype data provided by the UK Biobank to hard-call the single nucleotide polymorphism (SNP) genotypes for variants with an imputation quality score above 0.3. The hard-call-threshold is 0.1, setting the genotypes with probability ≤ 0.9 as missing. From the good quality markers (with missingness less than 5% and *p*-value for the Hardy-Weinberg test larger than 10^−6^, as determined in the set of unrelated Europeans) we select those with rs identifier, in the set of European-ancestry participants, providing a dataset of 23,609,048 SNPs.

We further subset the 23,609,048 SNP data by selecting markers with MAF ≥ 0.002 to give 8,430,446 autosomal markers. We then select additional two subsets of these data by taking one marker from a set of variants with LD *R*^2^ ≥ 0.8 within a 1MB window to obtain 2,174,071 markers, and selecting with LD *R*^2^ ≥ 0.5 to obtain 882,727 SNP markers. This results in the selection of two subsets of “tagging variants”, with only variants in very high LD with the tag SNPs removed. This allows us to compare analysis methods that are restricted in the number of SNPs that can be analyzed, but still provide them a set of markers that are all correlated with the full set of imputed SNP variants, limiting the loss of association power by ensuring that the subset is correlated to those SNPs that are removed.

We recall that rare variants are directly observed in WGS data as opposed to being largely inferred within imputed data. Rare variants are also more prone to imputation errors and the subset of variants that can be inferred accurately must, by definition, have some correlation to the variants measured on the genotyping array platform [8]. Thus, while we tried to compare gVAMP on WGS and imputed data using same MAF threshold of 0.0001, we find that the imputed data analyses give estimates with small out-of-sample prediction accuracy from either gVAMP (6% correlation), ridge regression (4% correlation), or an elastic net (3% correlation). All of these models appear to fit the training data well, but they do not generalise well out-of-sample, and this is likely due to the highly correlated nature of the dataset creating difficulties in separating the effects of correlated rare and common variants. Thus, we can only make comparisons between WGS and imputed data by increasing the MAF threshold used to 0.002, creating the sets of 8,430,446 and 2,174,071 imputed SNPs.

### Whole exome sequence data burden scores

We then combine this data with the UK Biobank whole exome sequence data. The UK Biobank final release dataset of population level exome variant calls files is used (https://doi.org/10.1101/572347). Genomic data preparation and aggregation is conducted with custom pipeline (repo) on the UK Biobank Research Analysis Platform (RAP) with DXJupyterLab Spark Cluster App (v. 2.1.1). Only biallelic sites and high quality variants are retained according to the following criteria: individual and variant missingness < 10%, Hardy-Weinberg Equilibrium *p*-value > 10^−15^, minimum read coverage depth of 7, at least one sample per site passing the allele balance threshold > 0.15. Genomic variants in canonical, protein coding transcripts (Ensembl VERSION) are annotated with the Ensembl Variant Effect Predictor (VEP) tool (docker image ensemblorg/ensembl-vep:release_110.1). High-confidence (HC) loss-of-function (LoF) variants are identified with the LOFTEE plugin (v1.0.4_GRCh38). For each gene, homozygous or multiple heterozygous individuals for LoF variants have received a score of 2, those with a single heterozygous LoF variant 1, and the rest 0. We chose to use the WES data to create the burden scores rather than the WGS data as existing well-tested pipelines were available.

### Phenotypic records

Finally, we link these DNA data to the measurements, tests, and electronic health record data available in the UK Biobank [9] and, for the imputed SNP data, we select 7 blood based biomarkers and 6 quantitative measures which show ≥ 15% SNP heritability and ≥ 5% out-of-sample prediction accuracy [10]. Our focus is on selecting a group of phenotypes for which there is sufficient power to observe differences among approaches. We split the sample into training and testing sets for each phenotype, selecting 15,000 individuals that are unrelated (SNP marker relatedness < 0.05) to the training individuals to use as a testing set. This provides an independent sample of data with which to access prediction accuracy. We restrict our prediction analyses to this subset as predicting across other biobank data introduces issues of phenotypic concordance, minor allele frequency and linkage disequilibrium differences. In fact, our objective is to simply benchmark methods on as uniform a dataset as we can. As stated, combining inference across different human groups, requires a model that is capable of accounting for differences in minor allele frequency and linkage disequilibrium patterns across human populations and, while our algorithmic framework can provide the basis of new methods for this problem, the focus here is on benchmarking in the simpler linear model setting. Samples sizes and traits used in our analyses are given in Table S2. Prior to analysis all phenotypic values were adjusted by the participant’s recruitment age, their genetic sex, north-south and east-west location coordinates, measurement center, genotype batch, and the leading 20 genetic PCs.

## Supplementary Note 3

### Historical development and informal description of the AMP framework for Bayesian regression

Approximate message passing (AMP) was originally proposed for linear regression [11–13] assuming a Gaussian design matrix ***X***. To accommodate a wider class of structured design matrices, vector approximate message passing (VAMP) was introduced in [14]. The performance of VAMP can be precisely characterized via a deterministic, low-dimensional *state evolution* recursion, for any right-orthogonally invariant design matrix. We recall that a matrix is right-orthogonally invariant if its right singular vectors are distributed according to the Haar measure, i.e., they are uniform in the group of orthogonal matrices. Empirical evidence [15, 16] suggests that the predictions for rotationally invariant models hold for a wide class of scenarios, and a line of work has aimed at showing that this is case for AMP algorithms [17, 18].

In the rest of this subsection, we provide an informal overview of the Approximate Message Passing (AMP) framework applied to the Bayesian linear regression problem.

We begin with the well-known LASSO optimization problem. For a given regularization parameter *λ* ∈ ℝ_+_, the estimator of the effect sizes 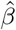 is defined as

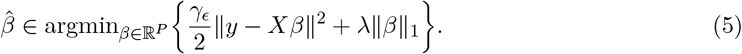

This formulation admits a Bayesian interpretation: it can be seen as a posterior mode estimation problem for a linear regression model with independent Laplace(0, 1*/λ*) priors on the coefficients.

A classical algorithm for solving (5) is the iterative soft-thresholding algorithm (ISTA) first introduced in [19], which initializes the estimate 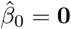 and then reads as follows:

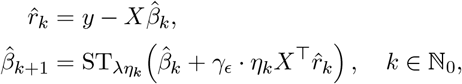

where 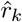 is the residual at iteration *k, η*_*k*_ > 0 is a deterministic step size, and ST_*t*_ : ℝ → ℝ denotes the *soft-thresholding operator* defined by

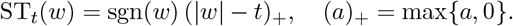

Beck and Teboulle (2009) in [20] proposed its extension, the *fast iterative shrinkage-thresholding algorithm (FISTA)*, which is closely related to AMP and is guaranteed to converge to the solution of (5). The iteration takes the form

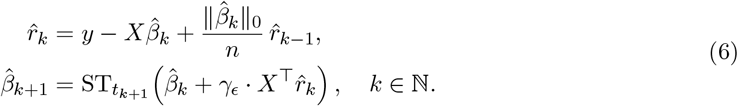

where *t*_*k*+1_ > 0 is a deterministic threshold and 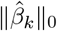 denotes the number of nonzero entries in 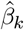. In contrast to ISTA, the residual 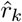 here includes a *memory correction term*, which is crucial for enabling exact characterization of the empirical distribution of the iterates.

Returning to the LASSO formulation in (5), one may wish to replace the Laplace prior with a more flexible alternative, such as a *spike-and-slab prior*, where the slab component is a Gaussian mixture. In the AMP framework, this modification can be achieved simply by replacing the soft-thresholding operator in (6) with the conditional expectation under the spike-and-slab prior, and adjusting the residual correction term accordingly.

This yields an AMP-based framework for Bayesian regression under spike-and-slab priors, where the prior parameters can be updated iteratively via the expectation–maximization (EM) algorithm. However, classical AMP is known to work reliably only when the design matrix **X** has i.i.d. sub-Gaussian entries. To address broader classes of matrices, including those with correlated entries, the *vector AMP (VAMP)* algorithm was developed [14], which extends the applicability of AMP well beyond the i.i.d. setting.

## Supplementary Note 4

### Explicit expressions for denoisers and the Onsager correction

From Bayes theorem, one can calculate the posterior distribution (which here has the form of a spike-and-slab mixture of Gaussians) and obtain its expectation. Hence, by denoting a generic component of ***r***_1,*t*_ as *r*_1_, it follows that

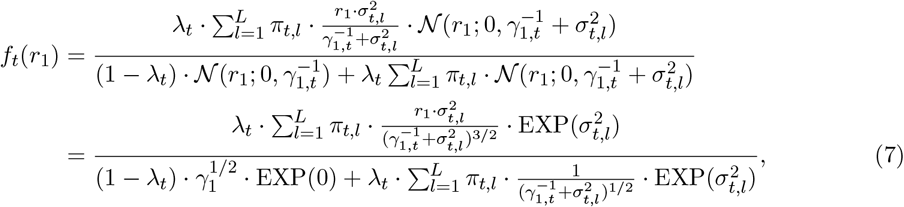

where 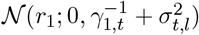 denotes the probability density function of a Gaussian with mean 0 and variance 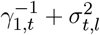 evaluated at *r*_1_. Furthermore, we set

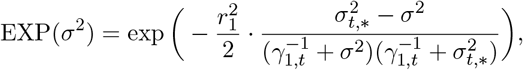

with 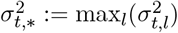. This form of the denoiser is particularly convenient, as we typically deal with very sparse distributions when estimating genetic associations. We also note that the calculation of the Onsager coefficient in line 17 of Algorithm 1 requires the evaluation of a conditional variance, which is computed as the ratio of the derivative of the denoiser over the error in the estimation of the signal, i.e.,

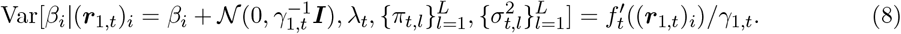

The calculation of the derivative of *f*_*t*_ is detailed in the following section.

### Onsager correction calculation

In order to ensure Gaussianity of residuals, gVAMP calculates the so-called Onsager correction based on (8). For such calculation, the derivative of the denoising function *f*_*t*_ defined in (3) is required. Let us denote the numerator and denominator of (7) with Num(*r*_1_) and Den(*r*_1_), respectively. Then,

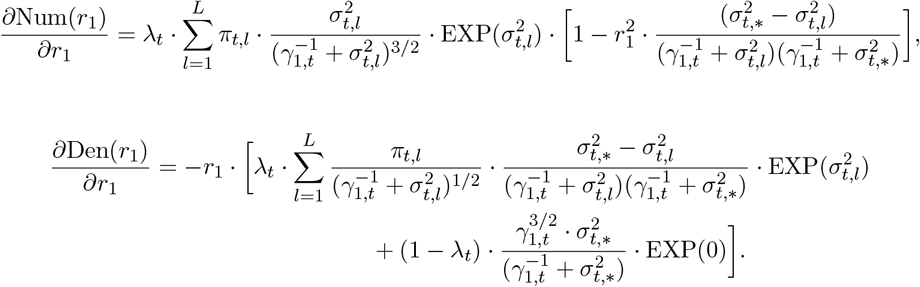

Thus, the Onsager correction reads

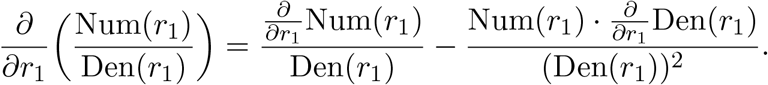

## Supplementary Note 5

### Conjugate gradient algorithm for solving linear systems

#### Algorithm 2

Conjugate gradient method for solving a symmetric linear system *Ax* = *b*.

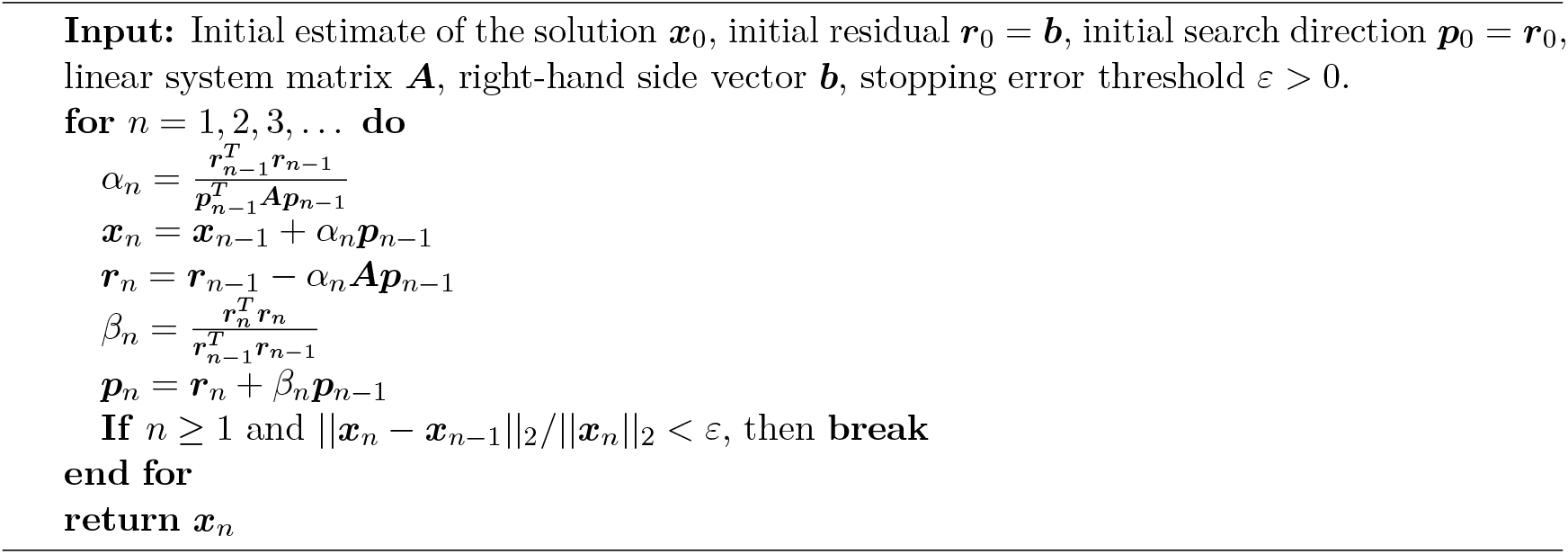

## Supplementary Note 6

### Estimation of the prior and noise precision via EM

In this section we give precise formulas for expectation-maximization updates of hyperparameters related to prior distribution of effect sizes and noise precision:

- Sparsity rate *λ*: We define 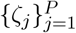 as:

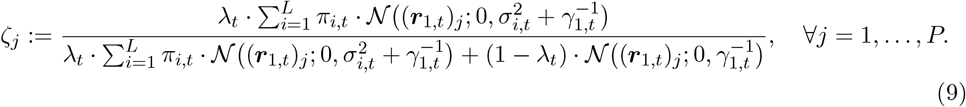

The intuition behind 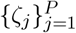 is that each *ζ*_*j*_ tells what fraction of posterior probability mass was assigned to the event that it has a non-zero effect. Then, the update formula for the sparsity rate *λ*_*t*+1_ reads as

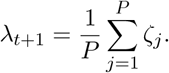

- Probabilities of mixture components in the slab part 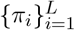: We define 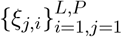 as

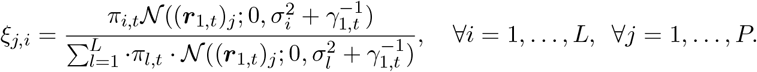

The intuition behind 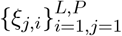 is that each *ξ*_*j,i*_ approximates the posterior probability that a marker *j* belongs to a mixture *i* conditional on the fact that it has non-zero effect. Thus, the update formula for *π*_*i,t*+1_ reads as

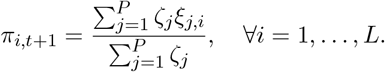

- Variances of mixture components in the slab part 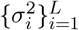: Using the same notation, the update formula reads as

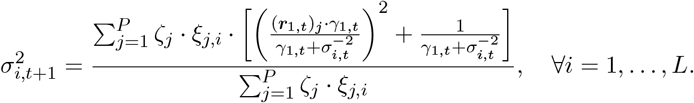

Here we also introduce a mixture merging step, i.e., if the two mixtures are represented by variances that are close to each other in relative terms, then we merge those mixtures together. Thus, we adaptively learn the mixture number.

- Precision of the error *γ*_*ϵ*_ : We define **Σ**_*t*_ := (*γ*_*ϵ,t*_***X***^*T*^ ***X*** + *γ*_2,*t*_***I***)^−1^ . Then, the update formula for the estimator of *γ*_*ϵ*_ reads as

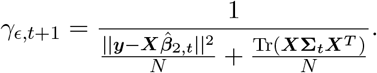

In the formula above, the term 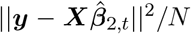 takes into account the quality of the fit of the model, while the term Tr(***X*Σ**_*t*_***X***^*T*^)*/N* prevents overfitting by accounting for the structure of the prior distribution of the effect sizes via the regularization term *γ*_2,*t*_. We note that the naive evaluation of this term would require an inversion of a matrix of size *P* × *P* . We again use the Hutchinson estimator for the trace to approximate this object, i.e., Tr(***X*Σ**_*t*_***X***^*T*^) = Tr(***X***^*T*^ ***X*Σ**_*t*_) ≈ ***u***^*T*^ (***X***^*T*^ ***X*Σ**_*t*_)***u***, where ***u*** has i.i.d. entries equal to −1 and +1 with the same probability. Furthermore, instead of solving a linear system **Σ**_*t*_***u*** with a newly generated ***u***, we re-use the ***u*** sampled when constructing the Onsager coefficient, thus saving the time needed to construct the object **Σ**_*t*_***u***.

## Notes

https://github.com/medical-genomics-group/gVAMP

### Competing Interest Statement

MRR receives funding from Boehringer Ingeleheim for a collaborative research project that is unrelated to this study.

### Summary of Updates

Very minor updates to grant acknowledgements.

